# Structural basis of directionality control in large serine integrases

**DOI:** 10.1101/2025.01.03.631226

**Authors:** Heewhan Shin, Ying Pigli, Tania Peña Reyes, Adebayo J. Bello, James R. Fuller, Femi J. Olorunniji, Phoebe A. Rice

**Affiliations:** Department of Biochemistry & Molecular Biology, The University of Chicago; Chicago IL, 60637, USA; School of Pharmacy and Biomolecular Sciences, Liverpool John Moores University; Liverpool, L3 3AF, UK

## Abstract

Large serine integrases (LSIs) catalyze unidirectional site-specific insertion of large DNA payloads, and in the presence of a cognate recombination directionality factor (RDF), catalyze unidirectional excision. Because neither reaction changes the net number of covalent bonds, the preferred direction must be controlled by the energetics of the changing protein-DNA complexes along these reaction pathways. However, a detailed understanding has been hampered by a lack of structural information. Here, we report 8 structures of SPβ integrase-DNA complexes along the integrative (-RDF) and excisive (+RDF) reaction pathways, at resolutions extending to 3.15 Å. These complexes include tetrameric intermediates before and after strand exchange and product-bound dimers for both pathways. Our findings reveal that both recombination-induced conformational changes and RDF-mediated repositioning of the integrase’s coiled-coil subdomain (1) dictate which pairs of DNA sites can be assembled into a synaptic complex to initiate recombination and (2) dictate which product complexes will be conformationally locked, preventing back reactions. Critically, we find that the synaptic complex in which excision occurs is fundamentally different from that in which integration occurs. These mechanistic insights provide a conceptual framework for engineering efficient and versatile genome editing tools.

## Introduction

Large serine integrases (LSIs; also called large serine recombinases) and their cognate recombination directionality factors (RDFs) mediate site-specific DNA recombination reactions that move the phage or mobile genetic elements that encodes them into and out of host bacterial genomes^1^. Despite extensive biochemical and protein engineering studies, structures have only been available for fragments of these enzymes, leaving many unanswered questions regarding how they recognize different attachment (*att*) sites, how their conformations and protein-protein contacts shift along the reaction pathway for strand exchange, and most importantly, how directionality is dictated^2–7^. The work presented here elucidates a unique and elegant system for control of recombination directionality of both the integration and excision reactions.

Several features of these systems make them useful as genetic tools^8,9^. For integrative recombination, only integrase and two short DNA sites are required: an ∼50-54 bp phage DNA site (*attP*) and an ∼40-46 bp bacterial site (*attB*)^10–12^. The products of this unidirectional reaction are two hybrid integrase binding sites flanking the integrated phage DNA, *attL* and *attR* (Figure 1A). Binding of the RDF to the integrase triggers the reverse (excision) reaction and inhibits the integration reaction^11–16^. Unlike insertions mediated by DDE-family transposases or CRISPR-Cas, no DNA repair is needed after LSI-mediated reactions because all broken phosphodiester bonds are religated (Figure 1B). The large number of known integrases (and smaller number of known integrase-RDF pairs) provides an array of sequence specificity options for mechanistic studies and synthetic biology applications. Finally, new tools for inserting *att* sites into heterologous genomes have streamlined the use of LSIs in genome editing^17–24^.

**Figure 1.**
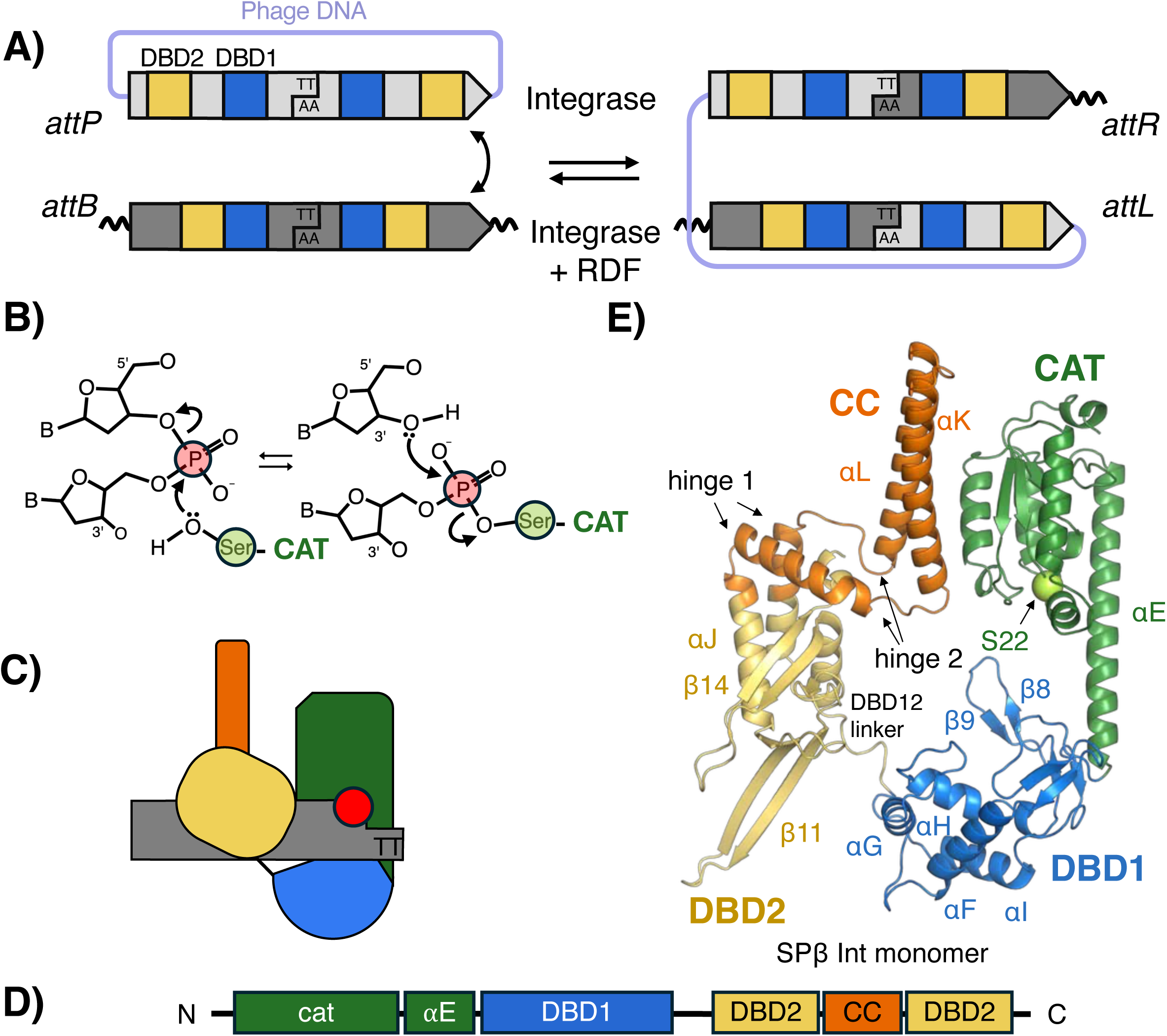
Large serine integrase – mediated DNA recombination. A. Schematic diagram of the reaction. The integrase recombines a specific site within the phage (*attP*; light gray) with one found in the host chromosome (*attB*; dark gray). DNA cleavage in the center of the site creates a 2 nt 3’ overhang. After exchange of the right-hand halves of the sites (as shown), the prophage is inserted within the host chromosome, flanked by hybrid sites termed *attL* and *attR*. Motifs recognized by the first DNA binding domain of the Integrase (DBD1; also known as a recombinase domain or RD) are shown in blue and motifs recognized by DBD2 (also known as zinc-binding domain or ZD) are shown in yellow. Note that *attP* and *attB* sites differ in the spacing of these two motifs. The RDF protein triggers the integrase to catalyze the reverse reaction. B. The catalytic domain (“cat”; green) is covalently linked to the 5’ ends of the cleaved DNA intermediate through a phosphoserine linkage. The phosphorous atom is indicated by a red circle and Cα of Ser 22 is indicated by a yellow green circle here and throughout the figures. C. Schematic representation of a large serine integrase bound along an *attB* half site D. Domain architecture of large serine integrase which consists of catalytic domain (green), DBD1 (blue), DBD2 (yellow); and a coiled coil motif (CC; orange) that is inserted within DBD2. E. Ribbon diagram of one SPβ Int monomer isolated from the synaptic complex.

Serine recombinases share overlapping biological roles with tyrosine recombinases, but their sequences, structures, and mechanisms for DNA strand exchange and directionality control are unrelated. Some tyrosine recombinases such as Cre and Flp require only simple DNA sites but catalyze bidirectional reactions. Other tyrosine recombinases such as phage lambda integrase, which was the subject of much historic work, do catalyze unidirectional insertion, but require a long *attP* site (240 bp for lambda), as well as host proteins^25^. Furthermore, their recombination directionality factors (xis) bind DNA as well as the integrase protein (in contrast to the RDFs for LSIs, which only bind the integrase protein)^25^. Nevertheless, we show here that serine integrases share a fundamental concept with tyrosine integrases: the use of structurally different synaptic complexes for integrative and excisive recombination.

LSIs share a conserved catalytic domain and strand exchange mechanism with smaller recombinases from the serine family such as resolvases and invertases^1,26–28^. Within a synaptic complex, which contains a recombinase tetramer and two partner DNAs, a key catalytic serine residue from each subunit attacks a DNA phosphate, displacing a 3’ OH and generating double-strand breaks with 2-nt 3’ overhangs. In this intermediate state, each 5’ end is covalently linked to a protein subunit (Figure 1B,C, and S1), and the two halves of the synaptic complex can swivel relative to one another about a central flat, hydrophobic interface to reposition the broken ends^29^. Following a ∼180° rotation, the chemical reverse of the cleavage reaction can occur, which religates the DNA in the recombinant configuration^29–31^. However, if the 2-nt overhangs (the “central dinucleotides”) do not base pair after 180° rotation, the complex can continue swiveling and eventually religate substrate DNAs^32–35^.

A key control point for serine recombinase regulation is the transition from inactive dimeric protein-DNA complexes to active tetrameric synaptic complexes. Interactions among catalytic domains are central to both oligomeric forms. However, formation of a tetramer entails more than just rigid-body docking of two DNA-bound dimers – it also entails large conformational changes that license catalytic activity and create the central flat interface about which rotation occurs. In the absence of additional tetramer-stabilizing factors, the dimer conformation is strongly favored^36–38^. The additional factors that control this conformational equilibrium vary widely among different subfamilies of serine recombinases^1,27,39,40^. For small serine recombinases that act as DNA resolvases and invertases, formation of a catalytically active tetramer is promoted by additional DNA-binding proteins that form a topologically defined complex^41–45^. As described below, for large serine integrases stabilization of substrate-bound tetramers is promoted by an additional domain of the integrase protein itself.

The forward progression of serine recombinase-catalyzed reactions is not driven by changes in chemical bond energy but rather by other energetic differences between the substrates vs. products^26^. For small serine resolvases, an overall negative ΔG is provided by the accompanying change in DNA topology^26,46^. The unidirectionality of LSI-catalyzed reactions does not rely on DNA topology and therefore must depend on differences between the substrate and product protein-DNA complexes. This is feasible in a thermodynamic framework because, unlike true catalysts that cannot change chemical equilibria, serine recombinases tend to remain bound to their products and therefore should be considered as components of the reaction itself. However, a lack of 3D structural information has hampered a detailed understanding of how LSIs apply this principle to accomplish directionality in general, as well as how RDFs reverse it.

The modular domain structure of LSIs facilitates stabilization of substrate-bound tetramers and control of reaction direction. All serine recombinases share a catalytic domain (“CAT”), followed by a long helix (“helix E”) that is sometimes considered to be part of the catalytic domain. Small serine recombinases carry a single DNA binding domain, usually at the C-terminus, but at the N-terminus for certain family members that act as transposases^47^. LSIs contain two DNA-binding domains that follow helix E, DBD1 and DBD2, also referred to as a recombinase domain (RD) and a zinc ribbon domain (ZD), respectively (Figure 1C and 1D)^5,10^. Inserted within DBD2, and flanked by flexible hinges, is an antiparallel coiled coil (CC) with a hydrophobic tip. As described below, interactions between pairs of CC tips can mediate either activating inter-dimer interactions or inhibitory intra-dimer interactions, depending on the context - that is, depending on which DNA sites the proteins are bound to, and the presence or absence of the RDF^2,5,48–52^.

Differentiating substrates from products requires differences in the partner DNA sites that alter how the protein interacts with them. For LSIs, *attP* and *attB* (the substrates for integrative recombination) are functionally symmetric (except for their central dinucleotides) but differ from one another in length and sequence (Figure S2). The critical difference between *attP* and *attB* sites was deduced from the structure of the C-terminal portion of phage LI integrase bound to half of an *attP* site^48^. The authors noted that the motifs recognized by DBD2 are 5 bp, or half a helical turn, closer to the center of *attB* sites than of *attP* sites^5^. Furthermore, they proposed that this positioning of DBD2 allows the CC within it to (1) promote synapsis of *attP* and *attB* sites via inter-dimer “handshakes” between CC’s and (2) after subunit rotation, swap interaction partners to lock the dimer-bound *attL* and *attR* products via intra-dimer “handcuffing”.

RDF binding was proposed to unlock the *attL*- and *attR*-bound dimers, allowing re-formation of an active synaptic complex^5,51^. In agreement with this idea, despite high variability among known RDFs in sequence and predicted structure, they share a common predicted binding site near the base of the CC-motif of their cognate LSIs^53–55^. However, this model for RDF function lacks a structural underpinning and struggles to explain the unidirectionality of the excisive recombination reaction - that is, it does not explain why the RDF not only activates *attL* x *attR* recombination but also inhibits *attP* x *attB* recombination.

To understand the structural basis of directionality control, and the role of the RDF in the process, we determined 8 cryo-EM structures of *B. subtilis* phage SPβ integrase (SprA) in the presence and absence of its cognate RDF, SprB^56^ and with different *att* sites, at resolutions ranging from 3.15-7.18Å. These structures include synaptic tetramers for both the integrative and excisive pathways, as well as product dimers for both pathways. These are the first complete LSI-DNA complex structures and the first experimental views of LSI-RDF interactions. The synaptic tetramers were captured in the covalent intermediate state, when the complex was free to swivel, and for each pathway, both the pre- and post- strand exchange states were populated.

Our results provide direct evidence to support and expand upon prior models for directionality control in LSI-mediated integrative recombination and provide a new model for unidirectionality of the excision reaction^2,49,52,57^. Our structures show that product dimers in both pathways, not just the integrative one, are handcuffed by CC-CC interactions. Furthermore, we show that the prevailing model for RDF function is incorrect in that rather than simply unlocking *attL*- and *attR*-bound dimers, RDF binding redirects the CC to dictate a very different synaptic complex for the excisive vs. the integrative reaction.

## Results

### Eight structures illuminate integrative and excisive recombination

To obtain structures along the integrative pathway, we used wild type (WT) SPβ integrase. For excisive pathway structures, we used the previously-described method of fusing the RDF to the C-terminus of the integrase with a flexible linker between the two^58^, which prevented accidental loss of the small RDF protein. The DNA duplexes used carried the WT *att* sequences, except for the central dinucleotides, as described below. For most structures reported here, an additional dA_8_ was added to the ends of *attP*-derived half sites to facilitate distinguishing them from *attB*-derived half sites. This “tag” proved unnecessary but was visible in the maps even though not contacted by the protein, presumably due to the unusually rigid nature of A tracts^59,60^. Note that in all our structures, the DNA strands are numbered such that the bases of the central dinucleotide are -1 and +1.

Product-bound dimer complexes containing uncleaved *attL or attP* DNA were determined using sites in which the central dinucleotide was symmetrized (AT rather than AA, denoted as *attLsym* and *attPsym*). This was unnecessary for cryoEM but facilitated accompanying biochemical experiments: either relative orientation of *attPsym* and *attBsym* can be recombined by the integrase, while two copies of *attLsym* can be recombined by the integrase-RDF fusion. Previous data showed that the only functional difference between WT *attL* and *attR* sites is their asymmetric central dinucleotide (Figure S3)^61^.

We also trapped synaptic complexes in the double-strand-break intermediate state in which Ser 22 of each subunit is covalently attached to a 5’ DNA end. To do so, we changed the central dinucleotide of both strands to TT (we termed these mismatched sites *attPmm, attBmm* and *attLmm*; Figure S1). These sites were designed to trap the covalent intermediate based on observations that serine recombinases can cleave substrates with central mismatches but cannot religate them^32^. These samples yielded structures of the pre- and post-rotation states for both the integrative and the excisive synaptic complexes, as well as two unplanned additional structures, as described below.

### The *attL*-bound product dimer: a 4bp shift between half sites allows CC-CC handcuffing

We focus first on the *attL*-bound integrase dimer structure because that map extended to the highest resolution (3.15 Å) (Figure S5). This dimer provides a model for both products of integrative (*attB* x *attP*) recombination, because the *attL*- and *attR*-bound dimers each contain one *attB*- and one *attP*-derived half site (Figure 2A).

**Figure 2.**
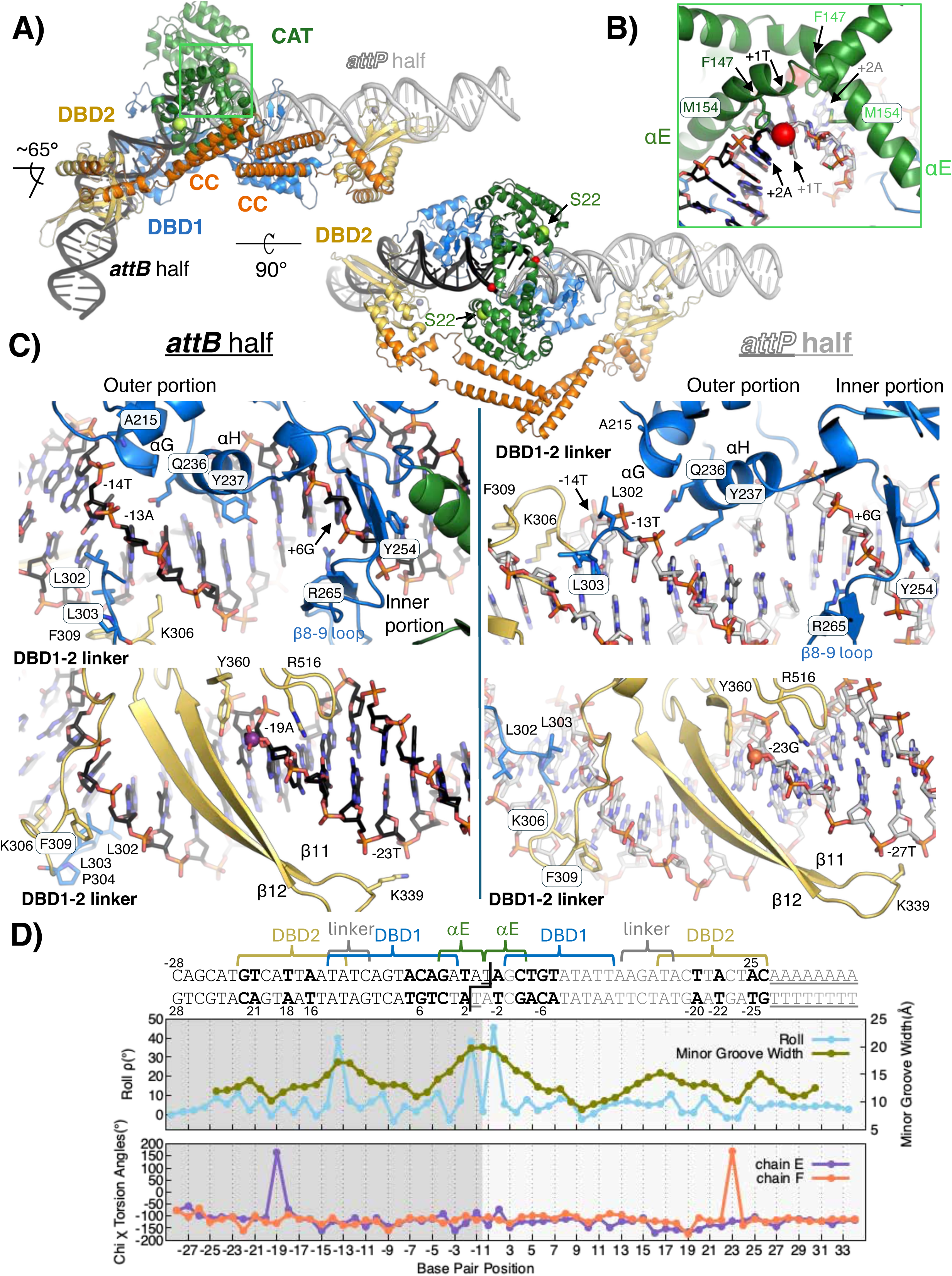
Distinct DNA attachment site binding: *attLsym* bound SPβ Integrase dimer. A. Cryo-EM structure of the SPβ Integrase dimeric complex with *attLsym* shown in side (left) and top (right) views. The coloring scheme corresponds to Figure 1D, and red spheres mark the scissile phosphates and yellow green spheres mark the catalytic S22. B. A close-up view near the central dinucleotide (boxed region in panel A, side view). This panel shows the bent E-helices in both subunits and the intercalation of residues F147 and M154 between +1T and +2A (cyan; color scheme corresponds to panel D) C. Structural comparison of DBD1 (top row) and DBD2 (bottom row) binding to *attB*-derived (left column) and *attP*-derived (right column) half sites. The outer portion of DBD1 (left side in both panel) and the DBD1-2 linker interact differently with the two types of half site. Binding of the DBD2 domains induces an unusual DNA kink at nucleotides -19/-23, indicated by spheres in colors corresponding to panel D). D. Sequence and structural parameters of *attLsym*. Underlined nucleotides differ from the WT *attL* sequence and bold nucleotides are identical in all four half-sites. Interacting protein domains are noted above the sequence, and structural parameters are plotted below (roll angle between successive base pairs, minor groove widths and chi χ torsion angles). The DBD2 domains exhibit a 4-bp shift between *att* sites.

The primary difference between the *attB* and *attP* sides of the complex is that DBD2 lies 4bp closer to the center on the *attB* half-sites. The overall architecture of the *attP*-side of the complex resembles that of the LI-*attP* half site complex structure^48^. However, although analysis of many LSI binding site sequences predicts a 5bp shift of DBD2 between *attP* and *attB* sites^3,5,51,54^, the density map was clear in this region (Figure S6), and the outer portions of the SPβ binding site sequences align better if a 4bp shift is assumed (Figures S2 and S6). A consequence of this 4-5 bp shift of nearly half a helical turn is that DBD2 of the *attP*-bound subunit docks onto same side of the double helix as DBD1, but for the *attB*-bound subunit, the opposite side. Furthermore, the DBD2s of the two different subunits bound to *attL* are positioned on the same face of the complex.

The *attL*-bound dimer structure visualizes the details of the product-stabilizing intra-dimer CC-CC interaction (Figure 2A and S6A) first proposed by Rutherford and Van Duyne^5,51^. This interaction is facilitated by the relative placement of the two DBD2s, which the CCs are inserted into. The DBD2-CC junctions are flexible, but the CCs themselves are well-ordered. As seen in structures of the isolated CCs of LI integrase, the two tips interact in an asymmetric front-to-back manner, in which the front of the *attP*-bound subunit’s CC interacts with the back of the *attB*-bound subunit’s CC, rather than twofold-symmetric manner (Figure S7A)^2^. The *attB*-bound subunit’s CC is sandwiched between that subunit’s catalytic domain and the *attP*-bound subunit’s CC (Figure 2A, S6A, and S7B). That one of the CCs would interact with a catalytic domain was unexpected.

The *attL*-bound dimer is clearly in an inactive state because the two active site serines (S22) are 9-10 Å from the phosphorus atoms that they would attack in a DNA cleavage reaction (Figure 2A and S6B). The E-helices are bent and kinked and interact with one another in a largely antiparallel fashion that differs from the more nearly parallel interaction seen in DNA-bound dimers of small serine recombinases^36,43,44^. This unusual catalytic domain dimer structure is discussed in more detail below.

The DNA in this complex is bent at the center by ∼65°, reflecting two kinks flanking the central dinucleotide with large roll angles between adjacent base pairs (Figure 2A and D). The C-terminal portion of helix E, which connects the catalytic domain to DBD1, docks into the minor groove, widens it, and inserts the hydrophobic side chains of F147 and M154 into the kink (Figure 2B, D and S6B). Similar groove-widening by helix E, sometimes including hydrophobic intercalation, was reported for small serine recombinases^43,44^. These interactions also likely contribute to indirect readout of the DNA sequence near the cleavage sites, as they do in LI integrase^57^. These helix E-DNA interactions are similar on the *attB*- and *attP* sides of the *attL*-bound dimer, but are not maintained in the complexes with cleaved DNA reported below.

### Partially modular protein-DNA interactions

Four different protein segments interact closely with the DNA: the C-terminal portion of helix E, DBD1, the linker between DBDs, and DBD2 (Figure 2 and S6 and supplemental movie LSI_att_sites_recognition.mp4). Most of the integrase-DNA interactions depend only on whether a subunit is bound to an *attP*- vs. an *attB*-derived half site, rather than on the larger context of the complex (e.g. cleaved vs. uncleaved, +/- RDF, or pre- vs. post-rotation). The sole exception is the Helix E, which interacts differently with cleaved vs. uncleaved DNA. Details of the integrase – DNA interactions are therefore only shown for the highest-resolution one, the *attL*-bound dimer (Figure S6). Comparing the *attB*- and *attP*-bound subunits shows that integrase-DNA interactions are only partially modular: the local interactions of helix E, the inner (closer to the center) portion of DBD1, and DBD2 are similar on *attB* and *attP* sites, whereas the outer portion of DBD1 and the interdomain linker interact differently with *attB* vs. *attP* DNA.

The inner portion of DBD1 binds base pairs 4-7 of both types of half site. The identity of these bases is conserved among all four half-sites. Of particular interest here is the long β-hairpin loop between αH and αI that makes multiple contacts with the phosphodiester backbone, inserts Y254 into the minor groove and inserts R265 into the major groove where it makes a sequence-specific bidentate interaction with the conserved G6. The flat guanidinium group of R265 is positioned to stack against T5 with its protruding methyl, forming an interaction similar to that seen in the TG- and 5-methyl-CG specific interactions of other proteins^62^. The length of this loop varies widely among LSIs, and its major groove-interacting tip is absent in LI integrase, where DBD1 lacks significant sequence specificity^57^.

The outer portion of DBD1 interacts differently with the two types of half site. It contains a helix-turn-helix motif (αF, αG and αH). Even though such motifs often display strong preferences for specific DNA sequences, the bases that this one contacts are not conserved among all four half-sites. Helix H docks into the major groove, but only Q236 and Y237 extend toward the unique edges of the bases themselves. The tip of Q236 is within H-bonding distance of a base on the *attB* side but not on the *attP* side, and the side chain of Y237 adopts different rotamers and contacts different bases (Figure 2C). On the *attB* site, a kink between base pairs 13 and 14 facilitates H-bonding between a backbone amide of A215 from the N-terminus of αG and phosphate -14 (Figure 2C). On the *attP* side, the major groove is wider at this point, the minor groove narrower, and the phosphodiester backbone is pulled away from αG. This observation correlates with a more AT-rich sequence in *attP* than *attB* (Figure 2D): AT-rich sequences intrinsically favor a narrowed minor groove^63^.

The DBD1-DBD2 linker also interacts differently with the two types of half site. On the *attB* side of the complex, L302 and L303 of the linker partially intercalate between base pairs 13 and 14, inducing and/or stabilizing a nearly 40° roll angle that greatly widens the minor groove (Figure 2C, D and S6). The linker also interacts with a widened minor groove on the *attP* side, but the DNA is more smoothly bent, and by only ∼ 30°. Furthermore, rather than L302 and L303, residues P304, K306 and F309 interact most closely with the floor of the minor groove. On the shorter *attB* side, the linker-binding region overlaps with the outer portion of the DBD1 binding site (Figure 2D).

The local interactions between DBD2 and DNA are similar on both types of *att* site despite the overall 4bp shift (Figure 2C and S6). Direct H-bonding interactions between protein side chains and DNA bases are sparse, implying that sequence recognition may rely heavily on indirect readout of sequence-dependent variations in the structure and flexibility of the DNA. The most prominent DNA-binding feature of DBD2 is a long beta hairpin that lies in the major groove and makes multiple contacts with the phosphodiester backbone, ending with K339 contacting the 5’ phosphate of T -23 (*attB* side) / T -27 (attP side). This hairpin is unusually long in SPβ integrase (Figure S8), and therefore its cognate *attB* and *attP* sites are longer than those for many LSIs: 46 and 54 bp, respectively. A second beta hairpin places R516 in the minor groove, where it donates H-bonds to two consecutive bases. These contacts by themselves are not sequence specific because A, T, C and G all have an H-bond acceptor in a similar location. However, together with the other beta hairpin and Y360, they form part of a grip across the DNA backbone that induces an unusual kink centered on A-19 (*attB* side) / G-23 (*attP* side). Integrase activity depends not only on DNA affinity but on distinguishing *attP* from *attB*^64^, which in turn depends on proper placement of DBD2. Figure S6 shows that placement of DBD2 probably depends not only on the presence of conserved sequence motifs in the correct place, but also on their absence from the wrong place.

### The integrative synaptic complex trapped in the swiveling state

Integrative synaptic complexes were formed by mixing SPβ Int with *attPmm* and *attBmm*. These substrates could be paired and cleaved by Int, but not religated, and therefore trapped the complex in the swiveling-competent covalent intermediate state. Two classes of particles contain Int tetramers along the integrative reaction pathway: one in the pre-rotation cleaved-substrate conformation, synapsing an *attP* and an *attB* site, and one in the post-rotation not-yet-ligated-product state in which one half of the complex is rotated by ∼180° relative to the other (Figure 3A). While there are no covalent bonds between the two half-complexes to restrain rotation, there are likely energy wells at the 0° and 180° positions that are deep enough to favor these positions but shallow enough to allow rotation driven by thermal energy. The global resolutions for these maps were 7.18Å and 6.92Å, respectively, and local resolution extended to 4.69Å (Figure 3A and S9).

**Figure 3.**
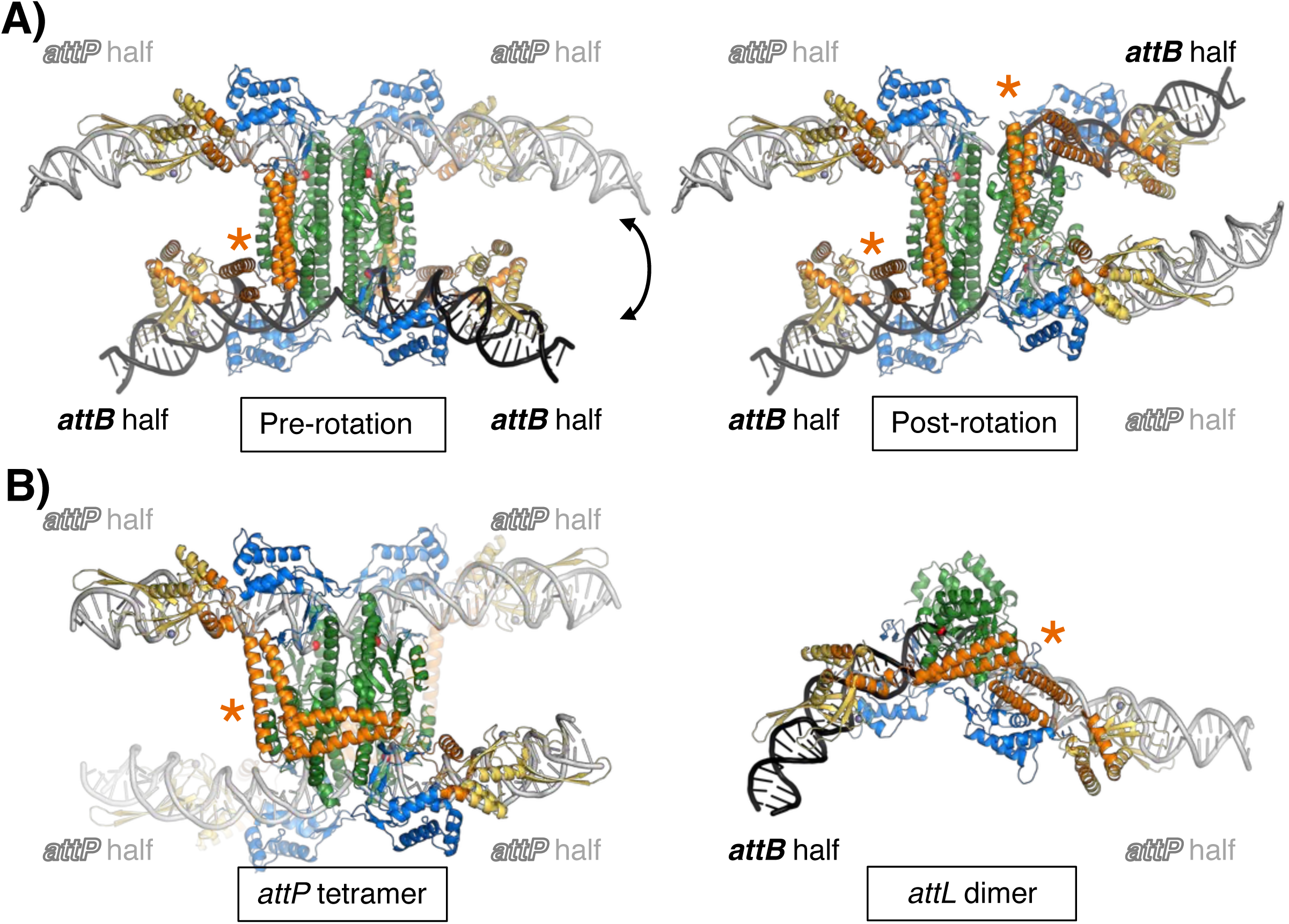
Structures of the synaptic complex during the integration (*attP* x *attB*) pathway (in the absence of the RDF). A. Structures of the SPβ integrase synaptic complexes in the pre-rotation (left) and post-rotation (right) states, captured in the covalent intermediate stage of integrative recombination. Colors are as in Figure 1, and orange asterisks mark approximately where CC:CC interactions occur. See also the closeup in Figure 5A. B. Additional complexes found in the same sample. Left: a tetramer synapsing 2 copies of *attP*, in a conformationally locked state with diagonal CC:CC interactions. Right: a dimer bound to cleaved product (*attL*) DNA.

Helix E is straight in these complexes and forms the central platform for rotation, as seen previously for small serine recombinases in the active tetrameric form^29,65^. Although most of the Int-DNA interactions with individual half-sites are similar to those seen in the *attL*-Int dimer complex, helix E and the catalytic domain form very different interactions with the DNA and with each other. Helix E is still docked into the minor groove, but due to DNA cleavage the DNA kinking and intercalation of the F147 and M154 side chains is no longer seen (Figure S10).

### The integrative synaptic complex is stabilized by inter-dimer CC:CC interactions

In these synaptic complexes, the CC motifs form inter- rather than intra-dimer interactions. The two cleaved DNAs lie on the outside of the synaptic complexes and are held together not only by tetramerization of the catalytic domains but also by inter-dimer handshakes between the CC motifs (Figure 3 and S9). The CC of each *attP*-bound subunit is sandwiched between the catalytic domain and the CC of its *attB*-bound partner. These tetramer-buttressing interactions occur on opposite sides of the pre-rotation complex but on the same side of the post-rotation complex. As discussed below, this geometry prevents formation of handcuffing intra-dimer CC-CC interactions before rotation and facilitates them after rotation.

An additional, unexpected 3^rd^ class of particles contained a tetramer of Int synapsing two *attP* sites, yielding a 4.81Å map (Figure 3B and S9). The presence of multiple *attP* sites in a cell is unlikely in a biologically relevant context, and these tetramers are unlikely to allow strand exchange because the CC:CC interactions occur diagonally across the tetramer, which is incompatible with rotation. Although the appearance of tetramers bound to two copies of *attP* was surprising, we found that SPβ Int – *attPmm* complexes do elute as tetramers in size exclusion chromatography (SEC) (Figure S4). Similar observations were recently reported for φC31 integrase^66^.

Conformational changes may be needed after rotation to bring the 3’ OH groups close enough for religation and to fully assemble the active site for catalysis. In all of the synaptic complexes described here, the two halves of each cleaved DNA are roughly co-linear, but the phosphoserines are ∼24Å apart (measured between P atoms), ∼10Å further than they would be in religated B-form DNA. Furthermore, the catalytically important arginines that are conserved among serine recombinases are not well ordered in these structures^67,68^. The proposed conformational changes may involve rigid-body motion of the DNA half-sites as well as adjustments within the catalytic domains. The *attB* half sites in our synaptic complexes tilt away from the catalytic domains, as does the DNA in most serine recombinase complexes^28,29,36,43,44^, but the *attP* half sites tilt towards the catalytic domains. This may reflect an intrinsic feature of *attP,* or it may reflect the inter-dimer CC-CC interactions pulling the *attP* DNA towards its *attB* partner. 3D variability analysis and 3D flexible reconstruction showed some flexibility in the relative orientation of the DNAs (see supplemental movies).

### The RDF dictates a different synaptic complex for excision

Three structures were determined from samples of the SPβ int-RDF fusion protein in complex with *attLmm* and were also trapped in the covalent intermediate state. We found three distinct classes of particles in these samples (Figure 4A and B): synaptic tetramers in the pre- and post-rotation states, and a smaller class of particles containing dimers bound to unligated product *attP* sites (Figure 4B, S11 and S12). We could not identify *attB*- bound dimers. The global resolution was 4.82Å for the pre-rotation state (with local resolution extending to 4.16Å), 5.00Å for the rotated state, and 7.1Å for the *attP*-bound dimer (Figure 4 and S11).

**Figure 4.**
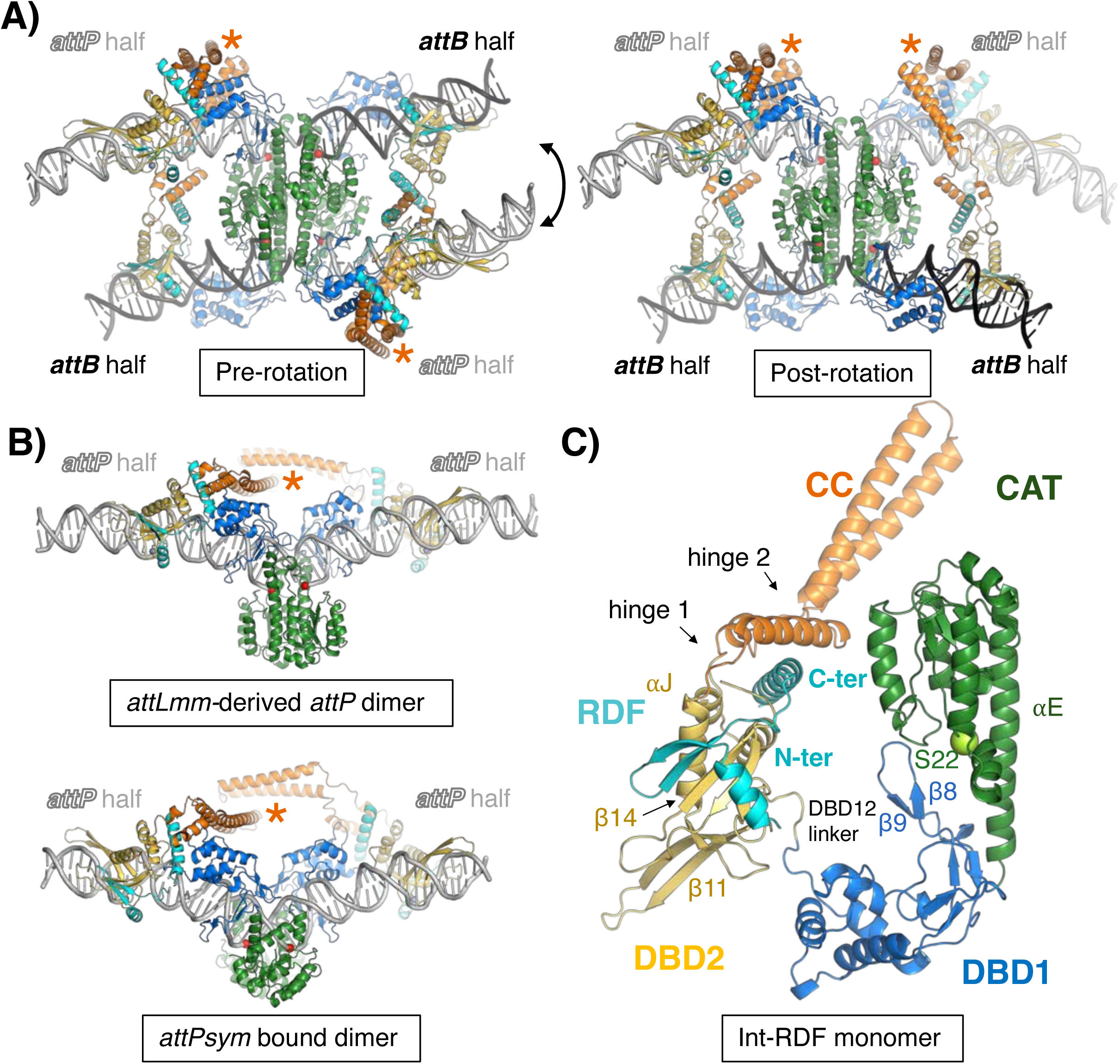
Structures of the Int-RDF synaptic complex during the excision (*attR* x *attL*) pathway. A. Structures of the SPβ integrase RDF fusion (Int-RDF) synaptic complexes captured in the covalent intermediate stage of excisive recombination. The pre-rotation state is shown on the left and the post-rotation state is shown on the right. Structures are colored as in Figure 1 with the RDF in turquoise. The orange asterisks indicate where the CC tips from the subunits bound to the upper and lower DNAs meet (see also the closeup in Figure 5B). B. *attLmm*-derived *attP* dimer (top) and *attP*-bound product dimer (bottom) structure, with orange asterisk highlighting where the CC tips of these two subunits interact to stabilize the dimer. C. Ribbon diagram of one SPβ Int-RDF fusion monomer isolated from the synaptic complex. The catalytic S22 is shown in yellow green sphere.

Comparison of the integrative and excisive synaptic complexes shows a major and unexpected difference: the CC – CC interactions that stabilize them occur on opposite sides (Figures 5 and S13). Nevertheless, the local interactions between the tips of the CCs are quite similar in both cases, except that “front” vs. “back” is switched - that is, without the RDF, the front of each *attB*-bound subunit’s CC interacts with the back of an *attP*-bound subunit’s CC, whereas in with the RDF, the backs of the *attB* CCs interact with the fronts of the *attP* CCs (Figure S7). Overall, RDF binding to Int does not simply disrupt the “handcuffed” *attL*- and *attR*-bound dimers that are the product of integration – it stabilizes a very different trajectory of the CC and promotes the formation of a different synaptic complex for excision.

**Figure 5.**
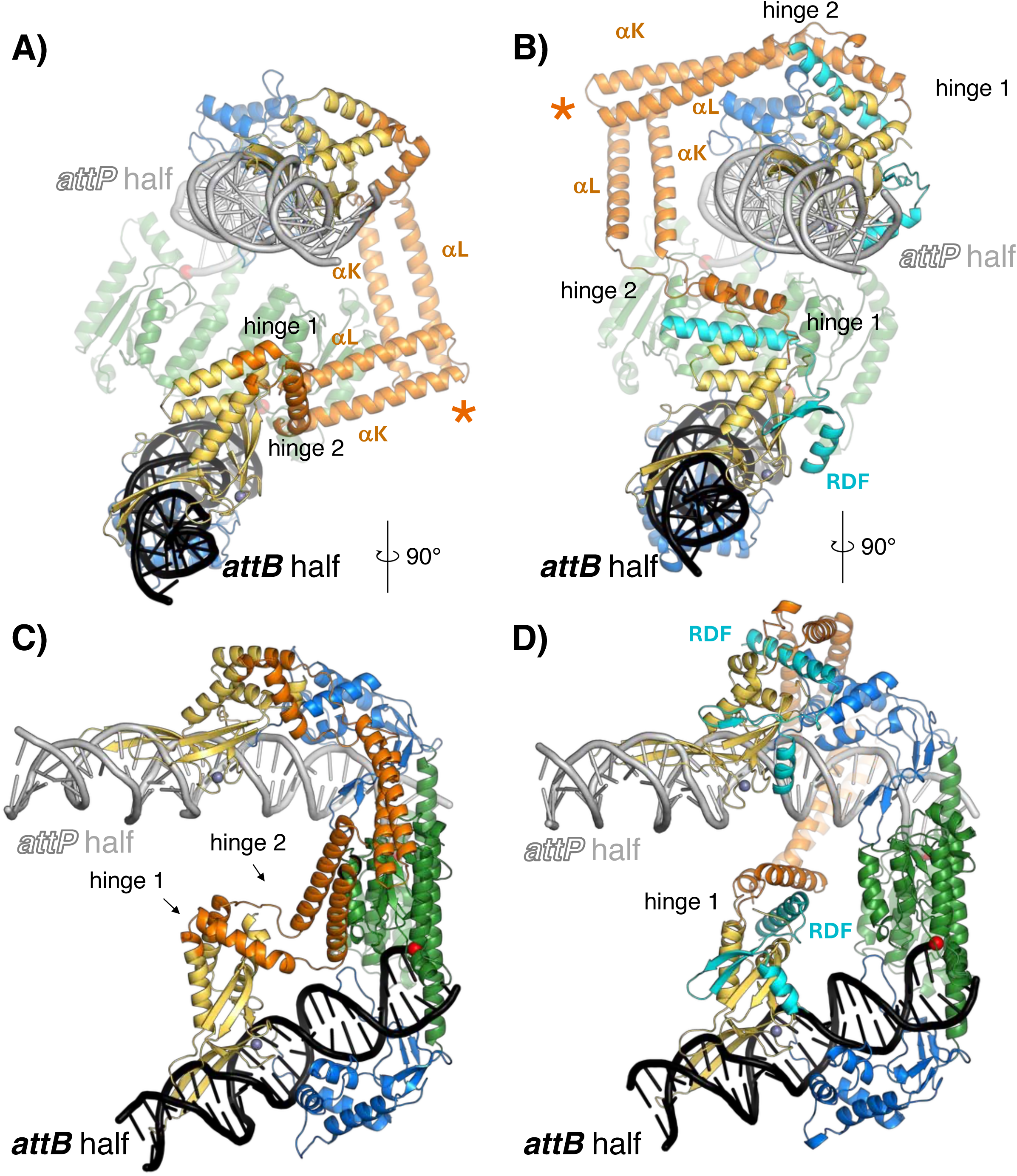
The RDF redirects the CC subdomain, dictating a different synaptic complex. A. DNA-end view of half of an integrative synaptic complex in the absence of the RDF, colored as in previous figures. The orange asterisk marks where two CC motifs meet. B. A similar view of half an excisive synaptic complex in the presence of the RDF. C. The same model as in (A), rotated ∼90° about the vertical axis. D. The same model as in (B), rotated ∼90° about the vertical axis.

### The RDF changes the pivot point for CC flexibility

The SPβ RDF is a small protein that binds both DBD2 and the DBD2-proximal portion of the CC, tying them together (Figure 4 and supplemental movie RDF_repositions_CC.mp4). Comparison of all the DBD2-CC conformations seen with and without the RDF shows that there are two hinges between DBD2 and the CC tip (Figure S14 and S15): hinge 1, located at the CC-DBD2 junction and hinge 2, further along the CC. Within each set of structures (-RDF vs. +RDF), hinge 1 remains rigid while hinge 2 serves as the flexible pivot point for CC motion. RDF binding to the inter-hinge region changes the conformation of hinge 1, which repositions hinge 2 and consequently changes the spatial range accessible to the CC tips. In general, in the absence of the RDF, hinge 2 lies on the front or back of the DNA (as seen in Figures S16) but in the presence of the RDF, it lies above or below the DNA (Figure S17). In *attL*- and *attR*-bound Int-RDF substrate complexes, the RDF-stabilized position of hinge 2 would prevent intra-dimer CC-CC handcuffing. However, within the synaptic tetramer, the subunit rotation that converts substrates to products changes the relative positioning of the *attP*-bound subunits’ hinge 2s from diagonally opposed across the tetramer to close proximity, which facilitates handcuffing of the *attP*-bound product. Although we could not determine an attB-bound product structure, Figure S17 suggests that it is probably also handcuffed.

We also noted that the RDF appears to destabilize the integration-competent trajectory of the CC while stabilizing the excision-competent trajectory. In the absence of the RDF, the helical inter-hinge segment (aa 381-397) N-terminal to the CC packs against DBD2 in the same location where a loop of the RDF binds, effectively shielding a small hydrophobic region (Figure 5C, D, and S18). This shielded pocket includes the extreme C-terminal region of the integrase (aa 530-545), notably F532, which is essential for catalyzing excision reaction but largely dispensable for integration^69^. Additionally, the CC-proximal ends of helices J (aa 376-379) and M (aa 474-484) within DBD2 are unwound in the presence of the RDF and form part of the inter-hinge region (Figure 5A and S18A). Both the experimental maps and AlphaFold models support this helical extension in the absence of the RDF (Figure S18). The endpoints of these two helices are also variable among the 4 copies in the asymmetric unit of the crystal structure of the LI integrase DNA binding domains^48^.

RDF binding displaces the inter-hinge helical segment (aa 381-397) by inserting its C-terminal helix and the loop that precedes it into the DBD2 hydrophobic region. This displacement destabilizes the helix M extension, exposing hydrophobic residues such as I472 and V477 to form new contacts with F40 of the RDF, thereby stabilizing the repositioned hinge 2 conformation. Concurrently, repositioning of the integrase C-terminus allows F532 and F534 to engage in close hydrophobic contacts with RDF residues including L37 and F40 (with additional support from Y41 and F44), stabilizing the new conformation of hinge 1. These hydrophobic interactions between DBD2 and the RDF align with functional data from Abe et al. 2021, who described the integrase C-terminus as a potential component of a “molecular toggle switch” controlling recombination directionality.

### CC-CC handcuffing of the *attP*-bound integrase-RDF dimer

Although a low-resolution structure of an *attP*-bound dimer structure was derived from the *attLmm*- SPβ int-RDF sample, the central mismatches trapped it in the cleaved state rather than the final religated state. We therefore directly formed product dimer complexes with uncleaved DNA by mixing the SPβ int-RDF fusion protein with *attPsym* and *attBsym* substrates. Under these conditions, only an *attP*-bound dimer species was observed (Figure S19), which yielded a structure at an overall resolution of 7.1Å. In both *attP*-bound dimers, the two copies of hinge 2 lie on the top face of *attP*, placing the CCs and their tips in close proximity, and enabling intra-dimer CC-CC interactions that likely drive or stabilize tetramer-to-dimer dissociation (Figure S17). No CC-CAT interactions are seen in either of these dimers.

Although both the *attLmm-* derived *attP* dimer and *attPsym-* bound dimer bind essentially identical *attP* half sites, they reveal markedly different degrees of CAT domain ordering and positioning (Figure S11, S12 and S19). In the *attLmm*-derived *attP* dimers –where the DNA is cleaved – the CAT domains are largely disordered, even after focused classification, reflecting high conformational flexibility around the active site. In contrast, the *attPsym* bound dimer, assembled on uncleaved *attP* DNA, exhibits significantly improved ordering of the CAT domains (Figure S12).

In the best-resolved 3D class, the dimer formed by the two CAT domains closely resembles the unusual (for serine recombinases) CAT domain dimer seen in the handcuffed *attLsym*-bound dimer in the absence of the RDF (Figure 2 and S12). Thus, while the dimer containing cleaved DNA samples a highly dynamic ensemble of CAT conformations with no preferred stabilization, the dimer containing uncleaved DNA is biased toward an open, E-helix bent, and catalytically inactive state.

Comparison with tetrameric synaptic complexes uncovers a conformational allosteric switch centered on the helix E. In active tetramers, the E helices are straight, allowing aromatic residues F138 and F141 to engage in pi-pi stacking across intra- and inter-dimer interfaces (Figure 6B and S10B). These residues lock the assembly in the rotationally competent state, providing a flat hydrophobic platform for subunit rotation. Although the catalytically critical arginine side chains are not well-ordered around the phosphoserine protein-DNA linkage in these complexes, their close proximity implies that only relatively minor changes are needed for catalysis of the religation reaction. Upon tetramer-to-dimer dissociation after religation, bending of helix E repositions these same aromatic side chains (F138 and F141), thereby disrupting the dimer-of-dimers interface and stabilizing a catalytically inactive conformation of the CAT domain (Figure 6A, B and S10B).

**Figure 6.**
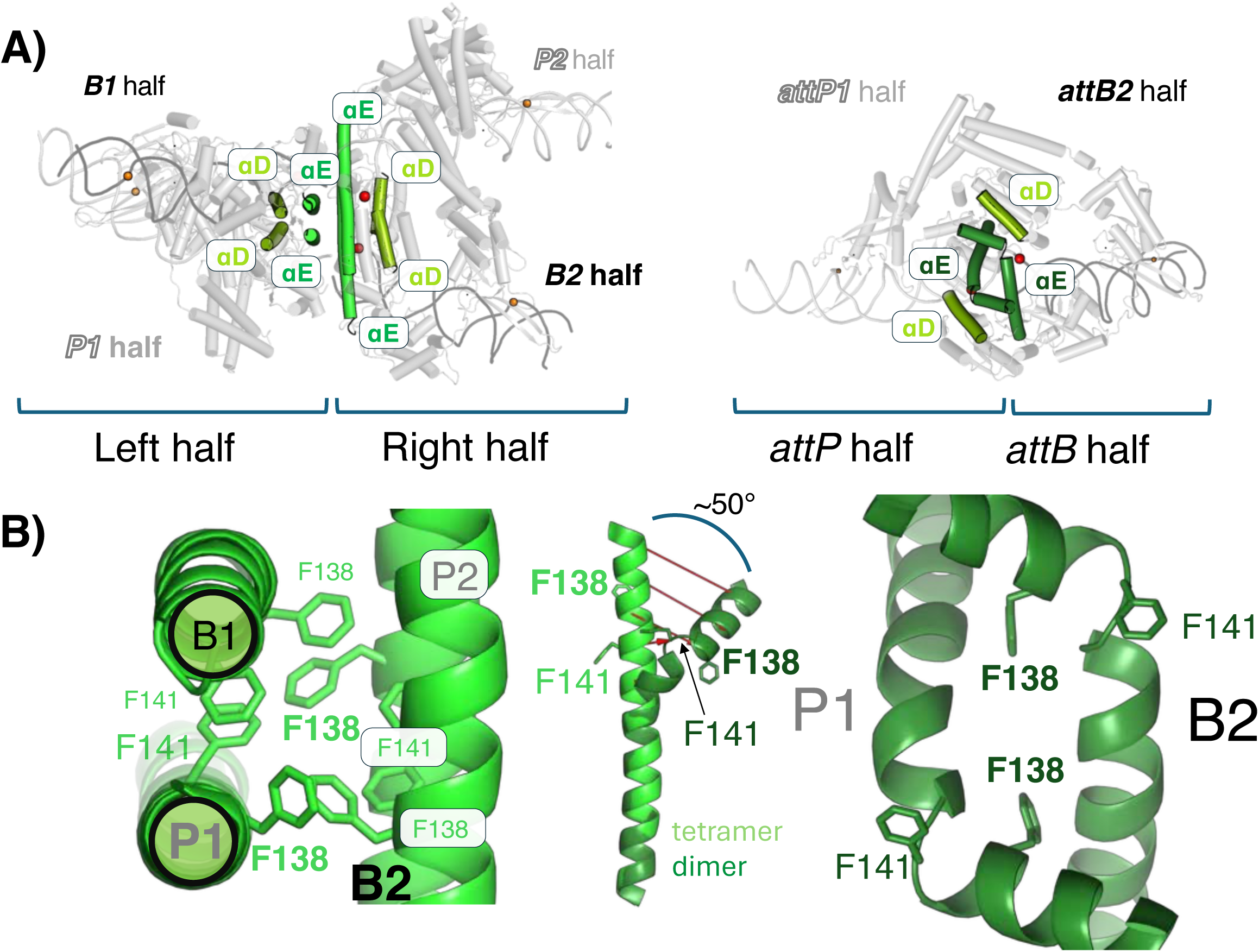
Bending of E-helices reorganizes the inter-subunit interaction surface in the dimer. A. Comparison of the post-rotation synaptic tetramer (left) and *attL*-bound dimer (right). Two helices of the CAT domains are shown in color: αD (residues 101-114; yellow-green) and the αE (residues 130-165; light green in the tetrameric complex; dark green in the dimeric complex). B. Close-up views of the complexes in A, in the same orientations. Two key interface residues, F138 and F141 are shown as sticks. In the dimer form, the E-helices are bent by approximately 50° and partially unwound at residues 140-145. The inset in the center shows one E-helix from each structure superimposed by aligning residues 146-164. Atomic displacements (≥ 4Å) between corresponding Cα atoms of the straight and bent E-helices are indicated by red arrows generated using modevectors, with every two Cα atoms skipped.

Thus, helix E bending, facilitated by post-rotation CC-mediated dimer stabilization, serves as a key allosteric switch that couples synaptic complex to catalytic inactivation, thereby ensuring strict unidirectionality of the recombination reaction.

### Mutational analysis reveals the RDF-CC interface is essential for redirecting CC trajectory

We employed an *E. coli*-based promoter inversion assay to investigate the functional importance of Int-RDF interactions (Figure 7 and S20)^53,70^. Int flips a promoter flanked by *attP* and *attB* sites to drive GFP rather than RFP expression, while the Int – RDF fusion acts on the *attL* and *attR* sites in the product plasmid, flipping the promoter back to RFP expression. Use of the Int – RDF fusion avoids artifacts due to unequal protein expression or weakened protein – protein affinity. Figure 7B shows that the WT proteins drove recombination to completion after only 2 hours of expression. Deletion of the N-terminal helix of the RDF (Int-RDF ΔN-ter helix 1-13), which interacts only with DBD2, had no effect on RDF function in this assay. In contrast, the RDF’s C-terminal helix (Int-RDF ΔC-ter helix 37-58), which packs between DBD2 and the inter-hinge region of the CC, is essential: when this helix was deleted, the fusion protein acted as a slow version of WT Int without RDF, eventually converting all substrates to the *attL*/att*R* configuration. Similar results were obtained for two less drastic changes in the C-terminal helix: pairwise mutations L37D/F40D and A45D/L49D. The first pair flank Int F532, which was previously shown to be important for directionality control^69^ as discussed above, and the second pair interact directly with the inter-hinge segment. These data support the hypothesis that restraining the inter-hinge region of the CC is critical to RDF function.

**Figure 7.**
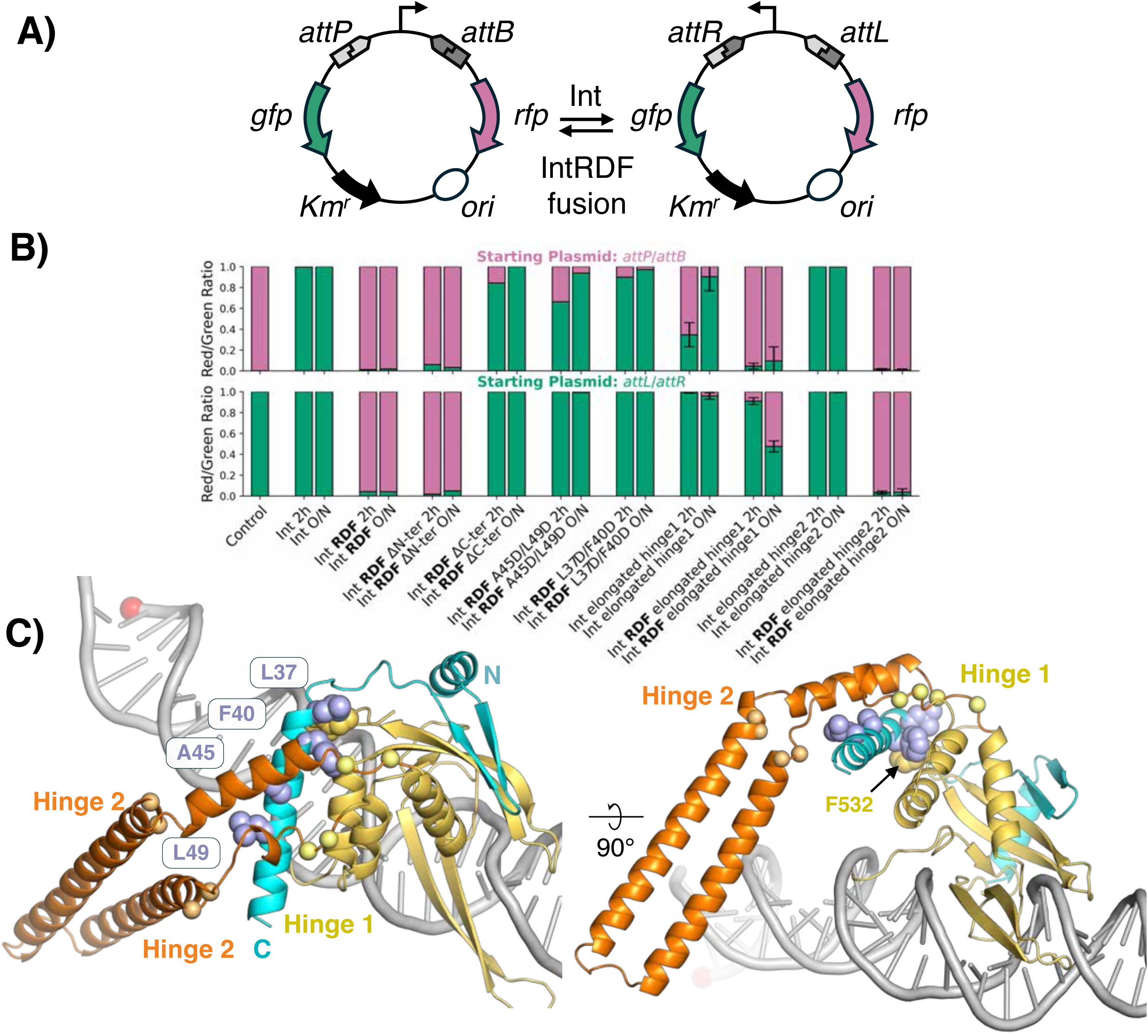
Mutational analysis of RDF and integrase hinge regions. A. Schematic diagram illustrating the *in vivo* DNA inversion recombination assay. B. Quantitative analysis of inversion efficiency. Stacked bar charts illustrate the proportion of red and green E. coli colonies, reflecting the state of the reporter plasmid after inversion assays. All mutants were tested on both starting plasmids (*attP*/*attB*, top and *attL*/*attR*, bottom). Int or Int-RDF fusions were expressed for 2 hours or overnight (see methods for details). ΔN-ter: deletion of the first helix of the RDF (Δ residues 1-13). ΔC-ter helix: RDF Δ residues 37-58. Elongated hinge 1: addition of GGSGSSG between D379 & L380 and between N479 & N480; Elongated hinge 2 : addition of GGSGSG between S401 & N402 and between D467 & T468. C. Close-up views of DBD2 and the RDF from at *attP*-bound subunit. Side chains that were mutated are shown as spheres (light blue) and the C⍺ atoms of residues flanking the hinge-elongating insertions are shown as spheres (yellow; hinge 1, light-orange; hinge 2).

We also tested the effects of lengthening the hinges in SPβ integrase. Hinge 1 was lengthened by inserting GGSGSSG into both the outgoing and returning chain (Figure 7C) and hinge 2 was similarly lengthened by inserting GGSGSG (see Figure 7C). The effects of elongating hinge 1 were minor: both integrative and excisive reactions were slower, and the int-RDF fusion was less cleanly switched – excisive recombination was still favored, but not as decisively. Elongating hinge 2 had no detectable effect on Int or Int-RDF function in our assays. Although elongating the CC of LI integrase by ∼two helical turns weakened recombination efficiency and directionality^3^, our results suggest that the most important function of the RDF is to change the pivot point of the CC rather than simply to restrain how far it can extend.

### Additional protein-protein interactions

The SPβ synaptic complexes include additional protein-protein interactions that were not predicted from partial structures and previous modeling (Figure S21). E258 of the long DNA-binding loop of one subunit’s DBD1 could form a salt bridge with the catalytic domain of the subunit bound to the DNA on the opposite side of the synapse, potentially helping to stabilize the synaptic complex. When the RDF is present, but not when it is absent, the CC from the *attP*-bound subunits contacts a small hydrophobic patch on DBD1 of the same subunit. Similar CC-DBD1 contacts are made by both *attP*-bound subunits within the tetramer and by one subunit within the *attP*-bound dimer. An initial investigation of mutations of E258 and the DBD1-CC contacts found little to no effect in a promoter-flipping assay after two or four hours of protein expression (Figure S22 and S23). However, these mutations could affect protein stability or reaction rates less dramatically than could be detected in our assays.

Contacts between the CC and CAT domains occur only in the absence of the RDF. The details vary between the tetramer and dimer states, but P54 (CAT domain) and F426 (CC) are included in both sets of contacts. Interestingly, that region of the catalytic domain is involved in activating contacts from other proteins or DNAs in small serine recombinases^27^. Because F426 also participates in CC-CC contacts, we could only test the importance of CC-CAT contacts by mutating the CAT domain. A triple mutation (T49A, I53D, P54D) was deleterious in both reaction directions, suggesting that it may have affected catalytic activity. However, it was less deleterious to the excision reaction, where CC-CAT domain contacts do not occur, than to the integration reaction, where CC-CAT domain contacts do occur. This result supports the idea that some mutations may affect the two reaction pathways differently.

## Discussion

Biology often requires that site-specific DNA recombination reactions proceed to completion in a particular direction even when the net number of phosphodiester bonds does not change. A number of different solutions to this conundrum have evolved. For phage integrases from the tyrosine recombinase family, additional DNA-binding proteins are used to set up a synaptic complex from which recombination is downhill in terms of conformational energy^25^. The closely related XerC/XerD tyrosine recombinases cooperate with each other and with the FtsK translocase to resolve chromosome dimers. The same two tyrosine recombinases can also resolve plasmid dimers, but in that case use additional DNA binding proteins to set up a synaptic complex in which recombination is downhill in terms of DNA supercoiling energy^71^. Small serine recombinases that resolve plasmid dimers (or replicon fusions in the case of transposon resolvases) also rely on additional DNA binding proteins to sense and be driven by DNA topology^46^. In contrast, unidirectional recombination by LSIs does not require a particular DNA topology nor does it require additional DNA binding proteins.

Understanding the control of reaction directionality for the LSIs has been a long-standing challenge. Our series of SPβ Int and Int-RDF complex structures can be assembled into a movie of an LSI "in action" (Supplementary Movie Integration_Excision.mov). This work provides direct evidence for an updated model, diagrammed in Figure 8 and Supplementary Figure 24, that agrees with current models for integrative recombination^51^ but fundamentally changes the model for excisive recombination. Unidirectionality in both pathways is explained by the interplay among 3 key features: the CC, the RDF, and the difference between *attP*- and *attB*- half sites. These control the oligomeric state of the integrase and consequently the conformation and activity of the catalytic domain.

**Figure 8.**
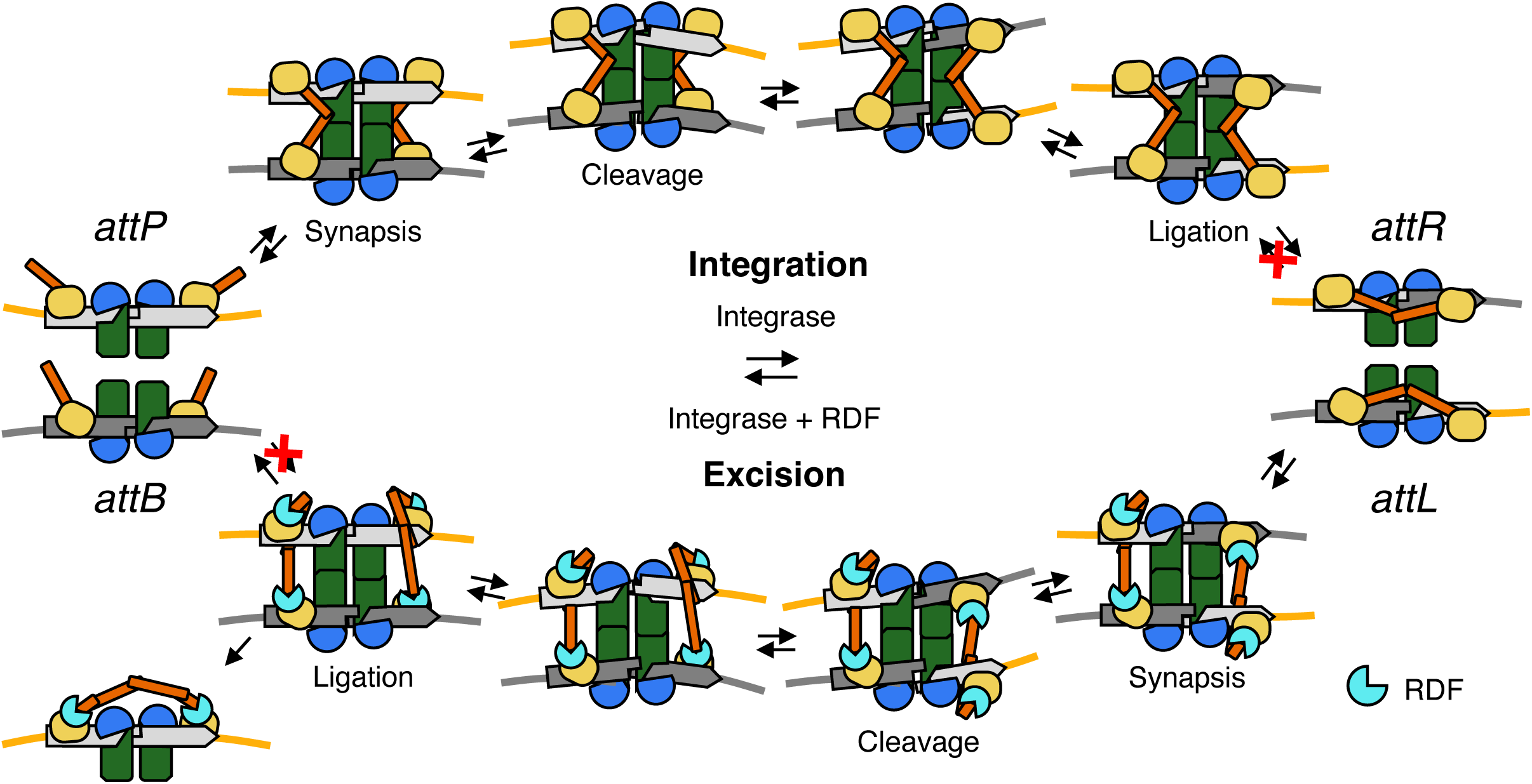
Schematic model of how CC-CC interactions and the RDF mediate directionality control. Pathways for integrative (upper) and excisive (lower) recombination are diagrammed with the protein domains and DNAs colored as in previous figures. CC subdomains (orange) buttress allowed synaptic complexes by mediating interactions between the upper and lower DNAs, but they can also re-arrange after recombination to form intra-dimer interactions that trap the products and prevent the reverse reaction from occurring (red X’s). The integration pathway is conceptually similar to that proposed by Rutherford and Van Duyne^5^ but the CC motifs in particular are redrawn to match the experimental data presented here. The RDF redirects the trajectories of the CC subdomains so that a different type of synaptic complex (between *attL* and *attR*) and a different product dimer (*attP*-bound) are stabilized.

For both reaction pathways, the CC:CC interactions stabilize the synaptic complex, then stabilize or “handcuff” at least one, if not both, product dimers. Dimer handcuffing prevents reaction products *(attL* and *attR* for integration and *attP* and *attB* for excision) from forming synaptic tetramers, thus preventing the back reaction. Integration and excision are driven forward by similar mechanistic logic, but occur within structurally different complexes. In the initial substrate state of both types of synaptic complex, the critical 2^nd^ hinges that are the CCs’ pivot points are located such that the CC tips can only interact with subunits bound to a partner duplex, thus stabilizing synapsis. However, rotation to the product state brings the 2^nd^ hinges of subunits bound to a single product duplex close enough that they can swap interaction partners to handcuff the product dimer.

The RDF not only unlocks the handcuffing of the *attL*- and *attR*- bound dimers that are the products of integrative recombination (as previously proposed), but also changes the location of the CCs’ 2^nd^ hinges. The RDF therefore favors a synaptic complex with different geometry from that used in integrative recombination, and allows trapping of the *attP*- (and probably *attB*-) bound dimers that are the products of excisive recombination. The use of structurally different synaptic complexes for integration vs. excision is conceptually similar to the regulation of phage lambda integrase (a tyrosine recombinase) even though the two systems differ wildly in mechanism and complexity^25,26^.

The work presented here provides a framework for engineering LSI-RDF pairs for use as genetic switches and genome editing tools. We suggest that optimization of reaction efficiency in a particular pathway could be streamlined by focusing on protein segments that adopt different interactions and/or conformations in the excisive vs. integrative pathways. The amino acid sequences of those segments probably reflect evolutionary compromises between pathways – not only to balance the optimal efficiency in both directions, but also to balance optimal inhibition of the “wrong” reaction in both cases. They are therefore likely to be hotspots for mutations that optimize reaction efficiency in one direction without regard for the other, as seen in the clustering of efficiency mutations in CC region reported in^72^ and for mutations that loosen directionality control, as seen in CC region in ^52^.

The creation of integrase variants with altered and/or enhanced sequence specificity is also of interest to the broader community. Two features of SPβ integrase that are of particular interest are the β8-9 hairpin extending from the inner portion of DBD1 and the β11-12 hairpin of DBD2. These hairpins are likely to be important in sequence recognition because they dock into the major groove and contact conserved sequence motifs. Furthermore, they are likely to be amenable to mutation because they protrude from the core DBD folds. Although the overall folds of LSIs are predicted to be highly conserved, variability in hairpin length (Figure S8) may contribute to differences in the stringency of sequence specificity and in overall *att* site length.

Consistent with this idea, SPβ integrase-DNA contacts extend across 46 and 54 bp of *attB* and *attP*, respectively, which is longer than is often assumed to be sufficient for LSI binding, and extensive characterization of LI integrase revealed very little sequence preference for its DBD1, consistent with its much shorter β8-9 hairpin.

A recent preprint describes integrative-pathway structures of φC31 integrase^66^ which confirm that, like SPβ integrase, it follows the generalized model for integrative recombination proposed by Rutherford and Van Duyne, in which CC-CC interactions stabilize both the initial synaptic tetramer and the product dimer. However, some architectural differences are apparent when comparing dimers, tetramers, and *att* site binding. The largest differences between the SPβ and φC31 structures are in their *attL*-bound dimers. Although the CCs contact the catalytic domains in both dimers, different regions of the catalytic domains are involved. Structural variations within dimers are likely biologically permissible because they are not the catalytically active species, allowing for conformational fluctuation without compromising the overall recombination pathway.

Beyond flexibility within the protein framework, SPβ and φC31 also display a minor difference in the relative placement of DBD2-binding motifs within their cognate *att* sites. SPβ integrase exhibits a 4 bp shift in DBD2 placement between *attP* and *attB* sites, rather than the 5 bp shift predicted for some LSIs and seen in φC31 structures^66^. Similar variations probably occur for other LSIs, given the variability in DBD1-2 linkers, hinge and inter-hinge regions and predicted CC length (e.g. Figure S8). A functional integrase only requires that *attP* can be differentiated from *attB* and that the appropriate CC-CC interactions can form throughout the recombination pathway. These observations suggest that future bioinformatic analyses of *att* sites should consider the shift in DBD2-binding motifs as a variable.

Additionally, less dramatic differences are apparent between synaptic tetramer structures for φC31 and SPβ integrases that may reflect different reaction stages. For φC31, an active site mutant was used to trap the pre-DNA-cleavage state, whereas for SPβ mismatched central dinucleotides were used to trap the post-cleavage state. The φC31 tetramer displays a smaller crossing angle between E helices at the center of the tetramer, a less extensive interface between the *attP*- and *attB*- bound subunits, and a greater distance between the two DNA duplexes. It is likely that there are multiple sub-states along the pathway from initial synapsis to the swiveling-competent cleaved state, with the actual moment of catalysis occurring somewhere between those two points. If the catalytic sites were always fully assembled and active, the system would risk: (1) premature DNA cleavage before verification of a proper *attP*- and *attB*- pairing and (2) hydrolysis of the phosphoserine linkage during subunit rotation. The latter consideration explains why the catalytic arginines were not well ordered around the phosphoserine in the SPβ tetramers.

For LSI-catalyzed DNA recombination to proceed efficiently, the reaction pathway must avoid deep energy wells before the final handcuffed product dimer state is reached. This may be achieved by balancing favorable macromolecular interactions with conformation strain, including the bending of the DNA. For example, we noted an unusual tilt of the *attP* DNAs towards the catalytic domains in post-cleavage SPβ tetramers that was not seen in pre-cleavage φC31 tetramers. We propose that there is a balance between having all four DBD2s bound to DNA with both CC:CC pairs interacting across the synapse vs. conformational strain in the DNA near the active site. Such a balance would explain the difficulties reported in obtaining pre-cleavage φC31 particles with well-ordered CC:CC pairs on both sides of the tetramer. DNA cleavage may relieve some conformation strain, driving the tetramer to the cleaved and swiveling state. Religation of the DNA in the product configuration would re-introduce the strain, but would also favor dissociation into dimers, which are then trapped by CC:CC handcuffing. In contrast, religation in the substrate configuration would re-introduce the strain with no energetic compensation.

Overall, the LSI reaction pathway is remarkably simple in its protein and DNA requirements, yet entails a delicate energetic balance in which numerous protein—protein and protein—DNA interactions act in concert to guide the reaction forward. The protein-protein interactions must be strong enough to stabilize key intermediates and enforce directionality, yet transient enough to avoid kinetic trapping prior to product formation. Directionality thus arises from dynamic control of conformational strain, DNA bending, and selective interaction switching rather than from any single dominant interaction. Together, this architecture enables tuning of LSIs to combine stringent reaction control with catalytic efficiency, and provides a general framework for understanding, and ultimately engineering directionality in complex DNA recombination systems.

## Acknowledgments and Funding

We thank the members of Research Computing Center (RCC) at the University of Chicago for providing their services. This work was completed using the Beagle3 high-performance computing cluster funded by the NIH through grant 1S10OD028655-01. Cryo-EM datasets were collected at the University of Chicago Advanced Electron Microscopy Core Facility (RRID:SCR_019198). Molecular graphics and analyses performed with UCSF ChimeraX, developed by the Resource for Biocomputing, Visualization, and Informatics at the University of California, San Francisco, with support from National Institutes of Health R01-GM129325 and the Office of Cyber Infrastructure and Computational Biology, National Institute of Allergy and Infectious Diseases; compiled and configured by SBGrid^73^. This publication is primarily supported by the National Science Foundation and UK Research and Innovation [collaborative grant NSF/BIO 2223480 and UKRI/BBSRC BB/X012085/1 to P.A.R. and F.J.O]. Funding for open access charge: National Science Foundation.

## Author contributions

Conception and design, Methodology: PAR, FJO, HS

Funding acquisition: PAR, FJO

Data acquisition: HS, AJB, JRF, YP, TPR

Data analysis: PAR, FJO, HS, JRF

Drafting and revision: PAR, FJO, HS

## Data and materials availability

All EM movies have been deposited at the Electron Microscopy Public Image Archive (EMPIAR). The cryo-EM maps and coordinates of the complexes determined in this study have been deposited into the Electron Microscopy Data Bank (EMDB) and the Protein Data Bank (PDB), respectively, with the following accession codes: EMDB-47288 and 9DXH for Int in pre-rotation state; EMDB-47289 and 9DXJ for Int in post-rotation state; EMDB-47290 and 9DXK for Int *attP*-bound tetramer; EMDB-72552 and 9Y66 for Int in *attL*-bound dimer; EMDB-47284 and 9DXD for Int-RDF in pre-rotation state; EMDB-47286 and 9DXF for Int-RDF in post-rotation state; EMDB-47287 and 9DXG for Int-RDF *attP*-bound dimer in cleaved state; EMDB-72632 and 9Y6V for Int-RDF *attP*-bound dimer in uncleaved state.

## Methods

### Cloning and plasmid construction

The DNA sequence of SPβ Integrase was optimized for expression in *E. coli* and cloned between the NdeI and XhoI sites of pET-28a(+) (Novagen). The proteins expressed from this vector have an N-terminal hexahistidine tag followed by a thrombin cleavage site (the tag was not cleaved off, but was not visible in the density maps). The DNA sequence encoding the SPβ integrase–RDF fusion protein was made by joining the coding sequences for SPβ integrase and its RDF via a 54 bp DNA linker sequence between the SpeI and XhoI sites in pET28-a(+). The DNA linker sequence is translated to a flexible 18-residue linker (TSGSGGSGGSGGSGRSGT) between the C terminal residue of SPβ integrase and the second amino acid residue in the RDF. Mutants of Integrase and Integrase-RDF were generated using site-directed mutagenesis, and all constructs were verified by whole plasmid sequencing of the expression vectors.

### Protein expression and purification

Expression plasmids containing SPβ Int or SPβ Int-RDF were used to transform *E. coli* BL21(DE3)pLysS competent cells. The transformed cells were grown at 37 ℃ in LB broth supplemented with 50µg/ml Kanamycin and 100 µM ZnSO_4_, until OD_600_ reached 0.7. Protein expression was induced with isopropylβ-D-1-thiogalactopyranoside (IPTG) at final concentration of 0.5 mM, and the cultures were grown overnight at 20 ℃. The cells were pelleted by centrifugation at 8,000 r.p.m. in a Fiberlite F9-6X1000LEX rotor for 7 minutes at 4 ℃ and stored at -20 ℃.

Cell pellets were resuspended in 50ml per liter of original culture of Ni A buffer (50mM Sodium Phosphate, 1M NaCl, 5% Glycerol, 1mM Tris(2-carboxylethyl) phosphine (TCEP), pH 7.5) plus 1 tablet of complete Mini EDTA-free protease inhibitor cocktail (Roche) and 200 µg/ml lysozyme, then lysed by ultrasonication in 3 1-minute pulses in an ice bath. The lysate was clarified by centrifugation at 18,000 r.p.m. in an SS-34 rotor for 1 hour at 4 ℃. The supernatant was collected, filtered and then loaded on a 5ml Ni Sepharose HisTrap HP affinity column (Cytiva). The column was washed with 95% Ni A / 5% Ni B buffer (Ni B = Ni A plus 500 mM Imidazole pH 7.5) until the baseline was flat, then eluted by a gradient from 5% to 100% Ni B) over 40’ with 2ml/min and 2ml/fraction. The protein peak fractions were analyzed on a Novex 8-16% Tris-Glycine Minigel (Invitrogen). The cleanest fractions were pooled and dialyzed into Ni A buffer at 4 ℃ O/N, then rechromatographed on the Ni column as before. The best fractions were pooled, then diluted into Heparin A buffer (20mM MES, 5% Glycerol, 1mM TCEP, pH 5.5) to a final salt concentration of 200-300mM, and then loaded onto a 20ml Heparin column (Heparin FF, 16/10, Cytiva), washed with 15% Heparin B buffer (Heparin A buffer plus 2M NaCl, pH5.5) and eluted with a gradient from 15% to 75% B over 90’ at 2ml/min and 2ml/fraction. The cleanest peak fractions were and concentrated to a small volume, then dialyzed into 20 mM Tris-Cl pH 8, 200 mM NaCl, 2 mM TCEP, and 20% glycerol. The protein concentration was determined by A_280_ and the presence of contaminating endonucleases was ruled out by incubating the sample with supercoiled pUC19 and 10mM MgCl_2_ followed by agarose gel electrophoresis.

### List of DNA substrates

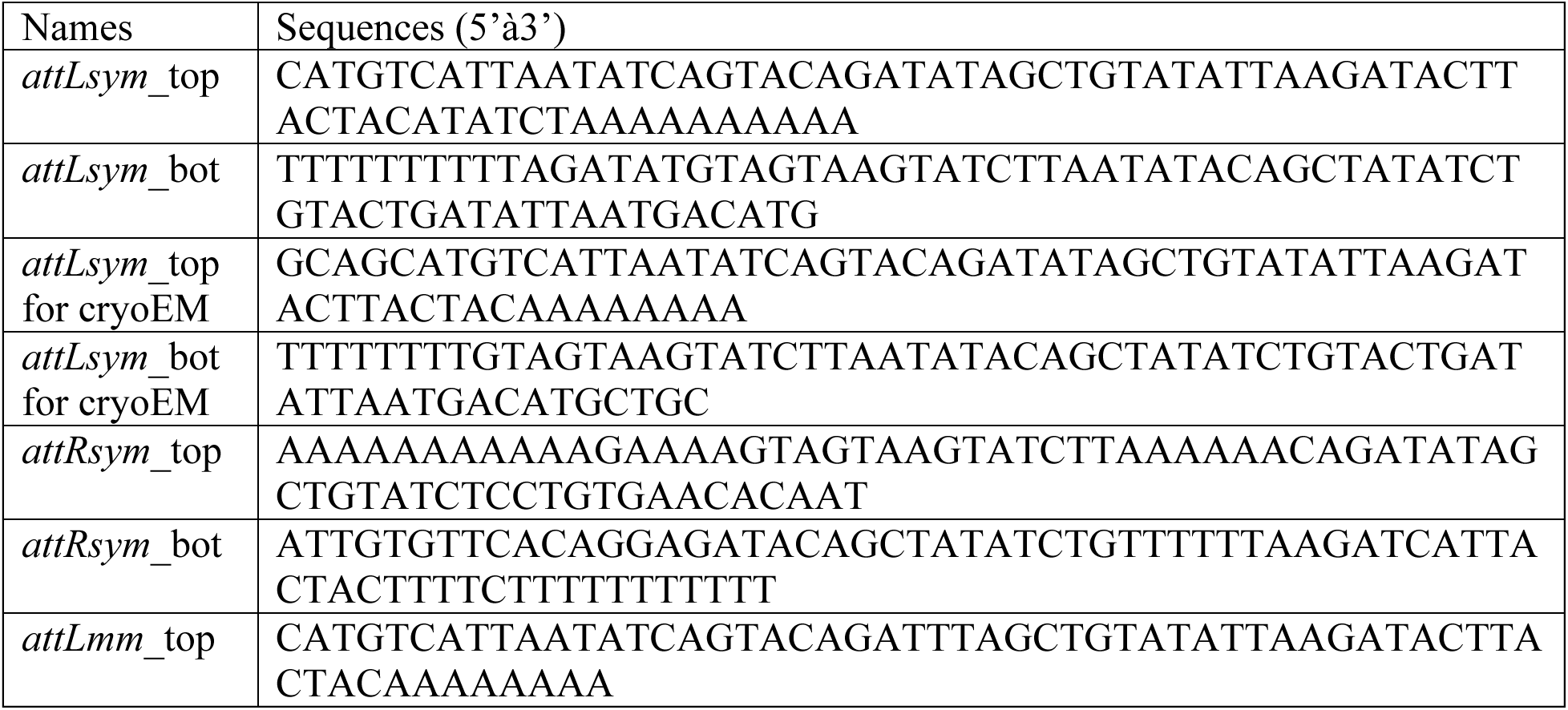
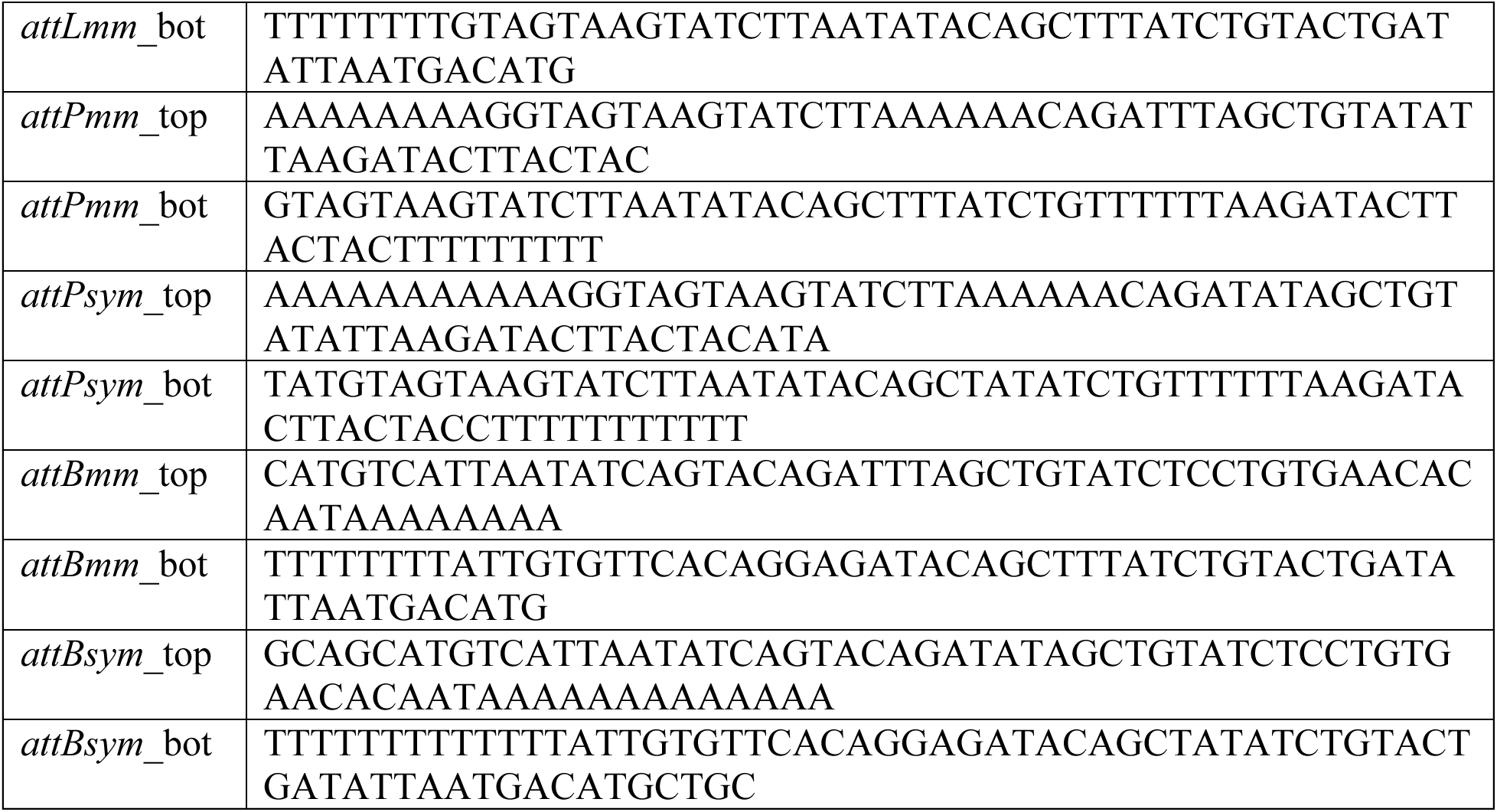

For the fluorescently labeled oligos, some of the oligos listed above were custom ordered with 5’ fluorescein label or 3’ Cy5 label from Integrated DNA technologies (IDT). Complementary oligos were mixed at a final concentration of 1 mM in TE buffer containing 100 mM NaCl and annealed by incubating at 80 ℃ for 20 minutes. Afterward, the water bath was turned off and samples were left in it to cool to room temperature.

### *In vivo* integrase mediated inversion assay

The design and protocol for the *in vivo* recombination assay were similar to those previously described^53,70^.The substrate for *in vivo* intramolecular recombination (inversion) is a reporter plasmid (kanamycin resistance) in which *attP*/*attB* or *attR*/*attL* sites are in inverted orientation and flank a constitutive promoter that drives either expression of GFP or RFP (Figure 7A). To assay recombination activities *in vivo*, *E. coli* DS941 cells were cotransformed with a substrate plasmid and an arabinose-inducible expression vector, pBAD33 (chloramphenicol resistance) containing the gene of interest (Int, Int-RDF fusion or its mutants cloned between NdeI and XhoI sites). To do so, 2µl of each plasmid at 20ng/µl was incubated with 100ul chemically competent DS941 cells on ice for 5’, then heat shocked at 37 ℃ for 5’, then iced for 2’. 200µl LB+0.2% glucose was then added, cells were allowed to recover 37 ℃ for 2 hours, then added to 10ml LB+0.2% glucose+50µg/ml kanamycin (kan) +30µg/ml chloramphenicol (cam) and grown overnight at 37 ℃. All liquid cultures for these experiments were shaken at 200 rpm. Glucose was added to suppress protein expression, and we noted that the overall experiment worked best if the overnight culture reached an OD_600_ of > 1.5 – if not it was best to restart with freshly made competent cells.

200µl of the overnight culture was used to inoculate 10ml LB + 0.2% Glucose + kan (50 µg/ml) + cam (30µg/ml), then grown at 37 ℃ for 1.5 hours, at which point OD_600_ was ∼0.4-0.5. Cells were spun down to remove the glucose, then resuspended in 10ml LB + 0.2% L-Arabinose + kan (50 µg/ml) + cam (30 µg/ml) and incubated at 37 ℃ for 2 hours or overnight, after which the plasmids were recovered by miniprep. 1µl of miniprepped plasmid was then used to transform 50ul chemically competent DS941 cells: 5’ on ice, 5’ heat shock at 37 ℃ and 2’ on ice then recovery in 500µl LB at 37C for 90’. We choose to isolate and then retransform the product plasmids to separate out individual copies of the substrate plasmid, which carries a pSC101 origin and can be present in multiple copies per cell. Two different volumes of each culture (20µl and 100µl) were then plated LB + Kan (50 µg/ml) agar plates and incubated at 37 ℃ overnight. Plates were visualized in a chemi-doc (Bio-Rad, Hercules, CA, USA) with blue epi illumination and a 530nm±28nm bandpass filter for GFP visualization and red epi illumination with a 695nm±55nm bandpass filter for RFP. Exposure times were set at 0.025s and 0.25s respectively.

All mutants were tested on both reporter plasmids as an internal control, and all individual assays were repeated at least twice. The mean and standard deviation are shown for the elongated-hinge mutants, which followed exactly the protocol described above. The initial assays for the other mutations were carried out with a slightly different protocol (cells were switched back to glucose and then grown longer before miniprepping, etc) and therefore, while they gave very similar results, their quantification was not be averaged with that shown in Figure 7.

### Recombination and binding assays

For binding and recombination assays, protein and DNA were used in ratio 2:1 (final concentrations: DNA, 0.2 µM; protein, 0.4 µM). The reactions were performed in a buffer containing 20 mM Tris-HCl pH 8.0, 100 mM NaCl, 10 % glycerol, 50 µg/ml BSA, 0.5 mM TCEP. Samples were incubated at 37 ℃ for 1 hr.

After the incubation, the samples were divided into two sets. To assess recombination, 10 % of Sodium Dodecyl Sulfate (SDS) and 6X DNA loading dye (6X Gel loading dye, purple—New England Biolabs) were added, resulting in final concentrations of 1.1% SDS and 1X loading dye. Both sets of samples were separated on Novex 6% TBE gels (Invitrogen) in 1X TBE running buffer at 160V for 45 minutes or until the dye front reached near the bottom of the gel. The gels were visualized using a Chemidoc gel imager (Bio-Rad).

### Sample preparation for single particle cryo-EM

Purified and concentrated SPβ integrase (Int) or Int-RDF fusion proteins in storage buffer (20 mM Tris-Cl pH 8, 200 mM NaCl, 2 mM TCEP, and 20 % glycerol) were mixed with annealed mismatched or symmetrized dsDNA substrates to assemble protein-DNA complexes. These complexes contained either covalently trapped reaction intermediates with cleaved central dinucleotides or product dimer complexes with uncleaved central dinucleotides.

To generate and visualize covalently trapped intermediates, Int was mixed with *attPmm and attBmm*, whereas Int-RDF was mixed with *attLmm.* For the product dimer complexes, Int was mixed with *attLsym*, and Int-RDF was mixed with *attPsym* and *attBsym*. In all preparations, the final protein and DNA concentrations were 80 µM to 40 µM, respectively, in low salt buffer (20 mM Tris-Cl pH 8, 100mM NaCl, and 0.125mM TCEP).

Samples were incubated at 37 ℃ for 1 hour and purified using size-exclusion chromatography (SEC) on a Superose 6 10/300 GL column (Cytiva) pre-equilibrated with low salt buffer (20mM Tris-Cl pH 8, 100mM NaCl, and 0.125mM TCEP). Fractions containing the target complexes were pooled and concentrated to 5mg/ml using using Amicon Ultra Centrifugal Filter (30kDa MWCO, MilliporeSigma).

Immediately prior to blotting and vitrification, samples were adjusted to a final concentration of 2mg/ml with low salt buffer (20mM Tric-Cl pH 8, 100mM NaCl, and 0.125mM TCEP) and supplemented with fluorinated octyl maltoside (FOM, Anatrace) to a final concentration of 0.02 % (w/v). Then, 3.5 µl of the sample was applied to a freshly glow discharged CFlat 4.0/1.0 300-mesh grid or Quantifoil 1.2/1.3 300-mesh grid, blotted for 5.5 seconds with blot force set to 0, and plunged into liquid ethane using a Vitrobot^TM^ MK IV (FEI) at 18 ℃ and 100 % relative humidity.

### Data collection for single particle cryo-EM

Imaging of particles was performed using a Titan Krios G3i (Thermo Fisher Scientific) operating at 300 kV and equipped with a Gatan K3 direct detection camera. Data collection was conducted at the Advanced Electron Microscopy Facility, the University of Chicago. Each movie consisted of 54 frames with a total exposure of approximately 65 electrons per Å^2^. The defocus range was set at -0.9 µm to -1.9 µm. Additional acquisition and imaging parameters are detailed in Extended Data Table 1.

### Image processing

All data processing steps were conducted using cryoSPARC (v4.6.0)^74^ and are summarized in Figures S5, S9, S11 and S19. For both datasets, micrograph movies were aligned, summed, and dose-weighted; contrast transfer function (CTF) parameters were estimated using patch based CTF estimation from cryoSPARC. Micrographs with significant non-vitreous ice contamination or outliers in motion correction and CTF estimation were excluded.

Initial particle picking was performed using the blob particle picker to generate two-dimensional (2D) classifications. After a second round of 2D classification, selected classes showing intact complexes were used as templates for automated particle picking. False-positive or poorly defined particles were filtered out after additional rounds of 2D classification. The remaining particles were used to train a Topaz^75^ model, which were applied to the combined micrographs for refined particle picking.

Particles classes showing clear secondary structures were iteratively classified, and three to five three-dimensional (3D) ab initio models were generated with C1 or C2 symmetry. Classes representing intact synaptic complexes were selected and further refined through heterogenous and non-uniform refinement steps. Resolutions for both Int and Int-RDF datasets were improved through focused refinement steps, where particles from both pre- and post-rotated classes were combined, resulting in the alignment of the half complex. Reference-based motion correction was applied, followed by particle subtraction to remove the unaligned part of the other half complex, and the final set of particles were used for local refinement.

To refine flexible regions of the complexes, built-in 3D variability analysis^76^ and 3D flex reconstruction^77^ from cryoSPARC^74^ were performed to address disordered regions and deconvolute the heterogeneous conformations of the complex into top three components ranked by significance in conformational variability. Using the outputs of these two analytical programs, we generated movies (see supplementary movies) depicting the various conformational variabilities in the consensus maps and in the focused maps within UCSF Chimera (v.1.17.3)^78^. Four maps from the last stage of the non-uniform refinement cycle, corresponding to pre- and post-rotated states in both the Int and Int-RDF datasets, were used as representative consensus maps. Two focused refined half-complex maps were then combined using the corresponding consensus map as a guide to generate composite maps for these rotational states. The final maps were sharpened using cryoSPARC’s built-in map sharpening tools, and the composite maps for pre- and post- rotated states were further processed with deepEMhancer (v.0.14)^79^.

### Model building, refinement and validation

Alphafold2-multimer^80,81^ was used to create the starting model of the synaptic complex. Specifically, the complex was predicted in overlapping fragments to allow flexibility in relative domain arrangements. These fragments were subsequently positioned and aligned according to the cryo-EM map using ChimeraX (v.1.4)^82^. Overlapping regions were manually corrected, adjusted, and refined in COOT (v.0.9.8.8)^83^. All of the DNA could be modeled except for the 3’ most of the two overhanging Ts created by the DNA cleavage reaction. While most protein residues were built, some regions containing flexible loops, the 20aa long N-terminal tag, 28aa C-terminal residues and the linker connecting the Int and RDF were not built because they were disordered in the maps. The register of the main chain was carefully checked and corrected based on bulky residues.

B-form DNA starting models for the DNA substrates were built in ChimeraX and fitted into the cryo-EM map with local restraints using COOT^83^. ISOLDE (v.1.4)^84^, embedded in UCSF ChimeraX^82^, was used to introduce phosphoserine linkages between the protein and DNA, and to correct most of peptide bond geometry, rotamer, and Ramachandran outliers.

Manual model building and refinement were performed iteratively using COOT^83^, ISOLDE^84^, and PHENIX (v.1.21)^85^ real-space refinement. Both unsharpened and sharpened maps, as well as 3D-flex reconstructed maps, were used in conjunction with previously solved truncated structures and Alphafold2^80,81^ predicted models to guide the model building process. Geometry restraints for zinc-binding sites were applied based on a zinc-binding model to ensure accurate representation of these regions.

Molecular graphics and structural analyses were conducted using UCSF ChimeraX^82^ and PyMOL (v. 3.1.3)^86^.

---

**Extended Data Table 1.**
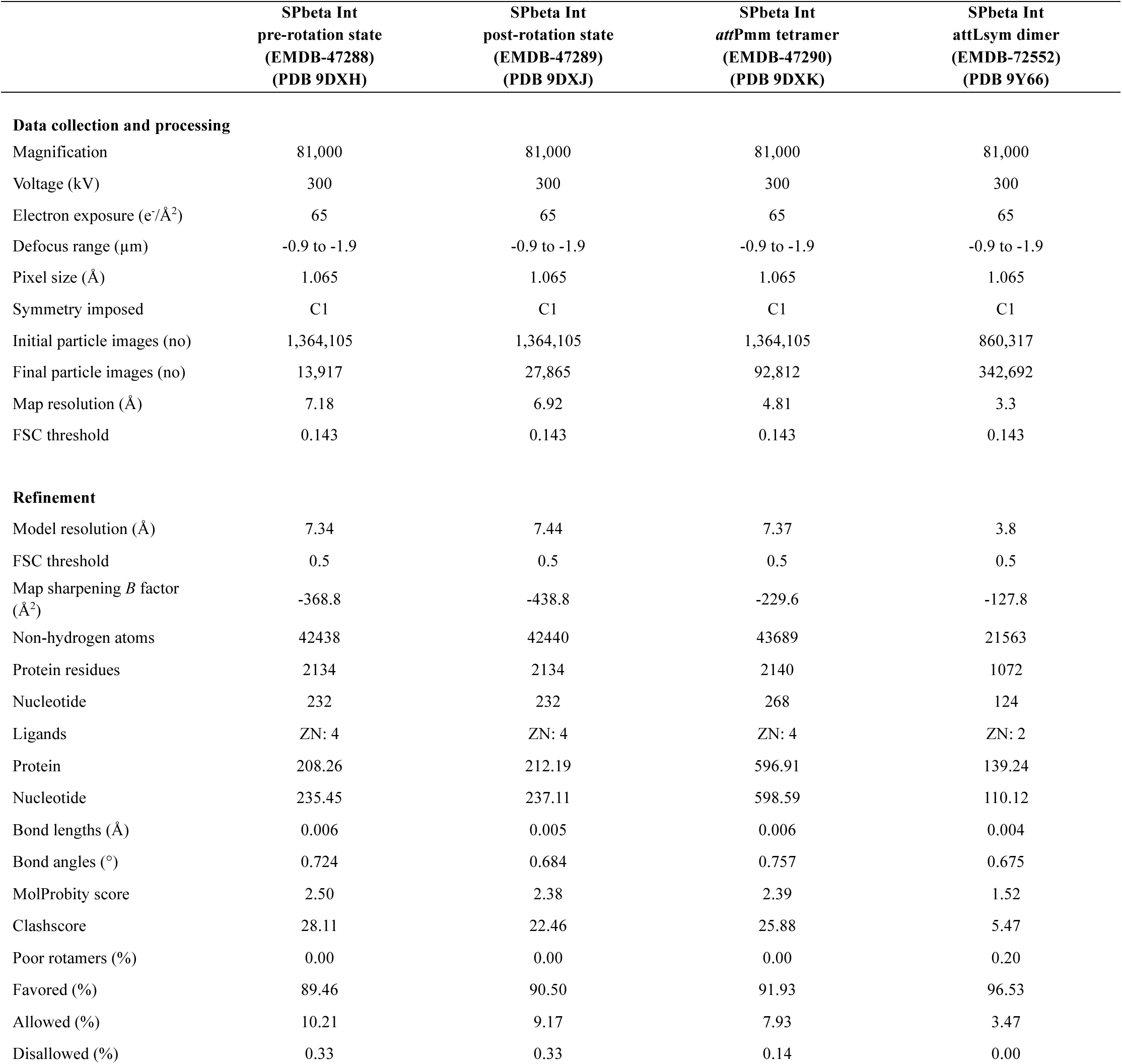
Cryo-EM Data Collection, Refinement and Validation Statistics.

|  | SPbeta Int<br>pre-rotation state<br>(EMDB-47288)<br>(PDB 9DXH) | SPbeta Int<br>post-rotation state<br>(EMDB-47289)<br>(PDB 9DXJ) | SPbeta Int<br>*attPmm* tetramer<br>(EMDB-47290)<br>(PDB 9DXK) | SPbeta Int<br>*attLsym* dimer<br>(EMDB-72552)<br>(PDB 9Y66) |
| --- | --- | --- | --- | --- |
| **Data collection and processing** |  |  |  |  |
| Magnification | 81,000 | 81,000 | 81,000 | 81,000 |
| Voltage (kV) | 300 | 300 | 300 | 300 |
| Electron exposure (e<sup>-</sup>/Å<sup>2</sup>) | 65 | 65 | 65 | 65 |
| Defocus range (μm) | -0.9 to -1.9 | -0.9 to -1.9 | -0.9 to -1.9 | -0.9 to -1.9 |
| Pixel size (Å) | 1.065 | 1.065 | 1.065 | 1.065 |
| Symmetry imposed | C1 | C1 | C1 | C1 |
| Initial particle images (no) | 1,364,105 | 1,364,105 | 1,364,105 | 860,317 |
| Final particle images (no) | 13,917 | 27,865 | 92,812 | 342,692 |
| Map resolution (Å) | 7.18 | 6.92 | 4.81 | 3.3 |
| FSC threshold | 0.143 | 0.143 | 0.143 | 0.143 |
| **Refinement** |  |  |  |  |
| Model resolution (Å) | 7.34 | 7.44 | 7.37 | 3.8 |
| FSC threshold | 0.5 | 0.5 | 0.5 | 0.5 |
| Map sharpening *B* factor<br>(Å<sup>2</sup>) | -368.8 | -438.8 | -229.6 | -127.8 |
| Non-hydrogen atoms | 42438 | 42440 | 43689 | 21563 |
| Protein residues | 2134 | 2134 | 2140 | 1072 |
| Nucleotide | 232 | 232 | 268 | 124 |
| Ligands | ZN: 4 | ZN: 4 | ZN: 4 | ZN: 2 |
| Protein | 208.26 | 212.19 | 596.91 | 139.24 |
| Nucleotide | 235.45 | 237.11 | 598.59 | 110.12 |
| Bond lengths (Å) | 0.006 | 0.005 | 0.006 | 0.004 |
| Bond angles (°) | 0.724 | 0.684 | 0.757 | 0.675 |
| MolProbity score | 2.50 | 2.38 | 2.39 | 1.52 |
| Clashscore | 28.11 | 22.46 | 25.88 | 5.47 |
| Poor rotamers (%) | 0.00 | 0.00 | 0.00 | 0.20 |
| Favored (%) | 89.46 | 90.50 | 91.93 | 96.53 |
| Allowed (%) | 10.21 | 9.17 | 7.93 | 3.47 |
| Disallowed (%) | 0.33 | 0.33 | 0.14 | 0.00 |

**Extended Data Table 2.** Cryo-EM Data Collection, Refinement and Validation Statistics.

|  | SPbeta Int RDF<br>pre-rotation state<br>(EMDB-47284)<br>(PDB 9DXD) | SPbeta Int RDF<br>post-rotation state<br>(EMDB-47286)<br>(PDB 9DXF) | SPbeta Int RDF<br>*attPmm* dimer<br>(EMDB-47287)<br>(PDB 9DXG) | SPbeta Int RDF<br>*attPsym* dimer<br>(EMDB-47284)<br>(PDB 9Y6V) |
| --- | --- | --- | --- | --- |
| **Data collection and processing** |  |  |  |  |
| Magnification | 81,000 | 81,000 | 81,000 | 81,000 |
| Voltage (kV) | 300 | 300 | 300 | 300 |
| Electron exposure (e<sup>-</sup>/Å<sup>2</sup>) | 65 | 65 | 65 | 65 |
| Defocus range (μm) | -0.9 to -1.9 | -0.9 to -1.9 | -0.9 to -1.9 | -0.9 to -1.9 |
| Pixel size (Å) | 1.065 | 1.065 | 1.065 | 1.065 |
| Symmetry imposed | C1 | C1 | C1 | C1 |
| Initial particle images (no) | 2,114,654 | 2,114,654 | 2,114,654 | 162,334 |
| Final particle images (no) | 550,169 | 248,916 | 54,766 | 24,616 |
| Map resolution (Å) | 4.82 | 5.00 | 6.61 | 6.9 |
| FSC threshold | 0.143 | 0.143 | 0.143 | 0.143 |
| **Refinement** |  |  |  |  |
| Model resolution (Å) | 4.23 | 4.22 | 12.70 | 7.10 |
| FSC threshold | 0.5 | 0.5 | 0.5 | 0.5 |
| Map sharpening *B* factor<br>(Å<sup>2</sup>) | -321.1 | -395.5 | -505.3 | -620.6 |
| Non-hydrogen atoms | 46399 | 46360 | 23758 | 23118 |
| Protein residues | 2366 | 2364 | 1182 | 1182 |
| Nucleotide | 232 | 232 | 134 | 114 |
| Ligands | ZN: 4 | ZN: 4 | ZN: 2 | ZN: 2 |
| Protein | 72.31 | 71.31 | 1333.16 | 617.12 |
| Nucleotide | 68.69 | 68.69 | 71.64 | 523.94 |
| Bond lengths (Å) | 0.005 | 0.005 | 0.004 | 0.003 |
| Bond angles (°) | 0.633 | 0.633 | 0.757 | 0.594 |
| MolProbity score | 1.86 | 1.90 | 2.36 | 2.23 |
| Clashscore | 7.96 | 8.63 | 22.42 | 14.07 |
| Poor rotamers (%) | 0.00 | 0.00 | 0.00 | 1.94 |
| Favored (%) | 93.39 | 93.23 | 91.23 | 94.89 |
| Allowed (%) | 6.41 | 6.69 | 8.77 | 5.11 |
| Disallowed (%) | 0.00 | 0.09 | 0.00 | 0.00 |

**Supplementary figure 1.**
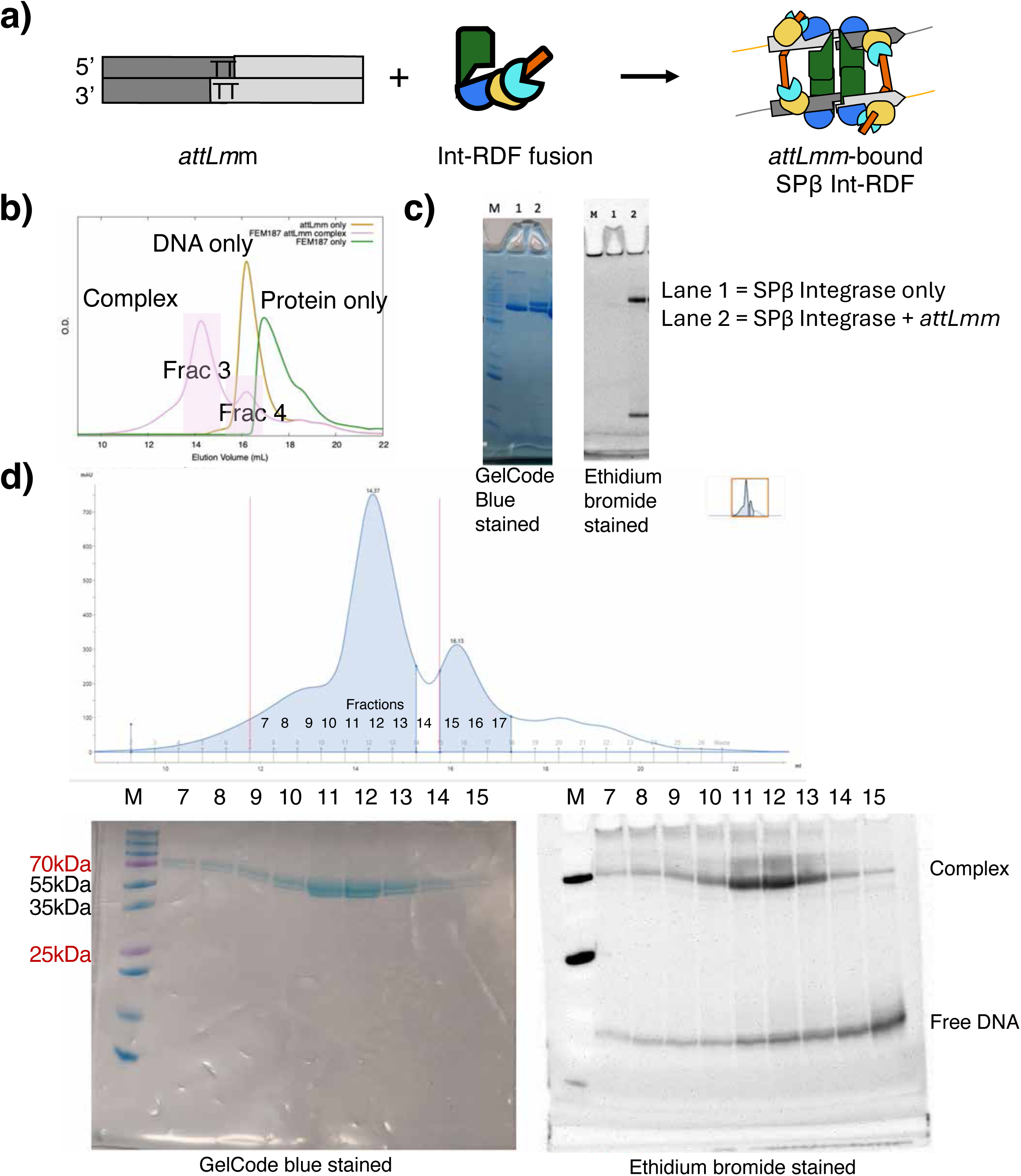
Purification of covalently trapped SPβ Int-RDF fusion – *attLmm* complex used for cryoEM. A. Schematic diagram of covalently trapped SPβ Int-RDF and *attLmm* synaptic complex. B. Size exclusion chromatography (SEC) elution profile of *attLmm* bound SPβ Int-RDF fusion, protein alone, and DNA alone. C. SDS-PAGE of SPβ Int-RDF fusion alone and *attLmm*-bound complex visualized on an SDS gel with GelCode blue staining (on the left) and ethidium bromide staining (on the right). D. Elution profile of *attLmm* bound SPβ Int-RDF from SEC and SDS gel electrophoresis of the fractions (stained with GelCode blue, left and ethidium bromide, right). Fractions 11 and 12 were pooled and applied to EM grids.

**Supplementary figure 2.**
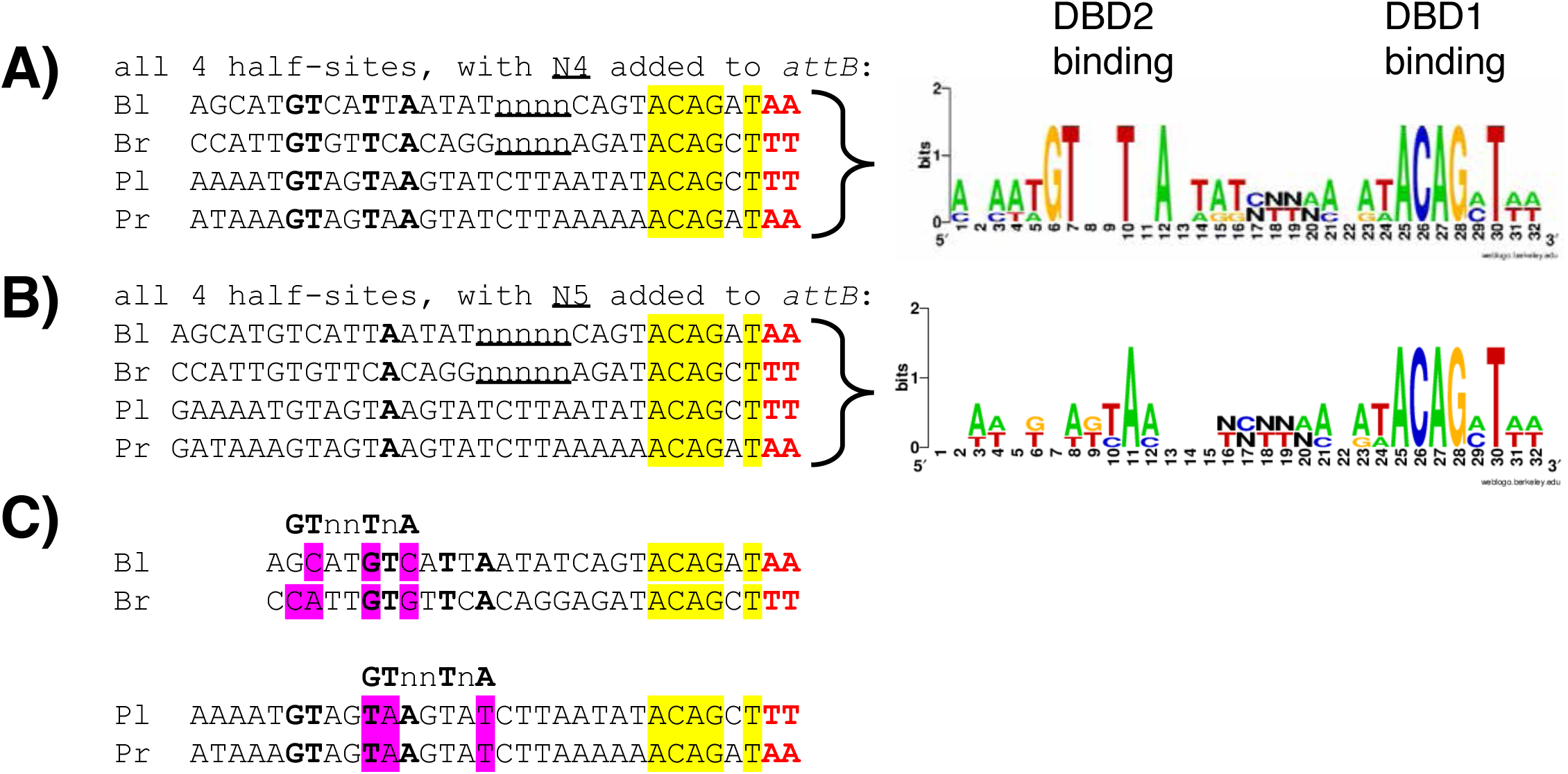
Differentiation of the two types of *att* site. A. Alignment of all 4 natural half-sites (oriented 5’ to 3’), assuming a 4-bp offset for the DBD2-specific motif between B- and P- type half sites, as supported by the experimental maps. The *att* sites for phage SPβ were taken from the genome sequence of *B. subtilis* (NCBI Accession: NZ_CP051860.2). Bl and Pl are the two halves that combine to form *attL*; likewise Br and Pr comprise *attR*. The central dinucleotide is highlighted in red. Nucleotides that are identical in all 4 half-sites in the DBD1-binding region are highlighted in yellow, while those in the DBD2-binding region are shown in bold. A sequence logo (https://weblogo.berkeley.edu/logo.cgi)^87^ for these alignments is shown on the right. B. A similar alignment of SPβ half-sites, but with N5 added between motifs, as may be more common among other large serine integrases^5,54^. C. *attP* and *attB* are distinguishable from one another not only by the presence of conserved bases in the correct positions but by their absence in the incorrect positions. The 4 half sites are shown, with conserved bases denoted as in part (A). Above each pair is the DBD2-specific consensus, shifted as it would appear in the opposite type of half-site. Bases that do not match the shifted consensus are highlighted in pink. For example, *attB* sites do not have a GT sequence in the position where the conserved GT motif is found in *attP* sites, and *attP* sites do not have a GT motif in the position where it is found in *attB* sites.

**Supplementary figure 3.**
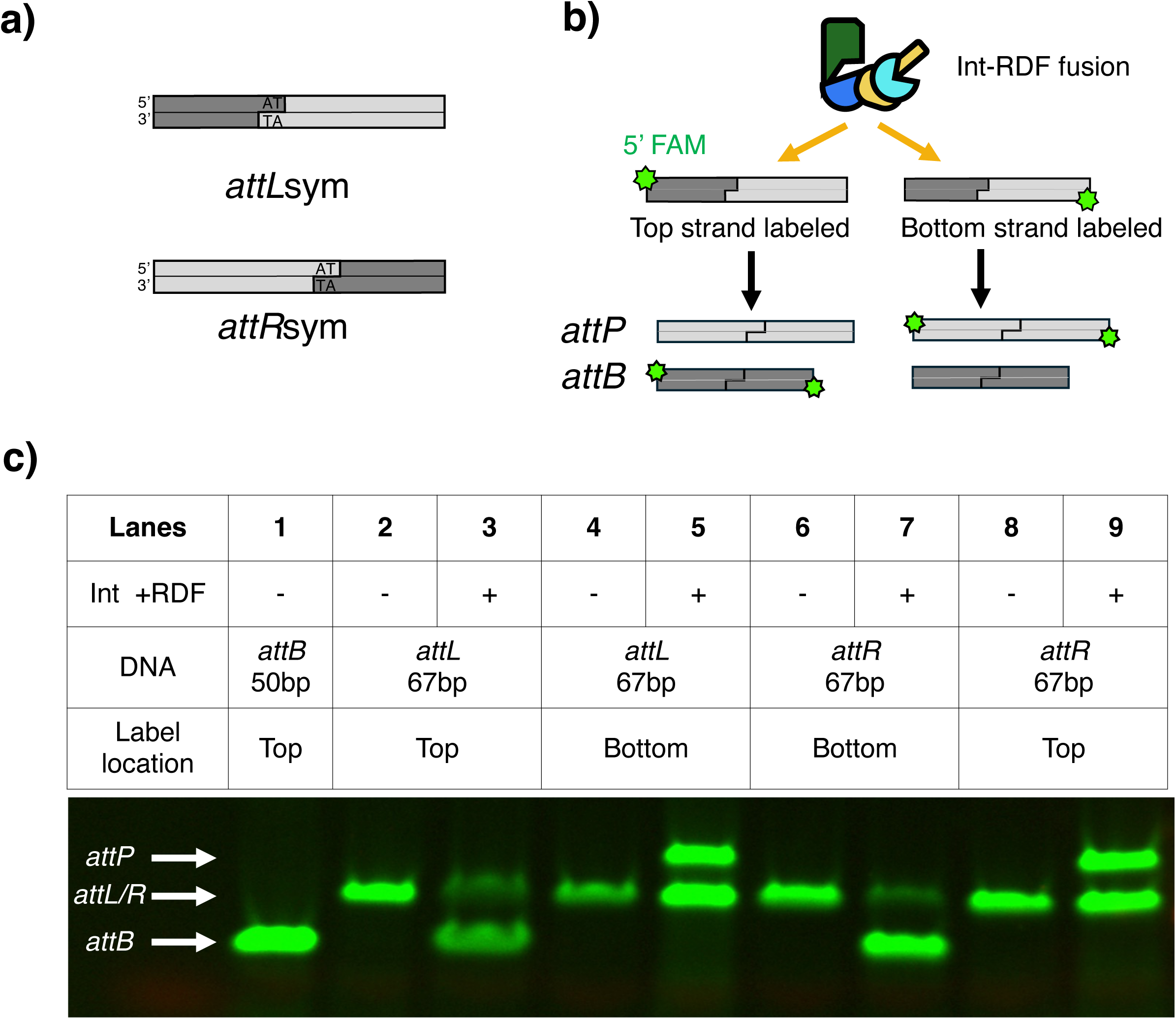
The SPβ Int-RDF fusion protein recombines *attLsym* and *attRsym* substrates with comparable efficiency. A) The *attLsym* and *attRsym* substrates (67 bp) carried the WT *attL* and *attR* sequences except that an extension was added to the *attP* side and the central dinucleotide (naturally TT) was symmetrized (now AT) to allow productive recombination between two copies of *attL* or two copies of *attR*. B) For each DNA substrate, the 5’ end of either the top or bottom strand is labeled with fluorescein (marked with a green star). The Int-RDF fusion protein mediates the recombination of *attLsym* × *attLsym* or *attRsym* × *attRsym* substrates to generate rearranged *attB*s (50 bp) and *attP*s (84 bp) products. C) Fluorescently labeled recombinant products are visualized on the gel, enabling the detection of one of the two recombination products.

**Supplementary figure 4.**
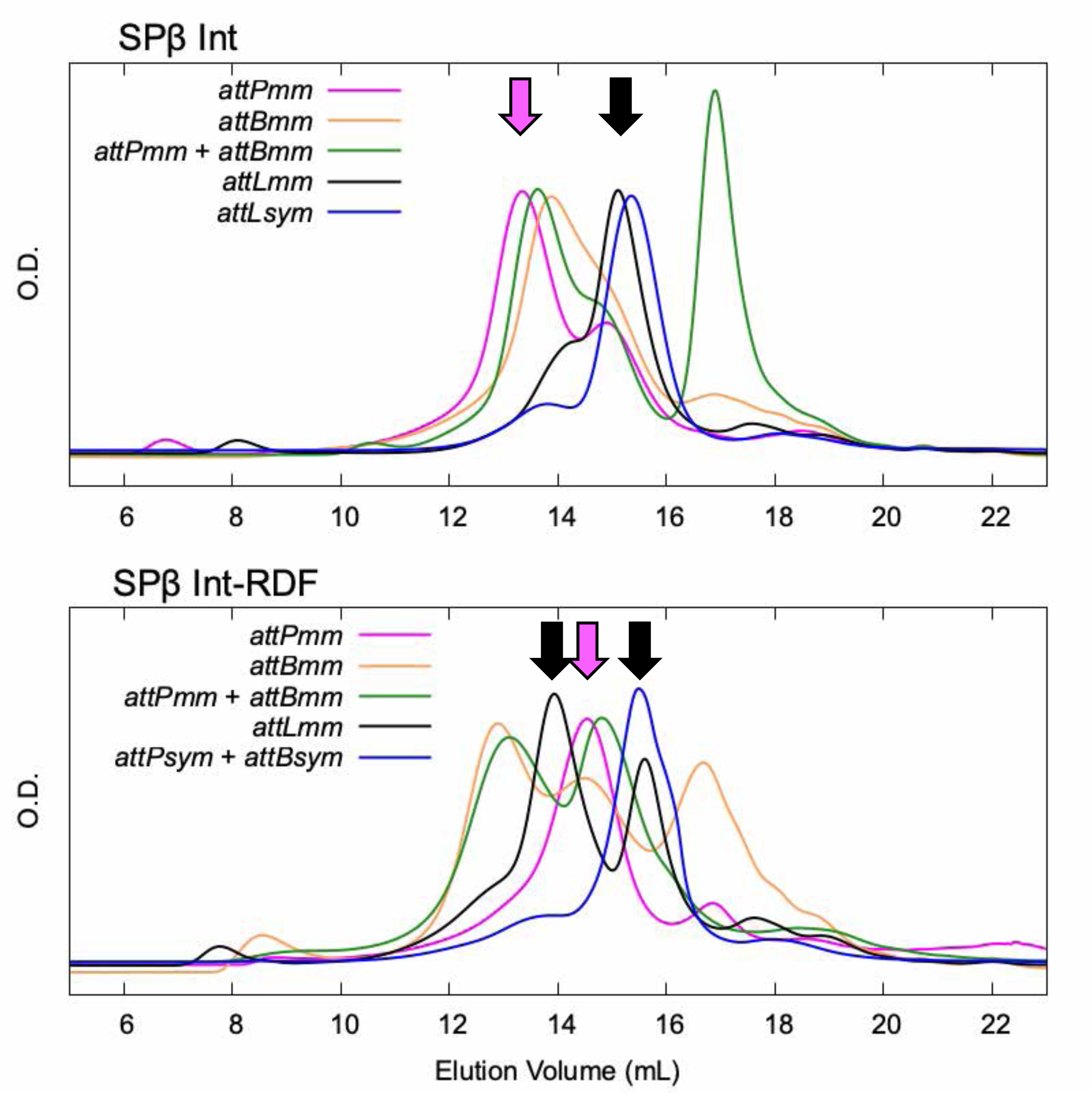
Elution profiles of various *att* site—SPβ (±RDF) complexes analyzed by size exclusion chromatography (SEC). Elution profiles of complexes formed with various *att* sites were analyzed via SEC. Top panel: SPβ integrase complexes in the absence of the recombination directionality factor (RDF). Bottom panel: SPβ integrase complexes in the presence of RDF. Tetramers synapsing two copies of *attPmm* were only seen in the absence of the RDF (pink arrows); When SPβ Int-RDF complexes were mixed with *attLmm*, two peaks were observed (black arrows), corresponding to: (1) covalently trapped *attLmm*—SPβ int-RDF tetramer complex, and (2) covalently trapped *attLmm*-derived *attPmm* dimer complex.

**Supplementary figure 5.**
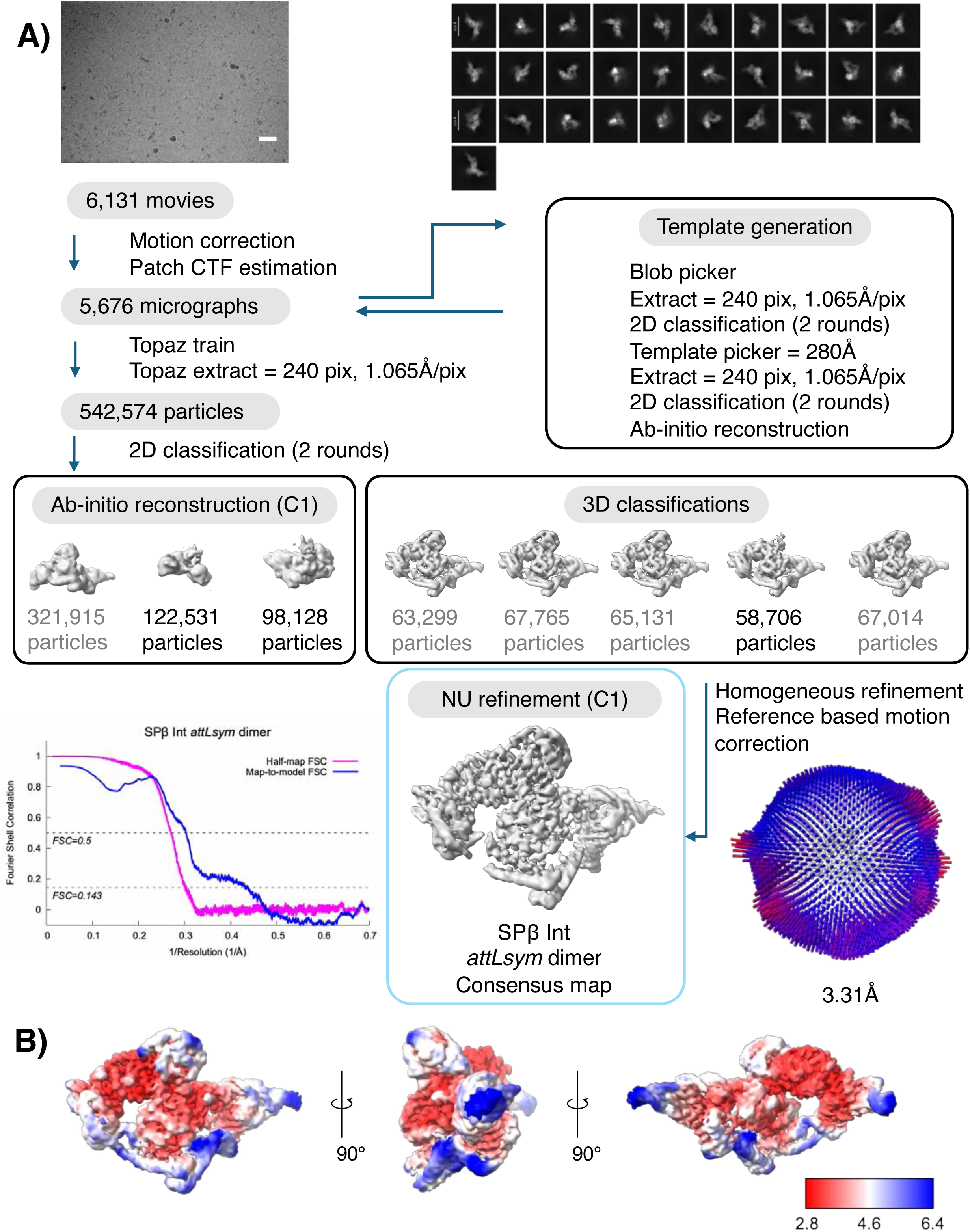
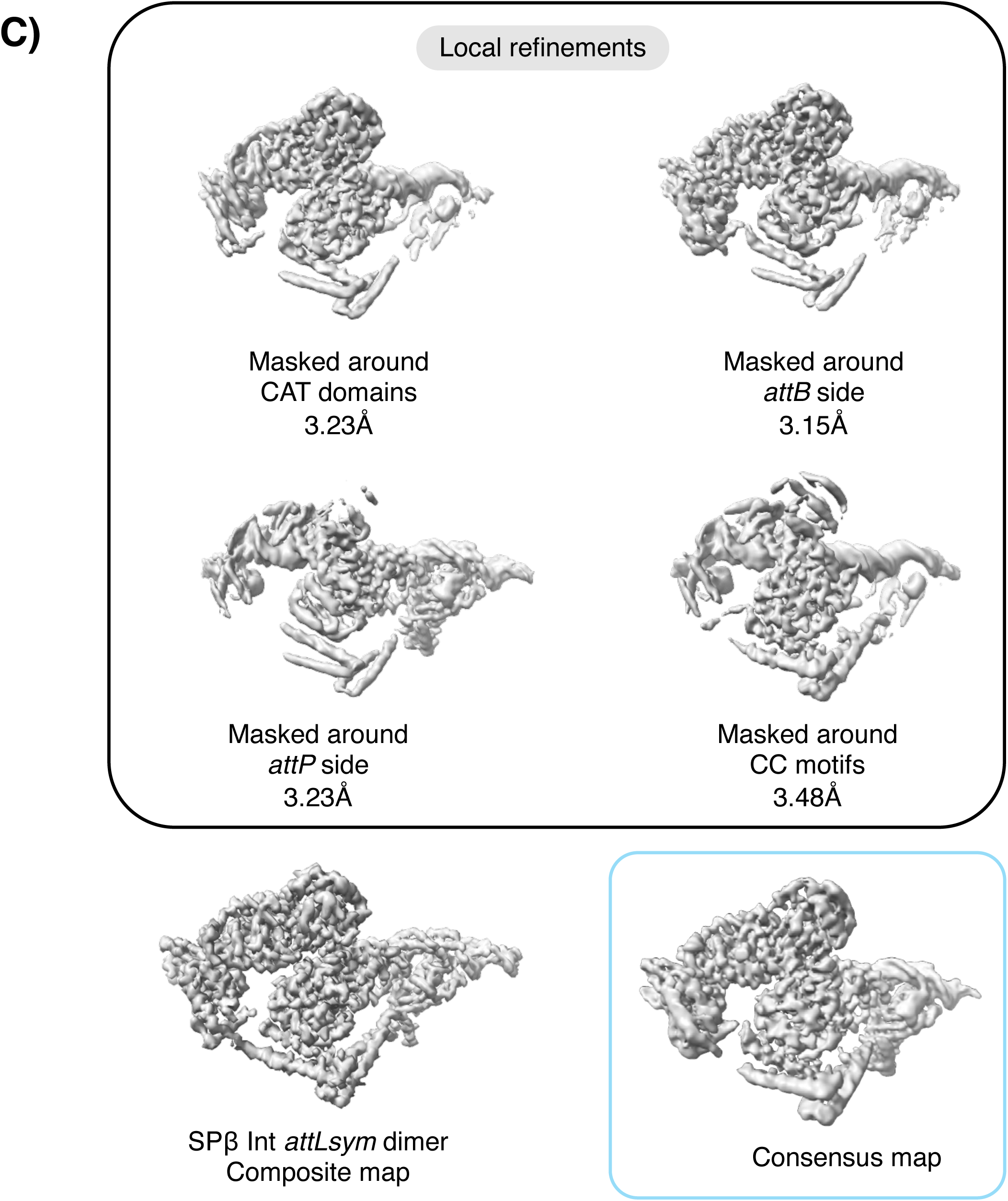
Data processing flow chart for the SPβ Int *attLsym* data. A. A visual overview of the data processing steps of SPβ Int *attLsym* dataset. The dataset was subjected to particle selection, iterative rounds of 2D classification and 3D classification. A representative micrograph (scale bar, 40nm) and representative 2D class averages are shown. The distribution of the Euler angles is shown next to the consensus map. Fourier shell correlation (FSC) curve of the masked map after cryoSPARC^74^ postprocessing. The resolution was determined by the FSC=0.143 (magenta). The map-to-model FSC curve is also shown (blue). B. Local resolution of the map was calculated using cryoSPARC^74^. C. The final maps include one consensus map outlined in light blue, four local refined maps, and a composite map.

**Supplementary figure 6.**
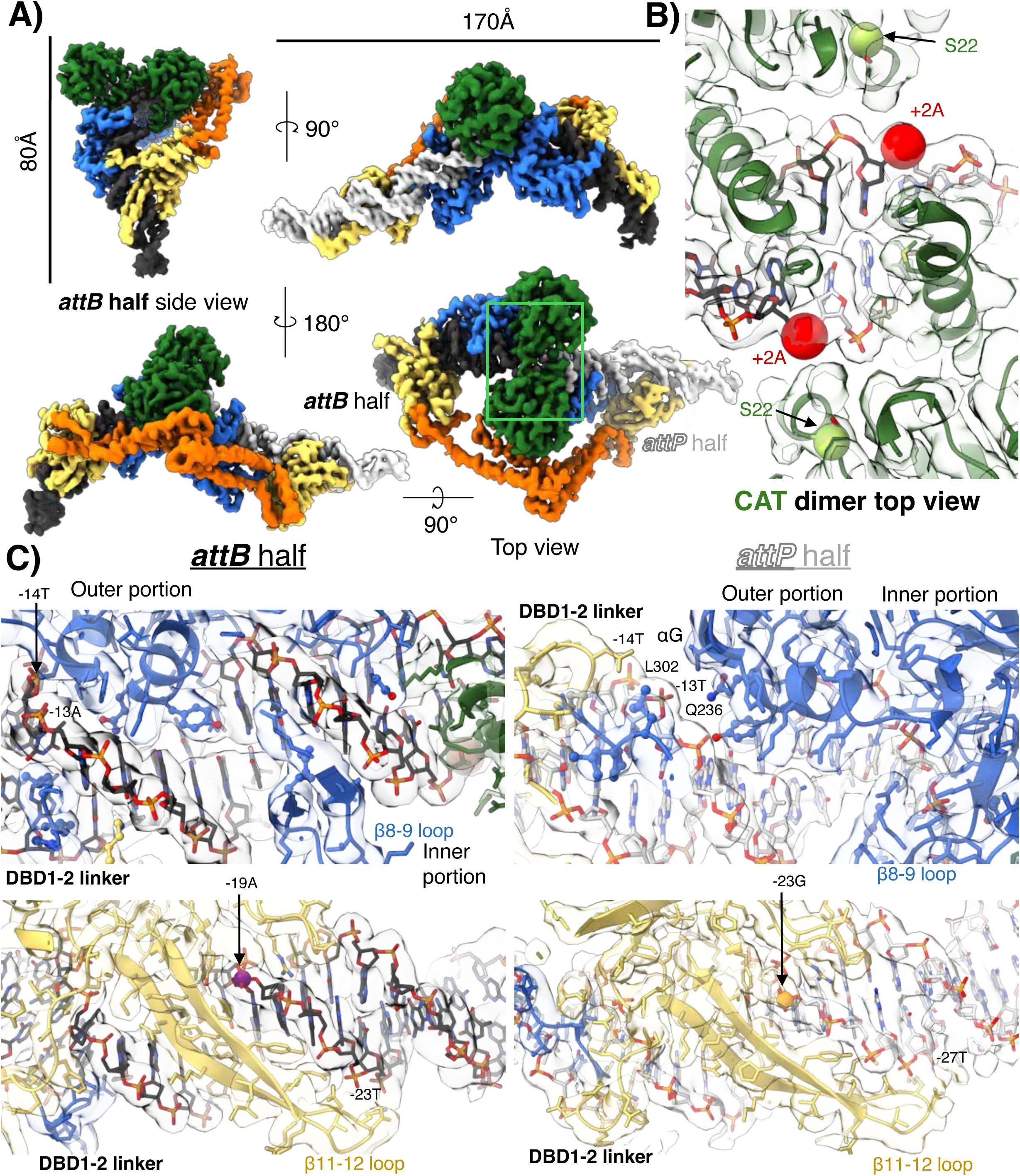
Density maps for the *attLsym* bound SPβ Integrase dimer. A. Multiple views of the composite cryo-EM map, colored as in Figure 1. B. A top view of the CAT dimer, indicated by a green box in panel A, is shown. S22 residues are displayed as yellow-green spheres, and the scissile phosphates are shown as red spheres. The density map is superimposed in transparent gray. C. Interactions of DBD1 (top row) and DBD2 (bottom row) with *attB*-half (left column) and *attP*-half sites (right column) are shown in the same orientations as in Figure 2, but with all side chains shown (those in Figure 2 and discussed in the text with spheres) and the density map is superimposed on the model.

**Supplementary figure 7.**
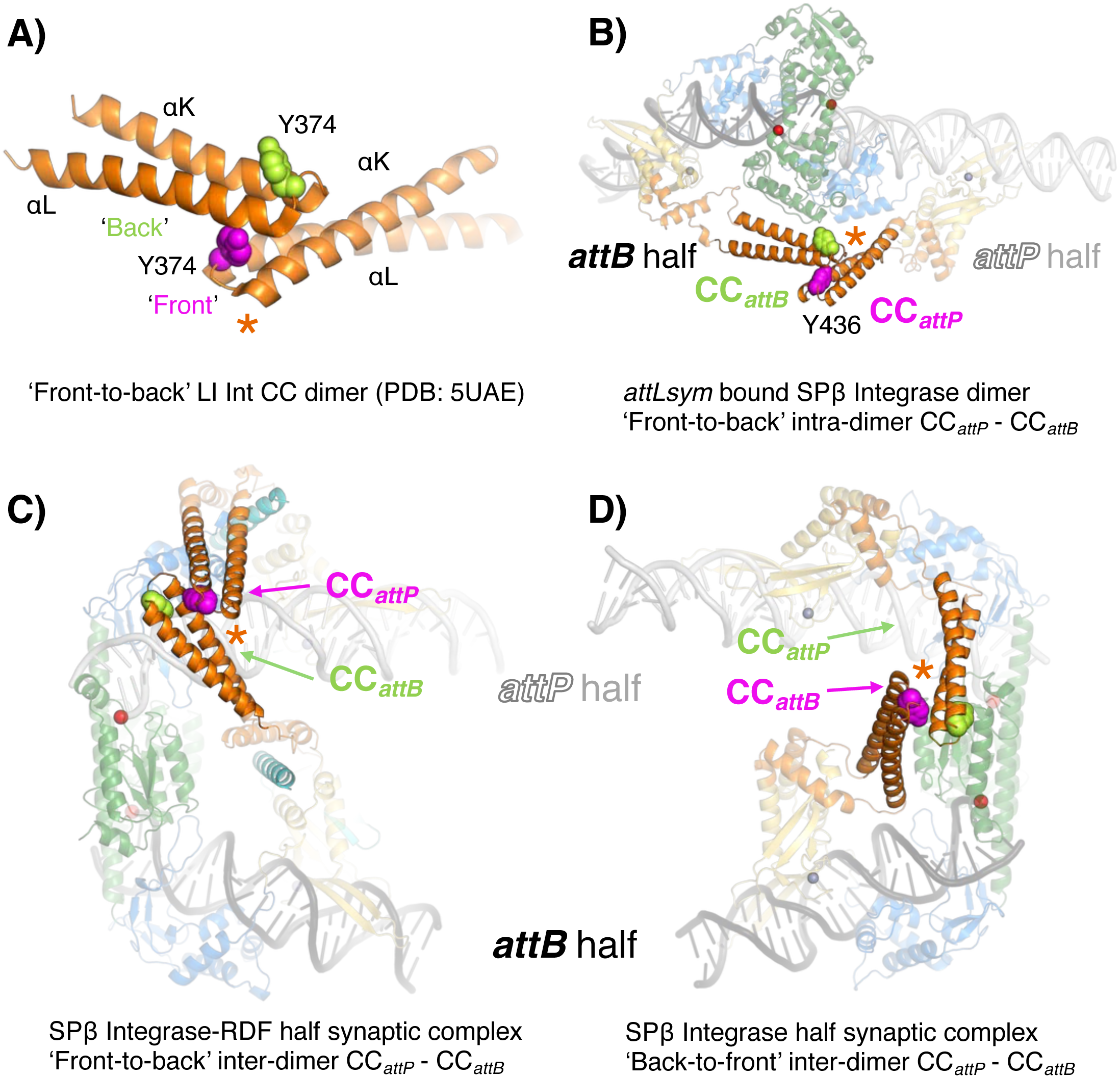
Front-to-back modes of CC-CC interactions. A. Structures of the isolated LI Integrase CC, residues 345-405; PDB: 5UAE)^2^. Y374, located at the tip of the αK-αL loop, is shown as spheres. The “front” side of one CC (magenta Y374) interacts with the “back” side of the other CC (with Y374 highlighted in light green). B. Structure of the SPβ integrase *attLsym* dimeric complex highlighting the CC-CC dimer, in which front side of the *attP-*bound subunit’s CC interacts with the back side of the *attB-*bound subunit’s CC. The structure is aligned according to the CCs shown in panel A. C. Structures of the SPβ integrase synaptic half complex in the presence (left) and absence (right) of RDF, illustrating the orientations of the CC*_attP_*-CC*_attB_* interactions adopted in synaptic complexes. The coloring scheme follows panel A.

**Supplementary figure 8.**
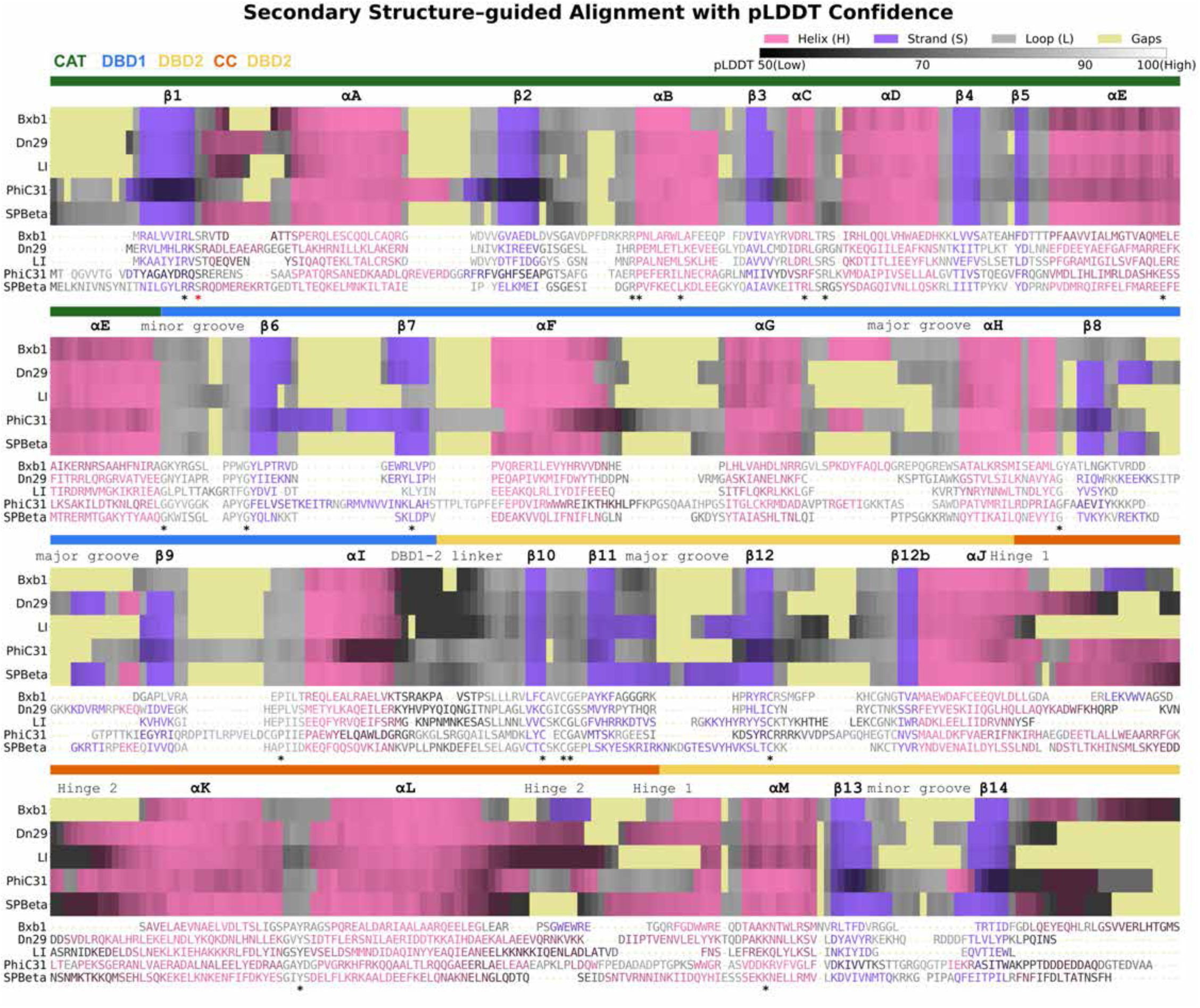
Secondary structure-guided sequence alignments. The structure of SPβ was aligned with four AlphaFold2^80^ predicted large serine integrases. Secondary structure assignments are color coded (helix-pink; strand-purple; loop-gray; gaps-yellow). Color value (intensity), scaled from 50 to100, reflects the pLDDT score from low to high confidence. The bar above the alignment is colored according to the protein domains and asterisks mark residues that are identical in all 5 LSIs.

**Supplementary figure 9.**
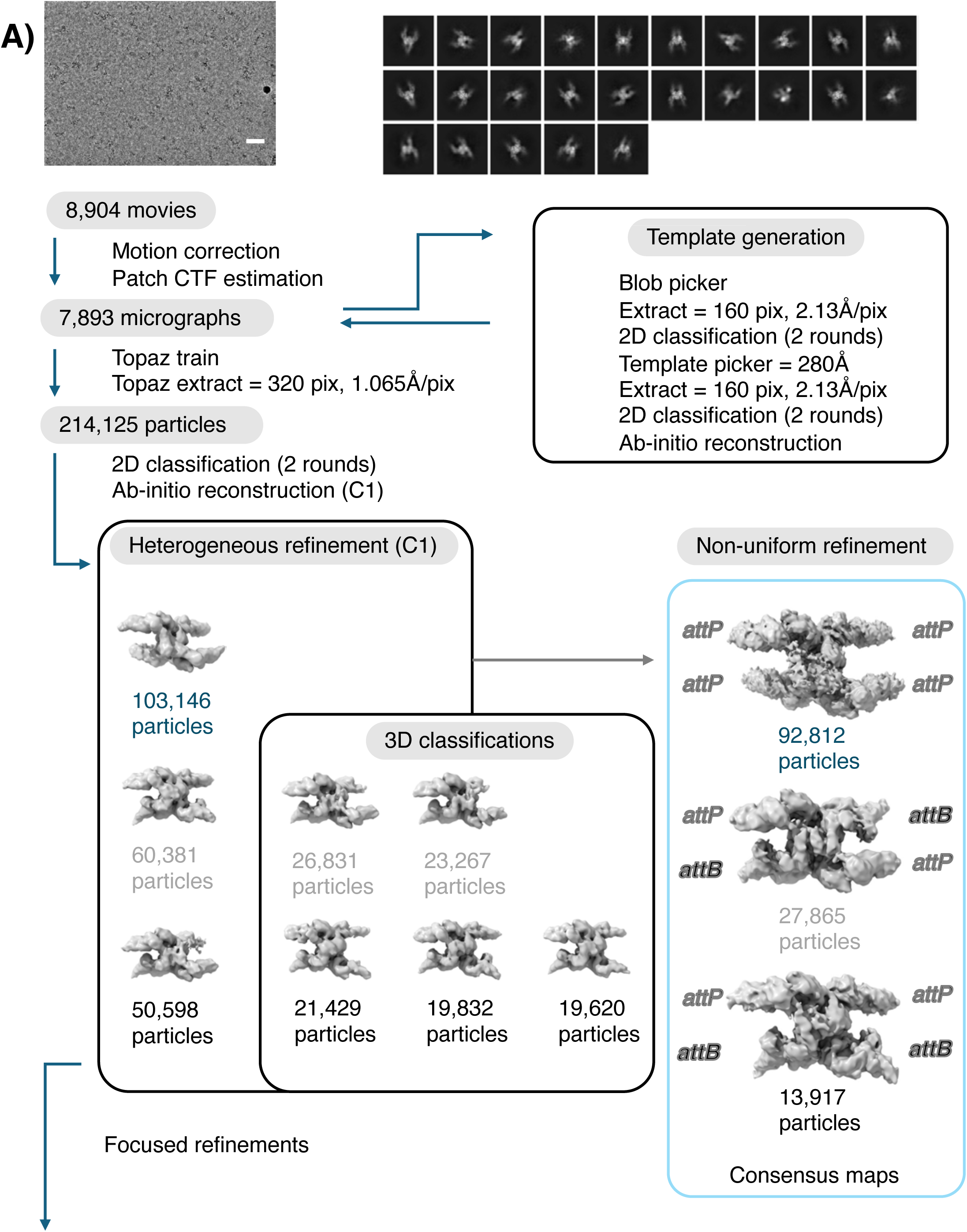
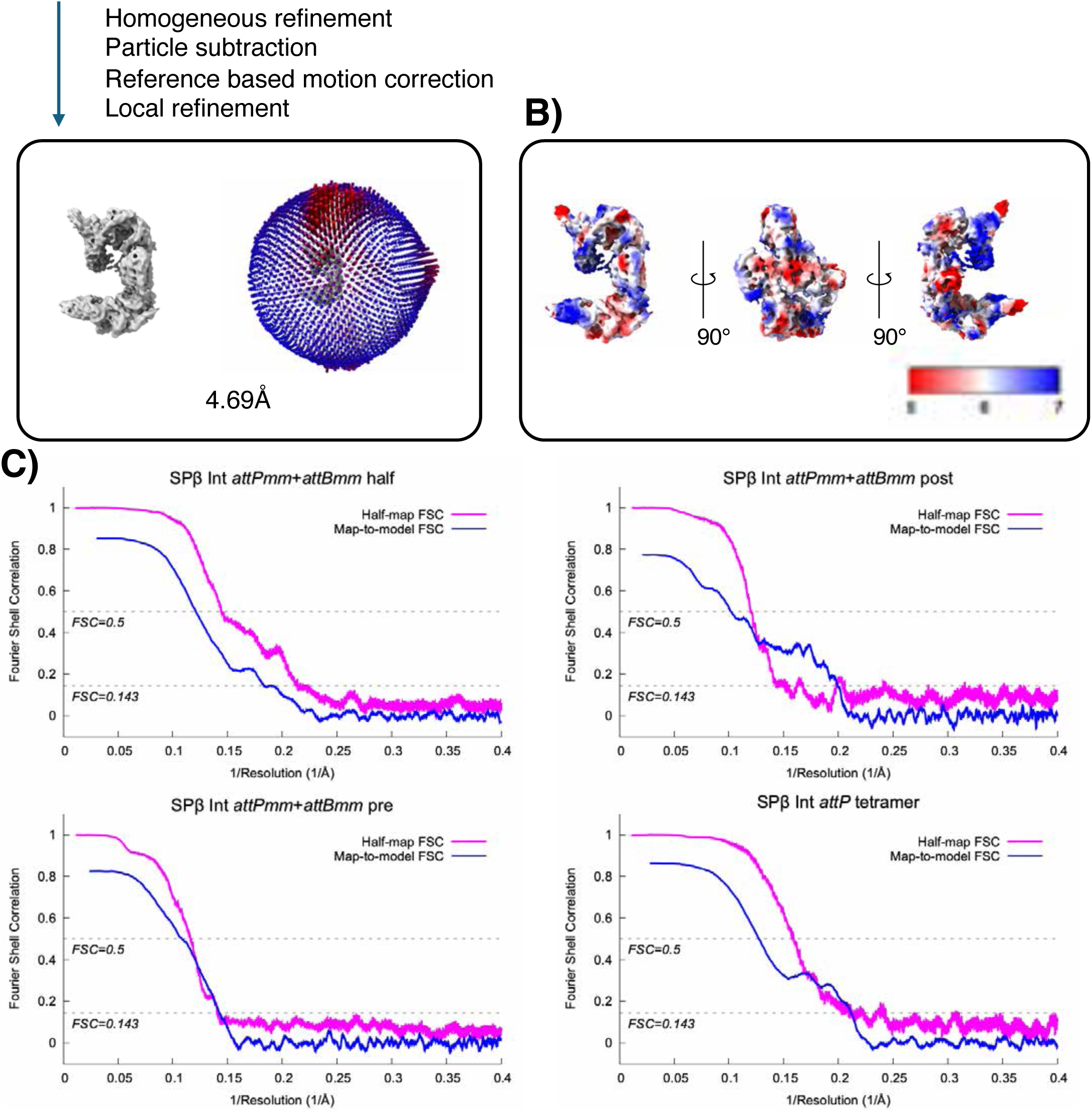
Data processing flow chart for the SPβ Int *attPmm* and *attBmm* data. A. A visual overview of the data processing steps for the SPβ Int *attPmm* and *attBmm* dataset. The dataset was subjected to particle selection, iterative rounds of 2D classification and 3D classification. A representative micrograph (scale bar, 40nm) and representative 2D class averages are shown. The distribution of Euler angles is displayed next to the half complex map. B. Local resolution of the map was calculated using cryoSPARC^74^. C. Fourier shell correlation (FSC) curve of the masked map after cryoSPARC^74^ postprocessing. The resolution was determined by the FSC=0.143 (magenta). The map-to-model FSC curve is also shown (blue).

**Supplementary figure 10.**
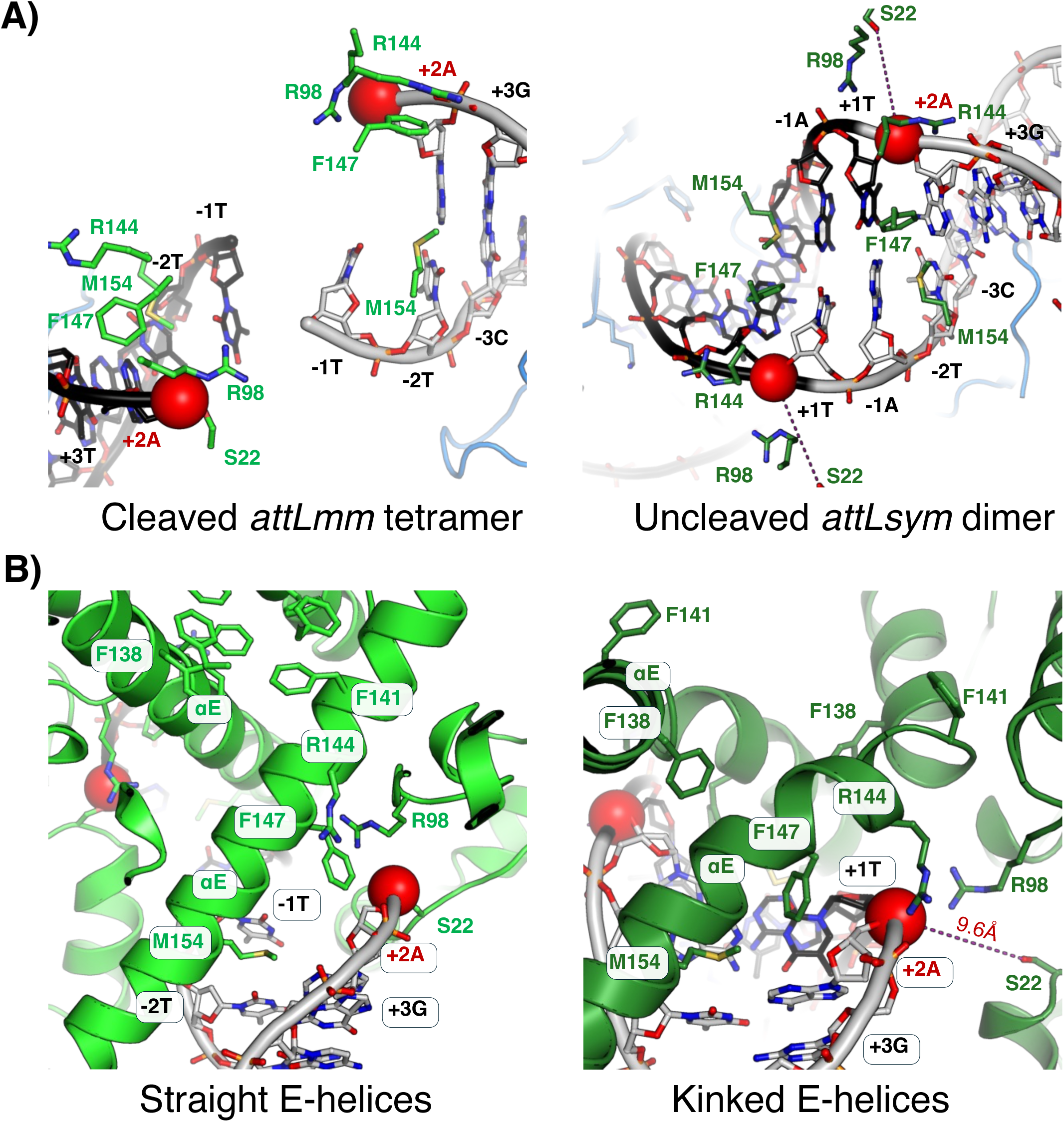
Additional views of the interactions of the E-helices at the DNA cleavage site. A. Key residues forming close atomic contacts with DNA at the cleavage sites in the cleaved *attLmm* (left) and the uncleaved *attLsym* (right) states. In the catalytically trapped cleaved *attLmm* synaptic complex, a phosphoserine linkage is formed between the oxygen atom of Ser22 and the phosphorous atom of dA2, highlighted as a red sphere. (Note that the +1 nucleotide of each chain was too poorly ordered to be modeled in the cleaved complexes). B. Ribbon diagrams of the two states illustrate different molecular interactions between the cleaved *attLmm* bound synaptic and uncleaved *attLsym* bound dimeric complexes.

**Supplementary figure 11.**
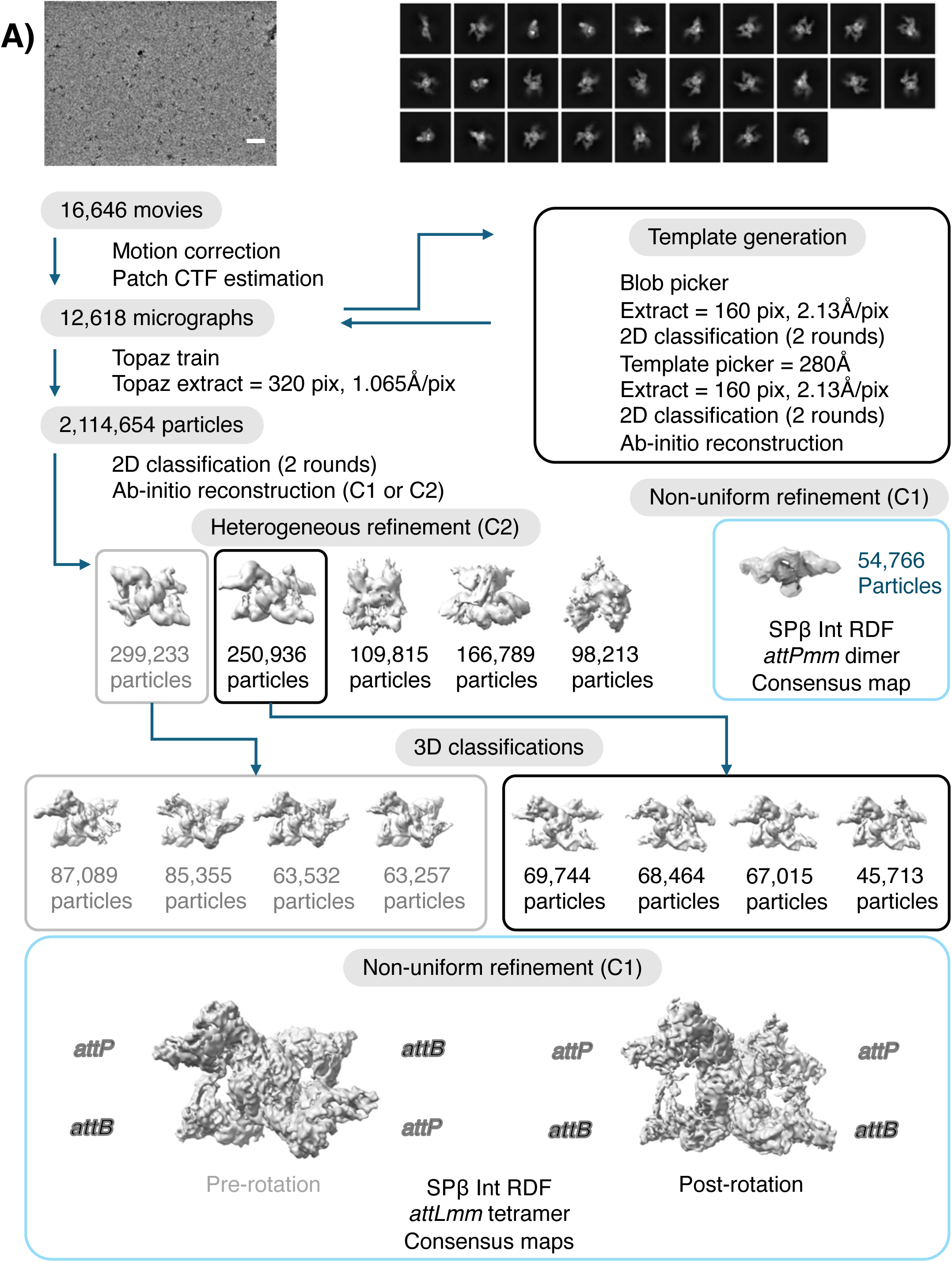
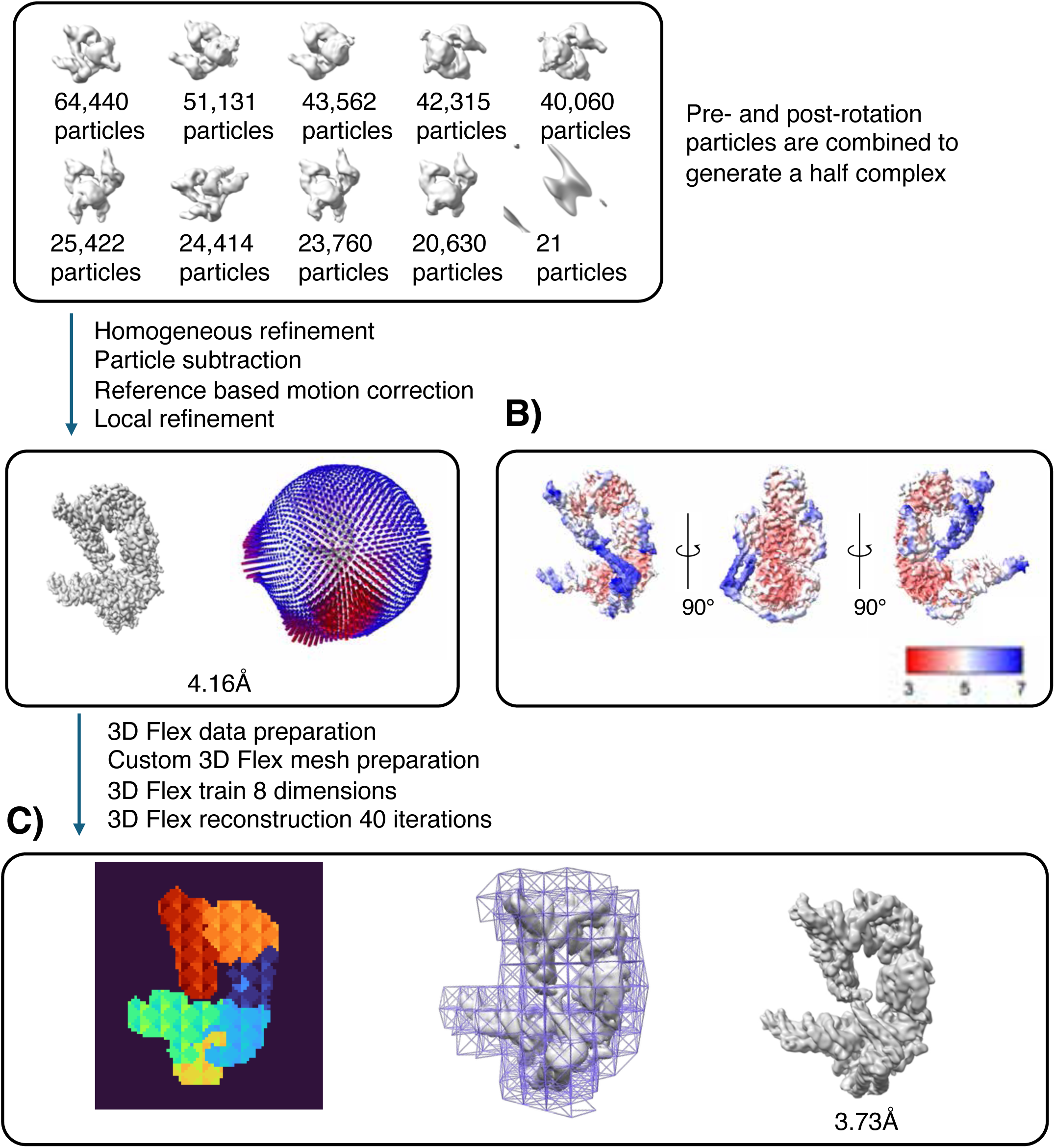
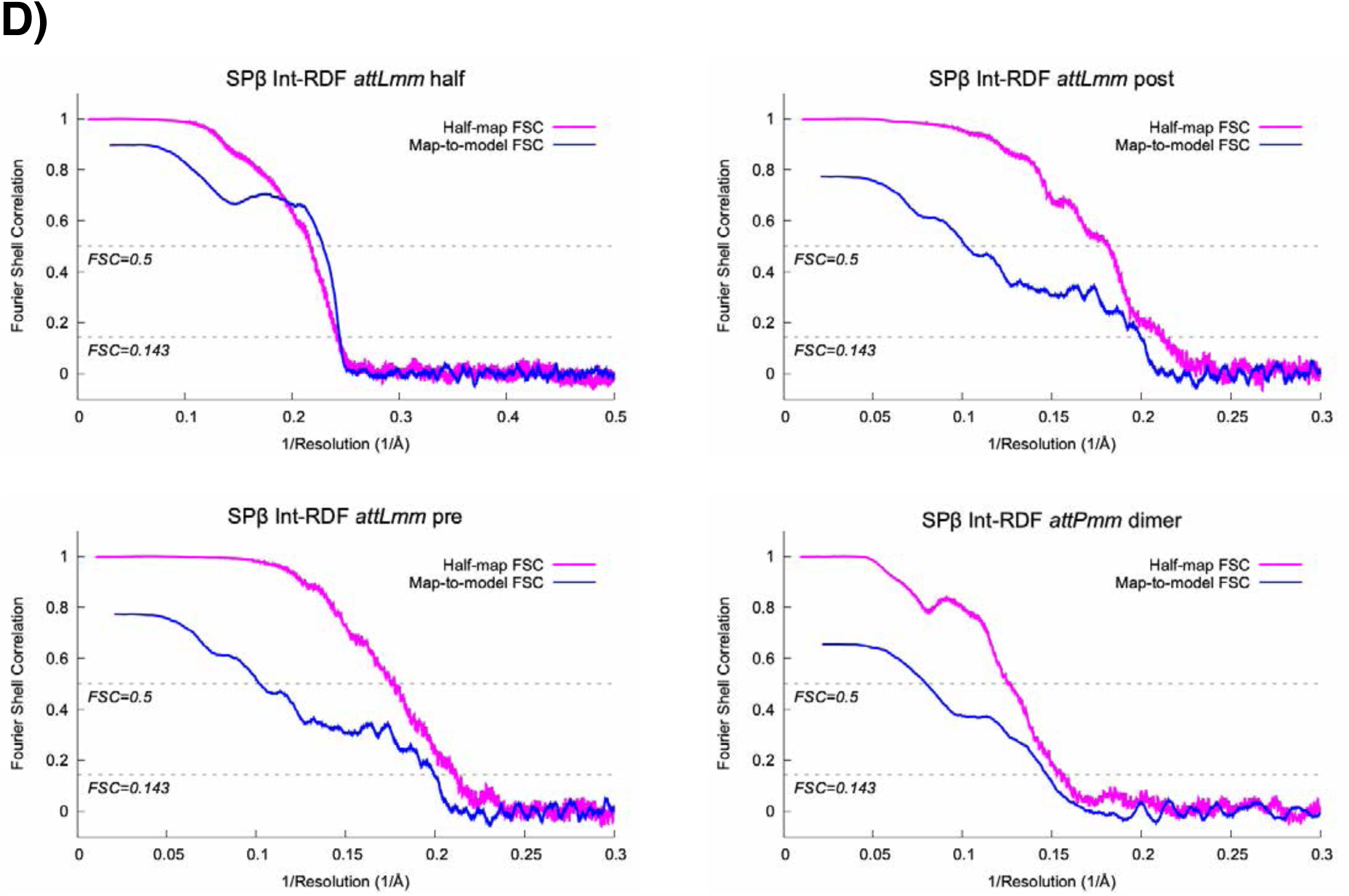
Data processing flow chart for the SPβ Int RDF *attLmm* data. A. A visual overview of the data processing steps for the SPβ Int RDF *attLmm* dataset. The dataset was subjected to particle selection, iterative rounds of 2D classification and 3D classification. A representative micrograph (scale bar, 40nm) and representative 2D class averages are shown. The distribution of Euler angles is displayed next to the half complex map. B. Local resolution of the map was calculated using cryoSPARC^74^. C. 3D Flexible refinement (3D Flex)^77^ was performed to model non-rigid or flexible regions. A custom mesh, consisting of six segmented regions, was generated to cover the half complex (left and middle), and a 3D Flex-refined map was obtained (right). D. Fourier shell correlation (FSC) curve of the masked map after cryoSPARC^74^ postprocessing. The resolution was determined by the FSC=0.143 (magenta). The map-to-model FSC curve is also shown (blue).

**Supplementary figure 12.**
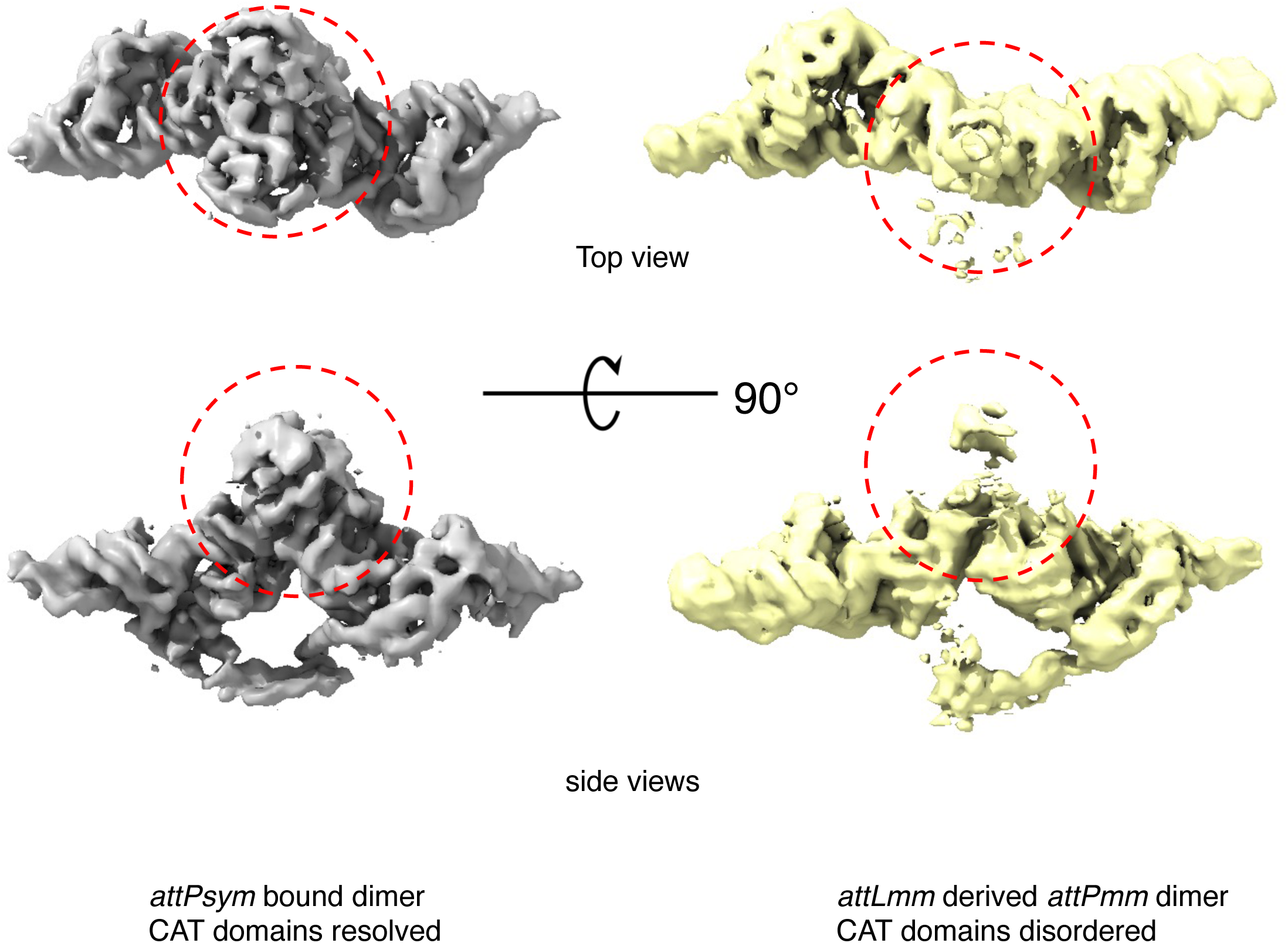
Cryo-EM maps of the *attPsym-* vs. *attLmm* derived *attPmm*-bound SPβ Integrase and RDF fusion dimeric complexes. Cryo-EM maps of the uncleaved *attPsym* bound SPβ integrase and RDF fusion dimeric complex (gray) and the cleaved *attPmm* bound SPβ integrase and RDF fusion dimeric complex (yellow). Regions corresponding to the CAT domains are circled in red.

**Supplementary figure 13.**
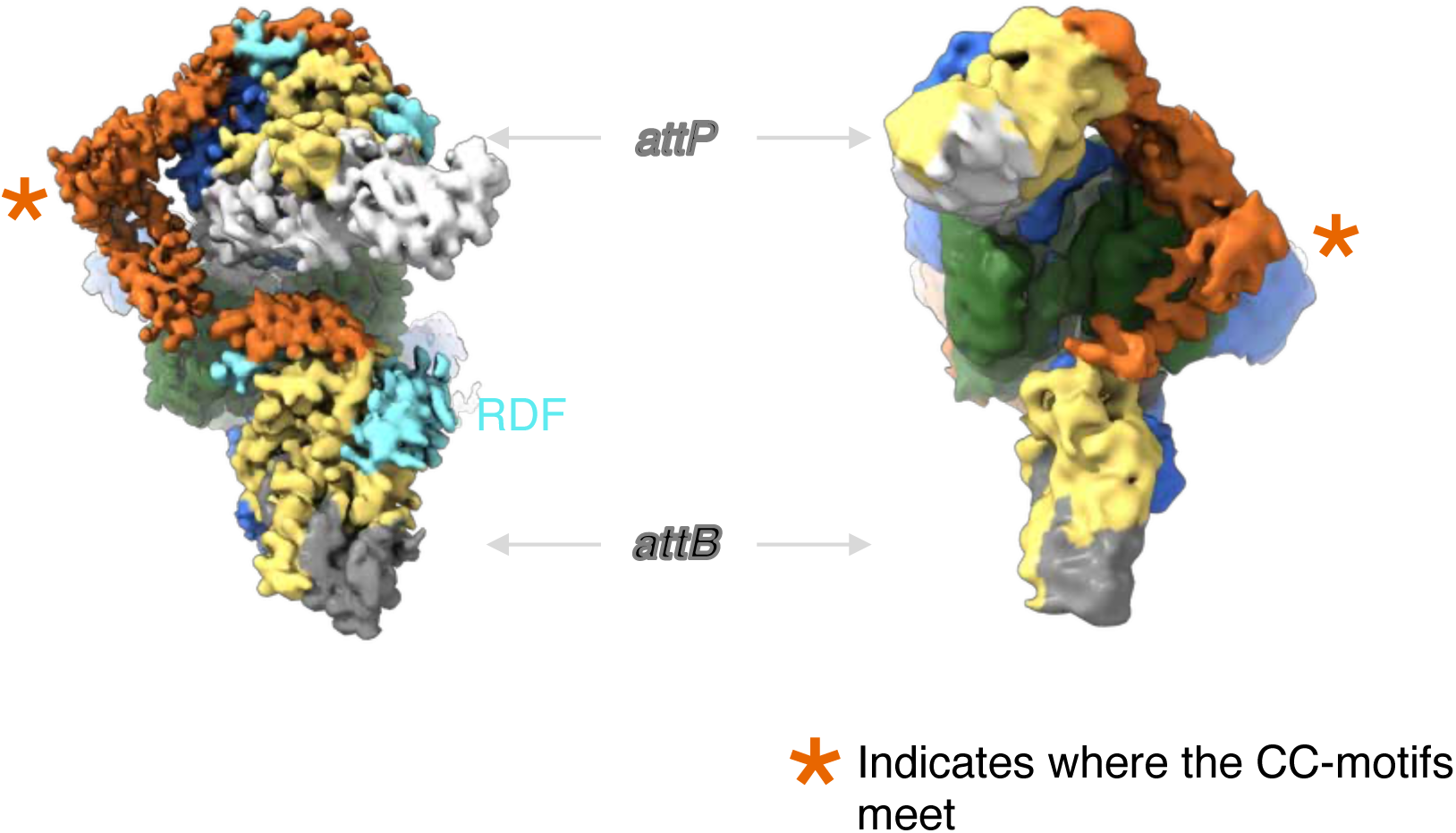
Redirection of the CC-motif trajectory by RDF confers distinct synaptic complexes. Side-by-side comparison of EM maps in the absence (right) and presence (left) of RDF (in cyan). In the presence of RDF binding, the CC-motifs are oriented in the opposite side of the synaptic complex. The location where the two CC-motifs meet is indicated by an orange asterisk.

**Supplementary figure 14.**
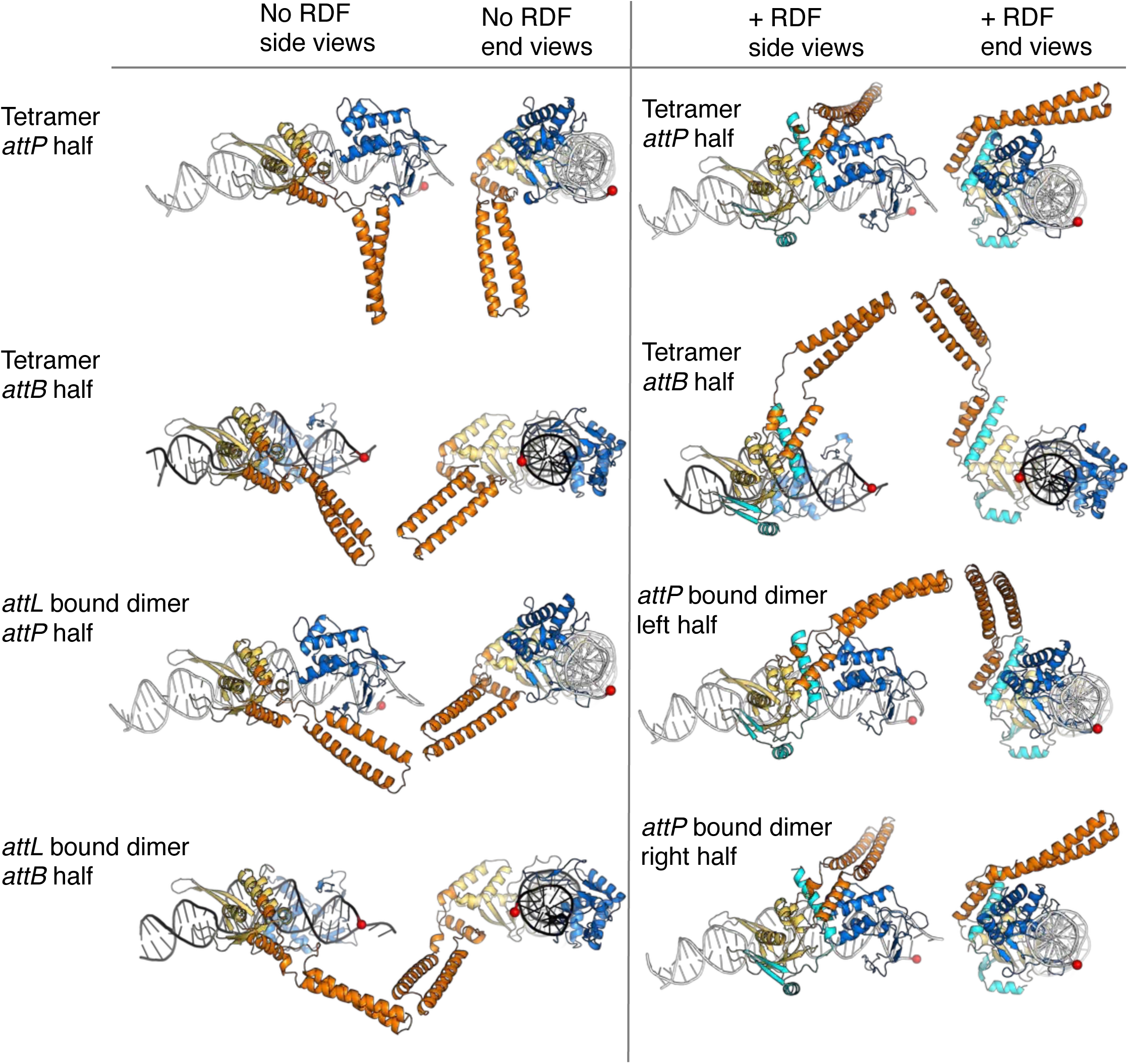
Comparison of all the DBD2-CC conformations in the presence and absence of RDF. For each conformation, two views are shown of the DNA half site and the subunit bound to it. All DBD2s are in the same orientation, and the CAT were domains removed for clarity.

**Supplementary figure 15.**
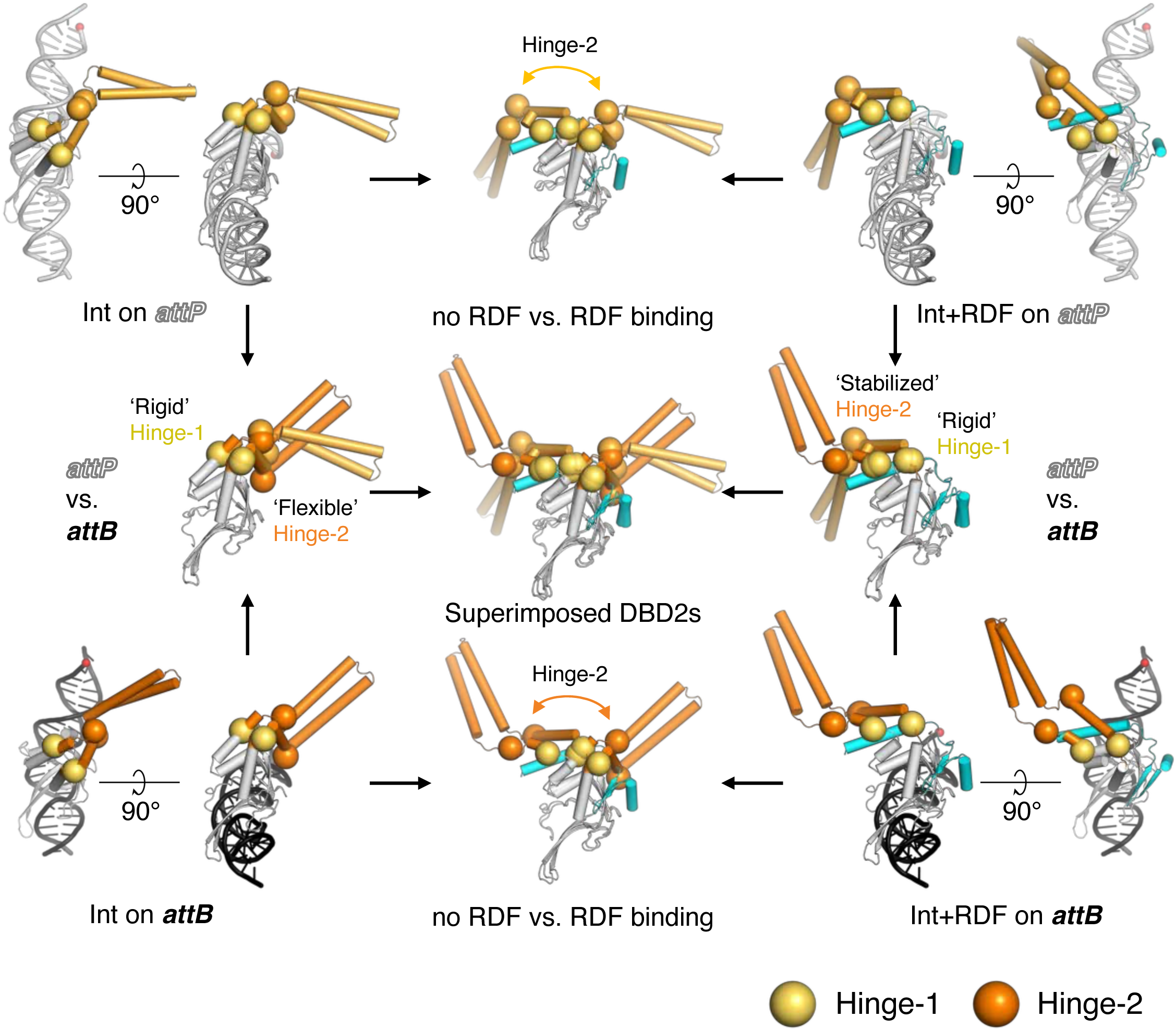
Structural details of Hinge-1 and Hinge-2 in DBD2. Comparison of CC motif projections in DBD2 domains bound to *attP* (top row) vs. *attB* (bottom row) substrates, in the absence (left column) and presence (right column) of RDF. Hinge-1 (residues 380 and 480) and Hinge-2 (residues 399 and 470) are indicated with spheres (light orange for *attB-*bound DBD2s and dark orange for *attP-*bound DBD2s). All DBD2 domains are aligned according to the DBD2 framework (aa 300-376 and 482-545), excluding the CC region (aa 377-481). Superimposed structures (visualized without DNA and shown in the middle row and column) are placed between the corresponding structures for comparison.

**Supplementary figure 16.**
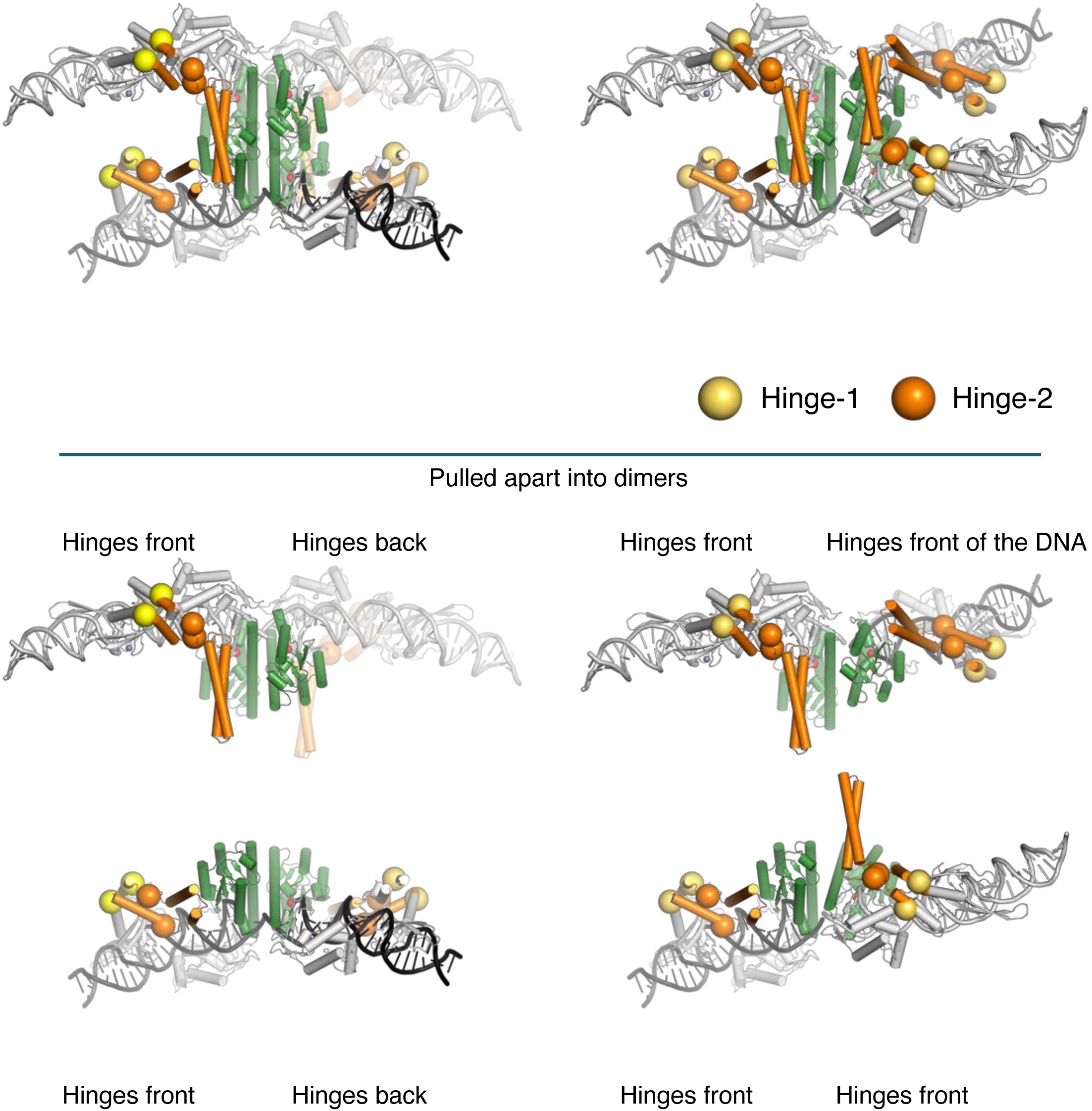
Positional changes of two hinges (or pivot points) in DBD2s during integration pathway. A. Comparison of SPβ Integrase synaptic complex in the pre-rotation (left) and post-rotation (right) states. Hinge-1s (aa 380 and 480) and Hinge-2s (aa 399 and 470) are indicated with yellow and orange spheres, respectively. B. The synaptic complexes are visualized as top and bottom dimers to simplify the view of hinge positioning and its influence on directionality control. Note that the Hinge-2s of the subunits that will form the product-bound dimers are closer together after rotation than before.

**Supplementary figure 17.**
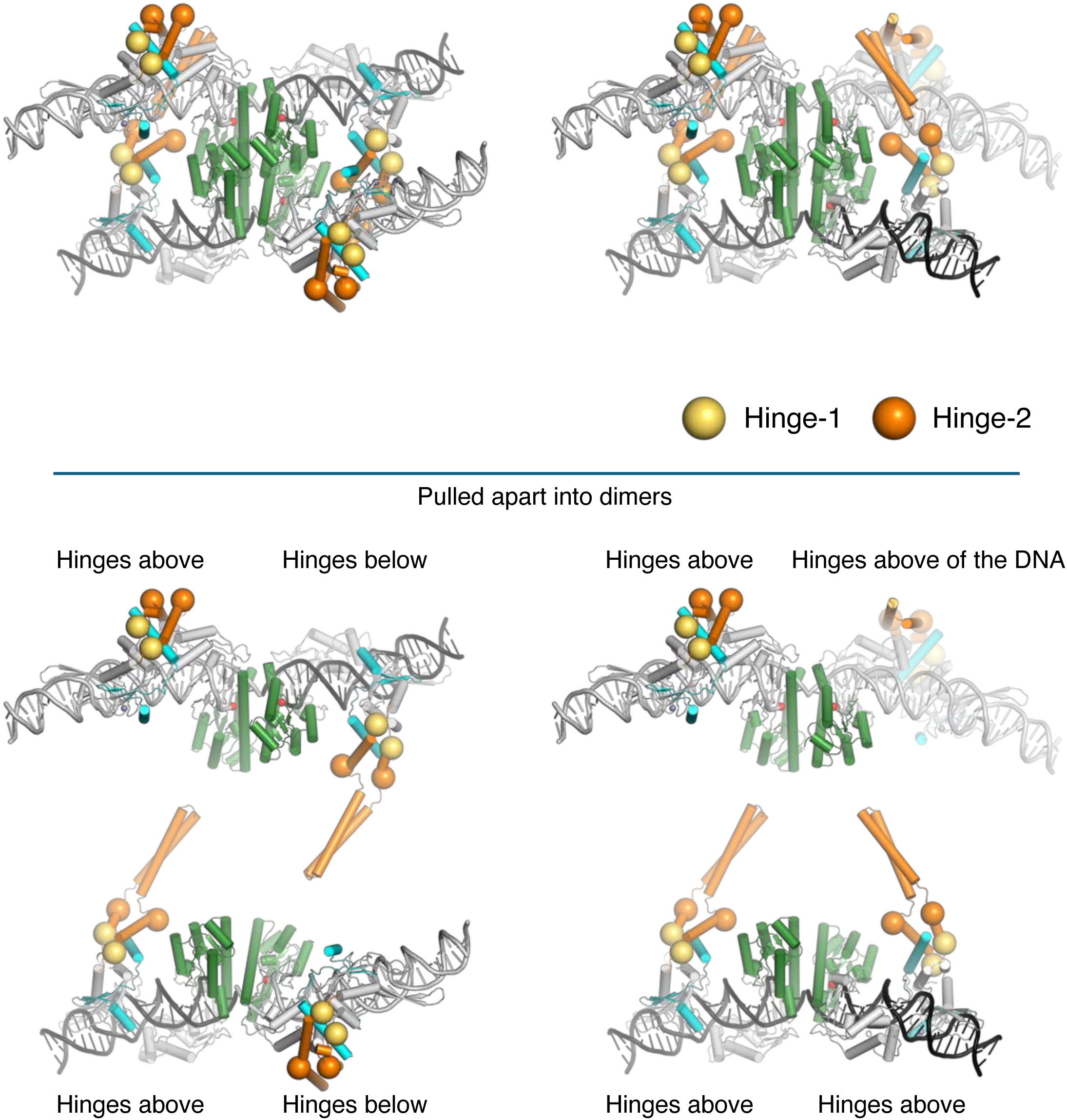
Positional changes of two hinges (or pivot points) in DBD2s during excision pathway. A. Comparison of SPβ Integrase-RDF synaptic complex in the pre-rotation (left) and post-rotation (right) states. Hinge-1s (aa 380 and 480) and Hinge-2s (aa 399 and 470) are indicated with yellow and orange spheres, respectively. B. The synaptic complexes are visualized as top and bottom dimers to simplify the view of hinge positioning and its influence on directionality control. Note that in both the integration and excision pathway, the Hinge-2s of the subunits that will form the product-bound dimers are closer together after rotation than before.

**Supplementary figure 18.**
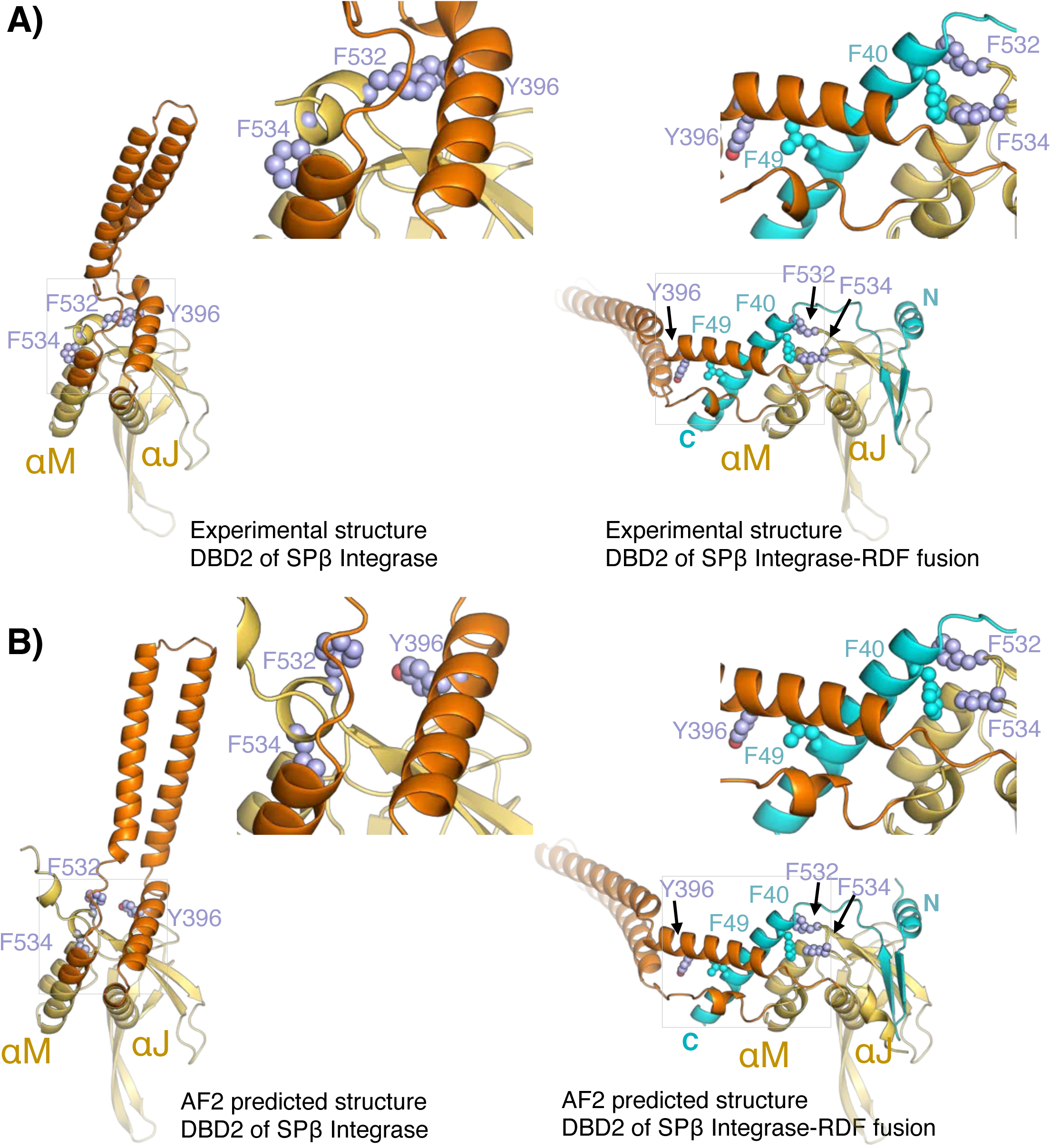
Repacking of the CC inter-hinge segment in the presence of RDF. A. Comparison of DBD2 domains in the absence (left) and presence (right) of RDF. Selected hydrophobic residues are shown in spheres (I389 and V477 from DBD2; L48 and L49 from RDF). In the absence of RDF, the C-terminal segment of αJ within the helical inter-hinge region packs against DBD2, effectively shielding the RDF binding region. Upon RDF binding, this helical segment unwinds. B. AlphaFold2^80^ predicted models illustrate the unwinding of this helical segment.

**Supplementary figure 19.**
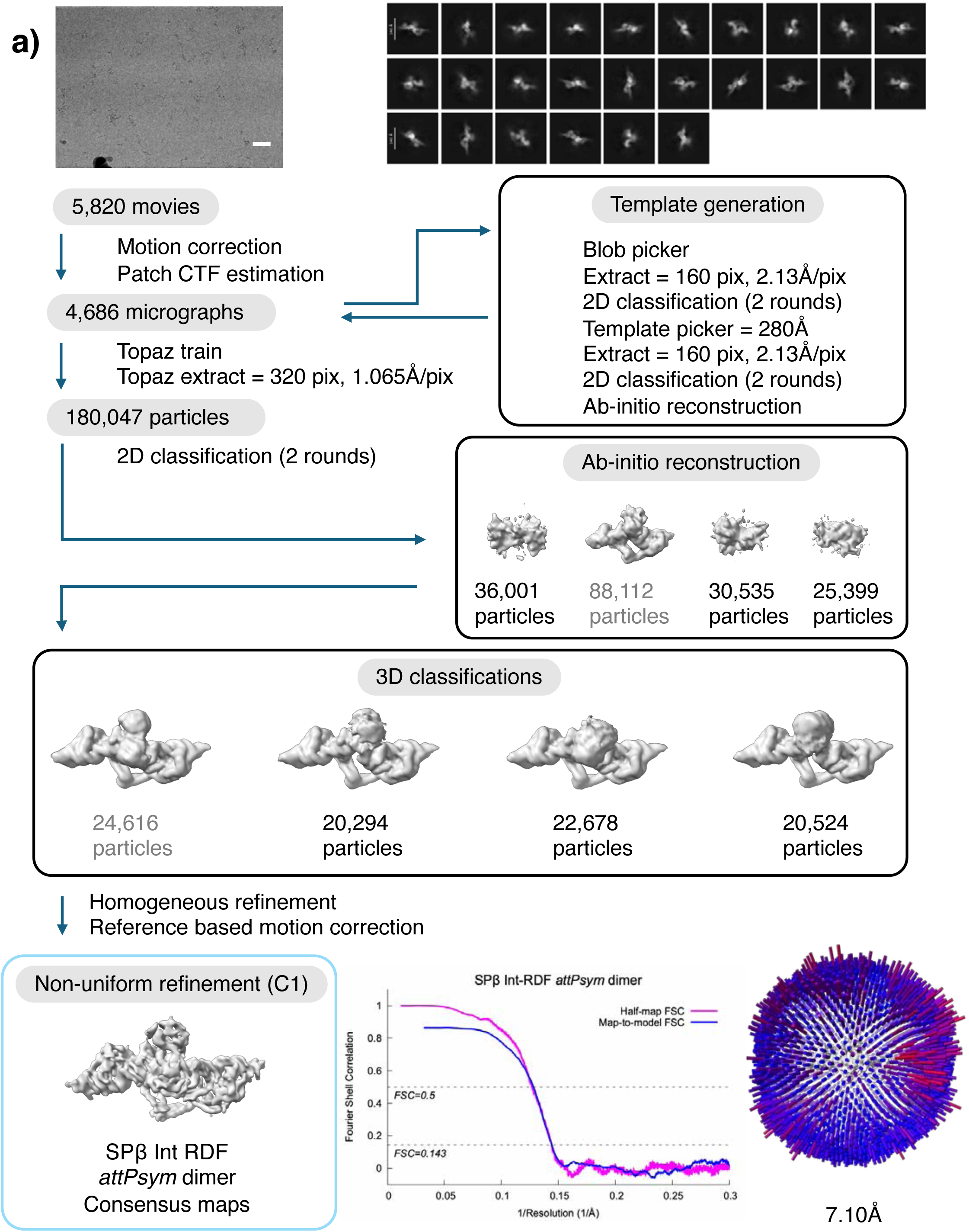
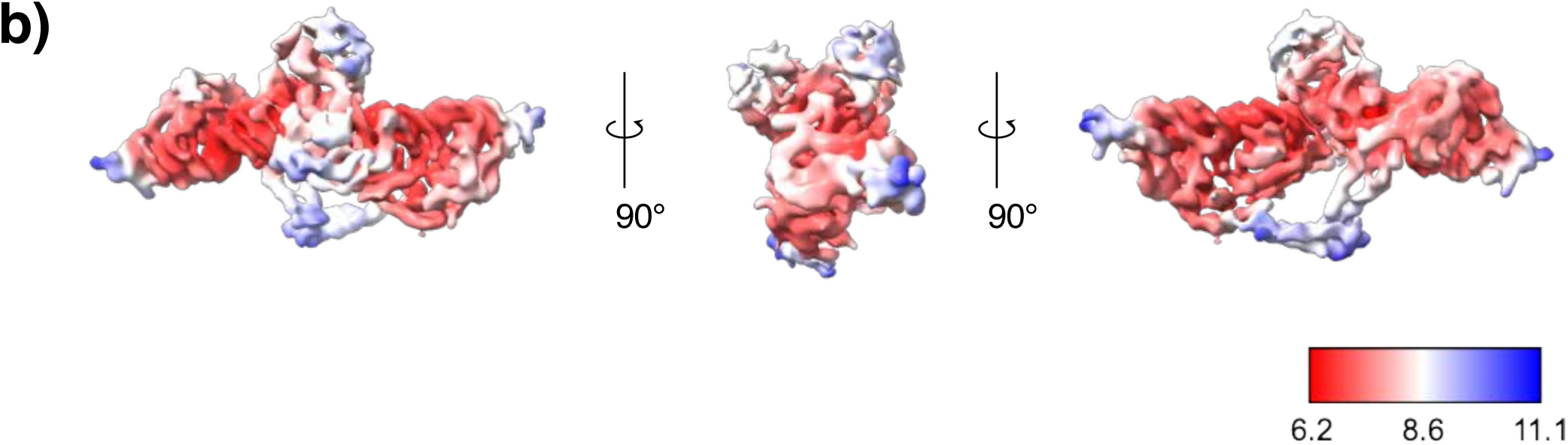
Data processing flow chart for the SPβ Int RDF *attPsym* data. A. A visual overview of the data processing steps for the SPβ Int RDF *attPsym* dataset. The dataset was subjected to particle selection, iterative rounds of 2D classification and 3D classification. A representative micrograph (scale bar, 40nm) and representative 2D class averages are shown. The distribution of the Euler angles is shown next to the consensus map. Fourier shell correlation (FSC) curve of the masked map after cryoSPARC^74^ postprocessing. The resolution was determined by the FSC=0.143 (magenta). The map-to-model FSC curve is also shown (blue). B. Local resolution of the map was calculated using cryoSPARC^74^.

**Supplementary figure 20.**
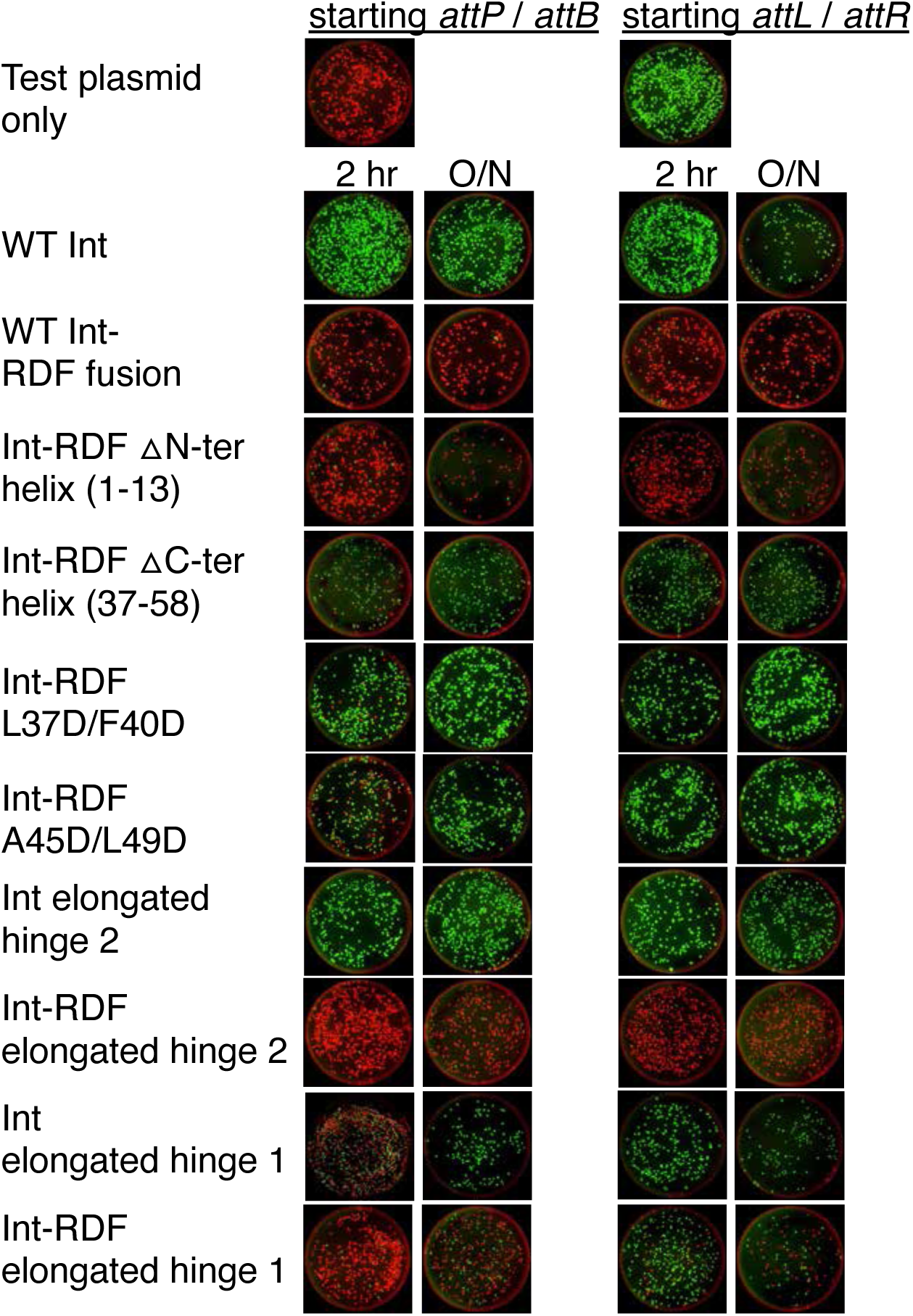
Mutational analysis of RDF and integrase hinge regions. Representative false-color images of plates of *E. coli* harboring the product plasmids from inversion assays showing red and green fluorescence overlaid. See Figure 7 for quantification. Int or Int-RDF fusions were expressed for 2 hours or overnight (see methods for details). ΔN-ter: deletion of the first helix of the RDF (Δ residues 1-13). ΔC-ter helix: RDF Δ residues 37-58. Elongated hinge 1: addition of GGSGSSG between D379 & L380 and between N479 & N480; Elongated hinge 2 : addition of GGSGSG between S401 & N402 and between D467 & T468.

**Supplementary figure 21.**
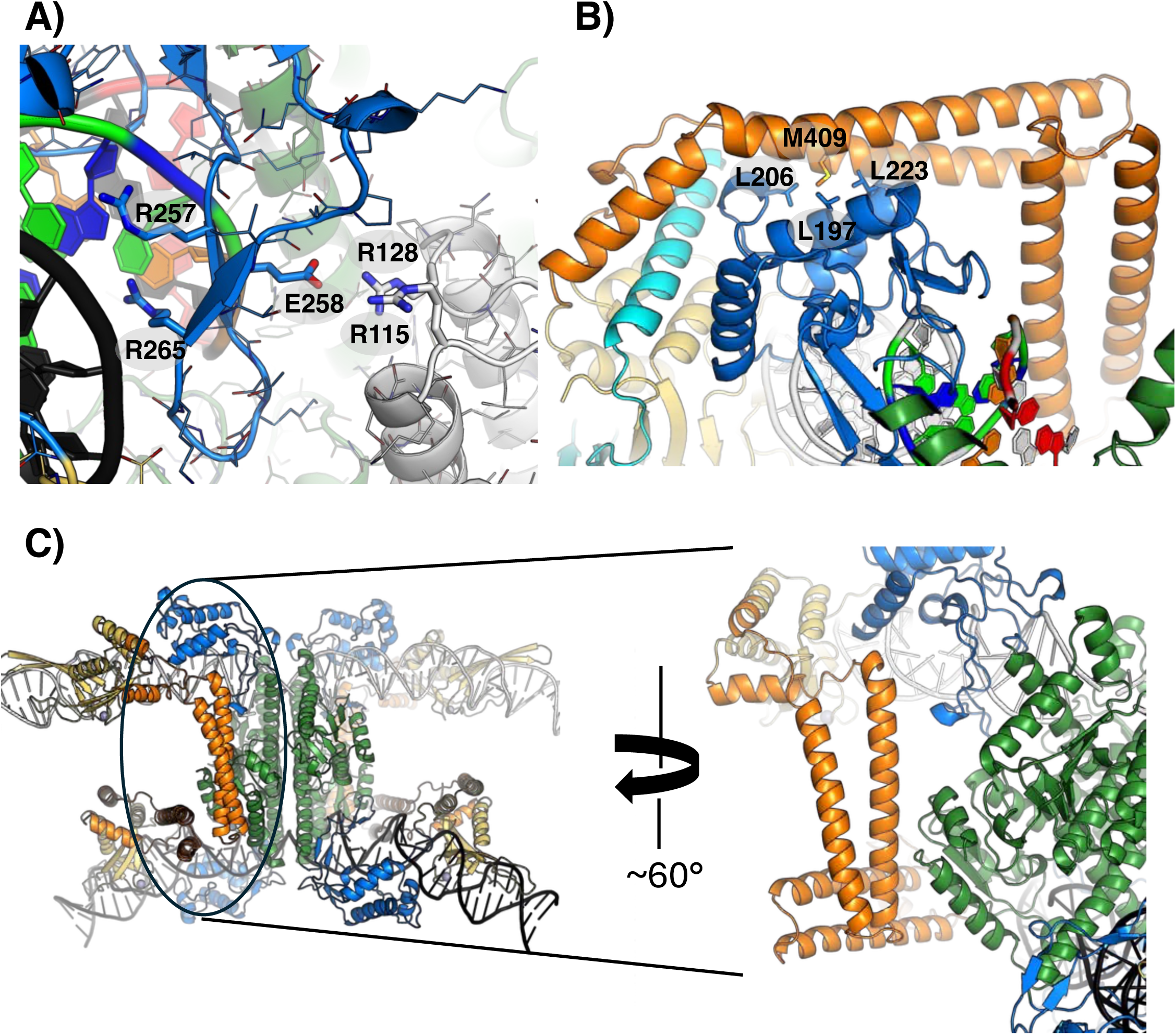
Additional protein—protein interactions. A. The DNA-interacting loop of DBD1 (blue) interacts with the cat domain (in gray). Similar interactions were noted in both in the presence and absence of the RDF and on both types of half-sites. B. In the presence of the RDF, the *attP* – bound DBD1 interacts with the CC motif (orange) and RDF (cyan). These interactions were not seen on *attB* sites or without the RDF. C. In the absence of the RDF, the CC motifs interact with the cat domains (green).

**Supplementary figure 22.**
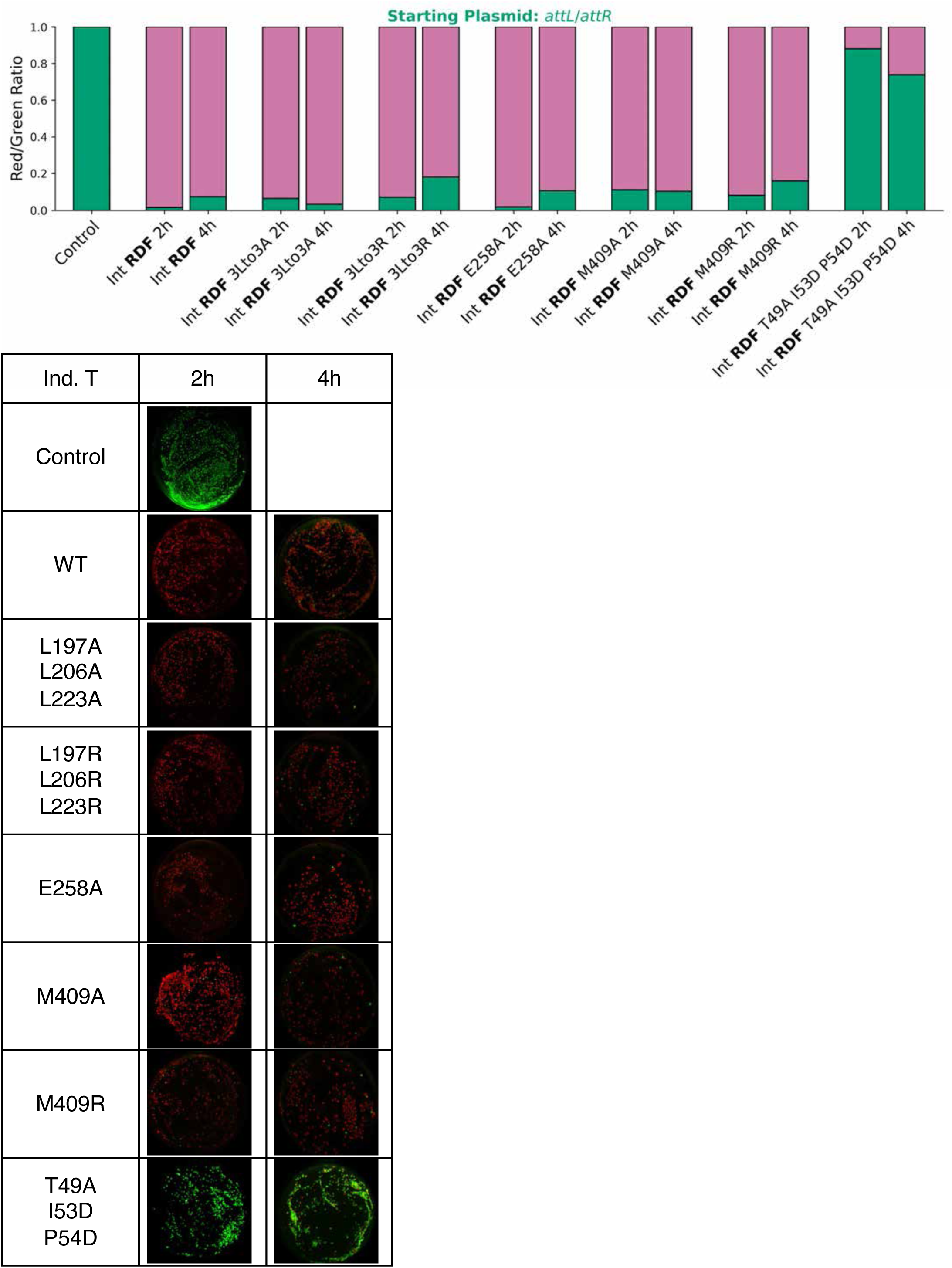
Mutational analysis of SPβ integrase-RDF fusion on additional protein interactions. *in vivo* inversion assays with *attL* x *attR* starting plasmid. SPβ Int-RDF fusion was induced for 2 hours and 4 hours

**Supplementary figure 23.**
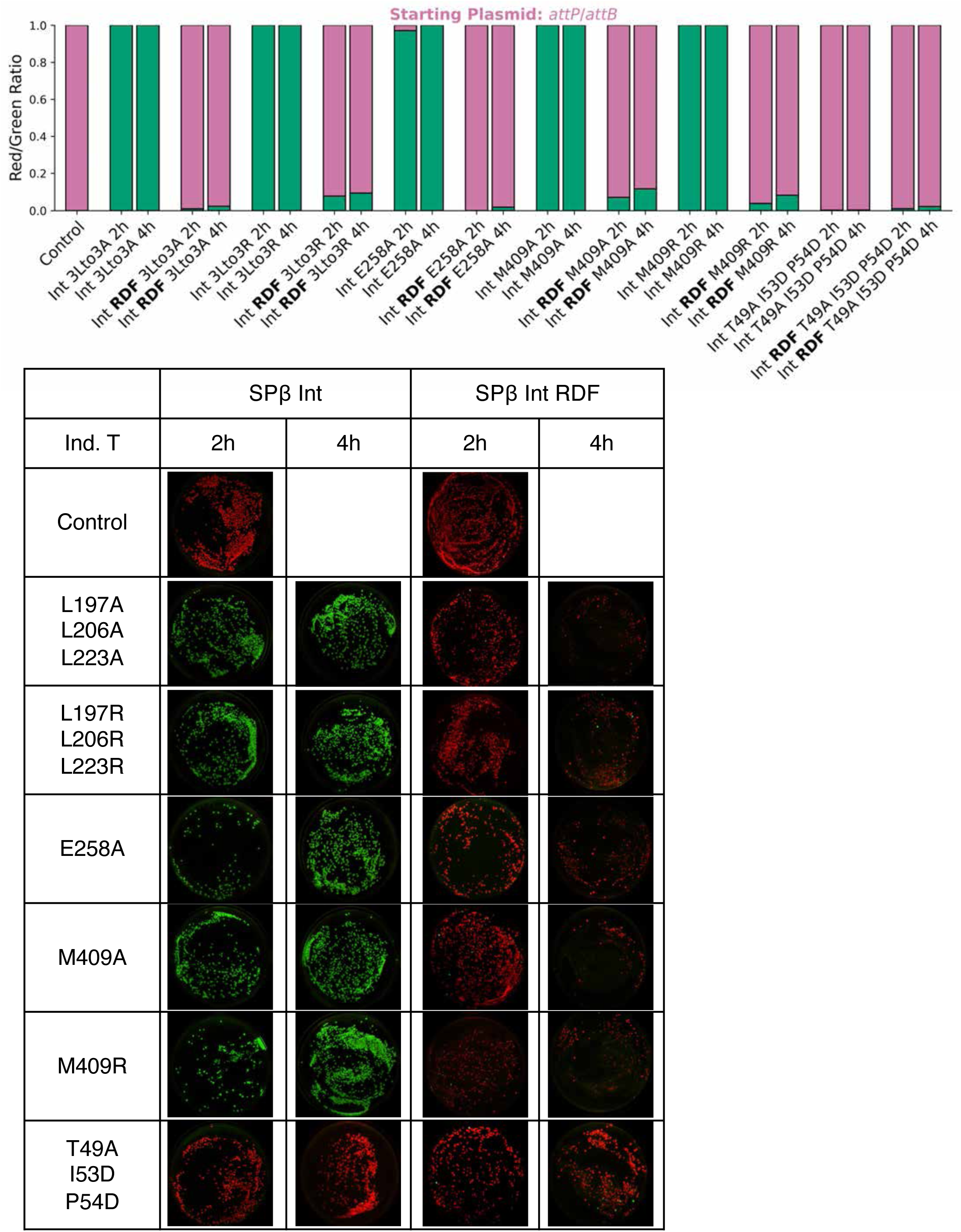
Mutational analysis of SPβ integrase on additional protein interactions. *in vivo* inversion assays with *attP* x *attB* starting plasmid. SPβ Int and Int-RDF fusions were induced for 2 hours and 4 hours

**Supplementary figure 24.**
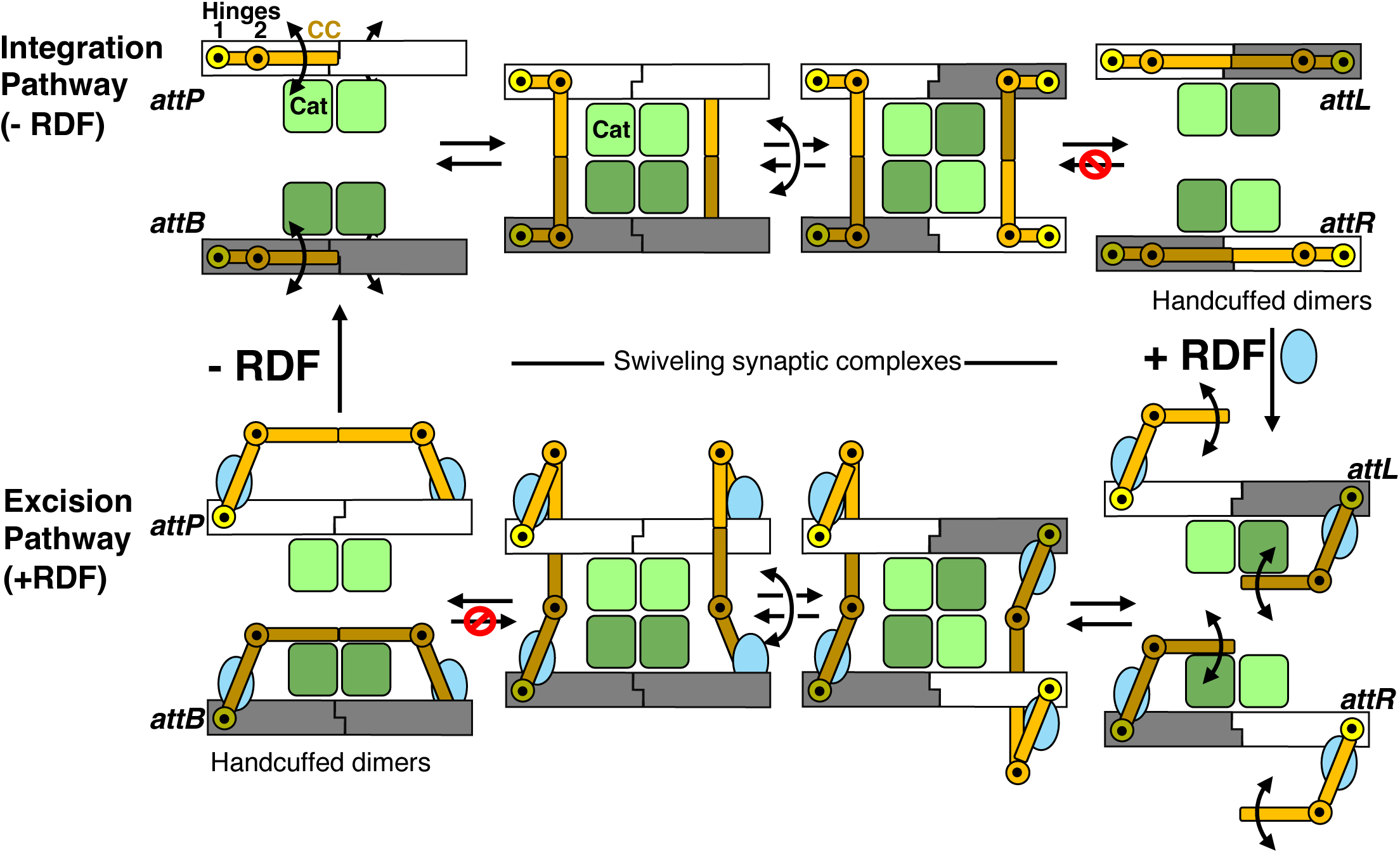
Simplified cartoon of reaction pathways. Only the catalytic domain (green), CC (orange), the hinges (black dots in yellow and orange circles), the RDF (blue), and the DNA (*attP* in white and *attB* in gray) are shown. DNA binding domains not shown for simplicity. Both Hinge 1 and Hinge 2 are critical to directionality. The location of Hinge 1 depends only on whether the protein is bound to an *attP*- or *attB*-derived half-site, and the location of Hinge 2 depends on the presence or absence of the RDF. Within each pathway, the pivot point for the CC is hinge 2.

